# Definition of a saxitoxin (STX) binding code enables discovery and characterization of the Anuran saxiphilin family

**DOI:** 10.1101/2022.06.09.495489

**Authors:** Zhou Chen, Sandra Zakrzewska, Holly S. Hajare, Aurora Alvarez-Buylla, Fayal Abderemane-Ali, Maximiliana Bogan, Dave Ramirez, Lauren A. O’Connell, J. Du Bois, Daniel L. Minor

**Author notes:** Equal contributions (Zhou Chen, Sandra Zakrzewska).

## Abstract

American bullfrog (Rana castesbeiana) saxiphilin (RcSxph) is a high-affinity ‘toxin sponge’ protein thought to prevent intoxication by saxitoxin (STX), a lethal bis-guanidinium neurotoxin that causes paralytic shellfish poisoning (PSP) by blocking voltage-gated sodium channels (NaVs). How specific RcSxph interactions contribute to STX binding has not been defined and whether other organisms have similar proteins is unclear. Here, we use mutagenesis, ligand binding, and structural studies to define the energetic basis of Sxph:STX recognition. The resultant STX ‘recognition code’ enabled engineering of RcSxph to improve its ability to rescue NaVs from STX and facilitated discovery of ten new frog and toad Sxphs. Definition of the STX binding code and Sxph family expansion among diverse Anurans separated by ∼140 million years of evolution provides a molecular basis for understanding the roles of toxin sponge proteins in toxin resistance and for developing novel proteins to sense or neutralize STX and related PSP toxins.

**Teaser:** A conserved STX recognition motif from frog and toad saxiphilins defines molecular principles of paralytic toxin binding.

## Introduction

Saxitoxin (STX), one of the most lethal non-peptidyl neurotoxins, blocks the bioelectrical signals in nerve and muscle required for life by inhibiting select voltage-gated sodium channel (Na_V_) isoforms (*1–3*). Cyanobacteria and dinoflagellate species associated with oceanic red tides produce this bis-guanidinium small molecule and its congeners whose accumulation in seafood can cause paralytic shellfish poisoning (PSP), a commer*c*ial fishing and public health hazard of growing importance due to climate change (*1, 3–5*). Its extreme lethality has also earned STX the unusual distinction of being the only marine toxin declared a chemical weapon (*1, 3*). Select vertebrates, particularly frogs, resist STX poisoning (*6–9*), a property that is thought to rely on the ability of the soluble ‘toxin sponge’ protein saxiphilin (Sxph) to sequester STX (*8, 9*). Recent structural studies (*10*) defined the molecular architecture of the American bullfrog (*Rana (Lithobates) castesbeiana*) Sxph (*Rc*Sxph) (*8, 11–14*) showing that this 91 kDa soluble, transferrin-related protein from frog heart and plasma has a single, high-affinity STX binding site on its C-lobe. Remarkably, even though *Rc*Sxph and Na_V_s are unrelated, both engage STX through similar types of interactions (*10*). This structural convergence raises the possibility that determination of the factors that underlie the high-affinity Sxph:STX interaction could provide a generalizable molecular recognition code for STX that would enable the identification or engineering of STX binding sites in natural and designed proteins.

To characterize *Rc*Sxph:STX interactions in detail, we developed a suite of assays comprising thermofluor (TF) measurements of ligand-induced changes in *Rc*Sxph stability, fluorescence polarization (FP) binding to a fluorescein-labeled STX, and isothermal titration calorimetry (ITC). We paired these assays with a scanning mutagenesis strategy (*15, 16*) to dissect the energetic contributions of *Rc*Sxph STX binding pocket residues. These studies show that the core *Rc*Sxph STX recognition code comprises two ‘hot spot’ triads. One engages the STX tricyclic bis-guanidinium core through a pair of carboxylate groups and a cation-π interaction (*17*) in a manner that underscores the convergent STX recognition strategies shared by *Rc*Sxph and Na_V_s (*17–22*). The second triad largely interacts with the C13 carbamate group of STX and is the site of interactions that can enhance STX binding affinity and the ability of *Rc*Sxph to act as a ‘toxin sponge’ that can reverse the effects of STX inhibition of Na_V_s (*8, 9*).

Although Sxph-like STX binding activity has been reported in extracts from diverse organisms including arthropods (*13*), amphibians (*11, 13, 23*), fish (*13*), and reptiles (*13*), the molecular origins of this activity have remained obscure. Definition of the *Rc*Sxph STX recognition code enabled identification of ten new Sxphs from diverse frogs and toads. This substantial enlargement of the Sxph family beyond *Rc*Sxph and the previously identified High Himalaya frog (*Nanorana parkeri*) Sxph (*Np*Sxph) (*10*) reveals a varied STX binding pocket that surrounds a conserved core of ‘hot spot’ positions. Comparison of the new Sxph family members further identifies dramatic differences in the number of thyroglobulin (Thy1) domains inserted into the modified transferrin fold upon which the Sxph family is built. Biochemical characterization of *Np*Sxph, *Oophaga sylvatica* Sxph (*Os*Sxph) (*24*), *Mantella aurantica* Sxph (*Ma*Sxph), and *Ranitomeya imitator* Sxph (*Ri*Sxph), together with structural determination of *Np*Sxph, alone and as STX complexes, shows that the different Sxphs share the capacity to form high affinity STX complexes and that binding site preorganization (*10*) is a critical factor for tight STX association. Together, these studies establish a STX molecular recognition code that provides a template for understanding how diverse STX binding proteins engage the toxin and its congeners and uncover that Sxph family members are abundantly found in the most varied and widespread group of amphibians, the Anurans. This knowledge and suite of diverse Sxphs, conserved among Anuran families separated by a least 140 million years of evolution (*25*), provides a starting point for defining the physiological roles of Sxph in toxin resistance (*9, 24, 26*), should facilitate identification or design of other STX binding proteins, and may enable the development of new biologics to detect or neutralize STX and related PSPs.

## Results

### Establishment of a suite of assays to probe *Rc*Sxph toxin binding properties

To investigate the molecular details of the high-affinity *Rc*Sxph:STX interaction, we developed three assays to assess the effects of STX binding site mutations. A key criterion was to create assays that could be performed in parallel on many *Rc*Sxph mutants using minimal amounts of purified protein and toxin. To this end, we first tested whether we could detect STX binding using a thermofluor (TF) assay (*27, 28*) in which STX binding would manifest as concentration-dependent change in the apparent *Rc*Sxph melting temperature (Tm). Addition of STX, but not the related guanidinium toxin, tetrodotoxin (TTX), over a 0-20 µM range to samples containing 1.1 µM *Rc*Sxph caused concentration-dependent shifts in the *Rc*Sxph melting cure (Fig. 1A)(ΔTm = 3.6°C ± 0.2 versus 0.3°C ± 0.4 for STX and TTX, respectively). These differential effects of STX and TTX are in line with the ability of *Rc*Sxph to bind STX (*8, 13, 14*) but not TTX (*8, 9*) and indicate that ΔTm is a consequence of the *Rc*Sxph:STX interaction.

**Figure 1.**
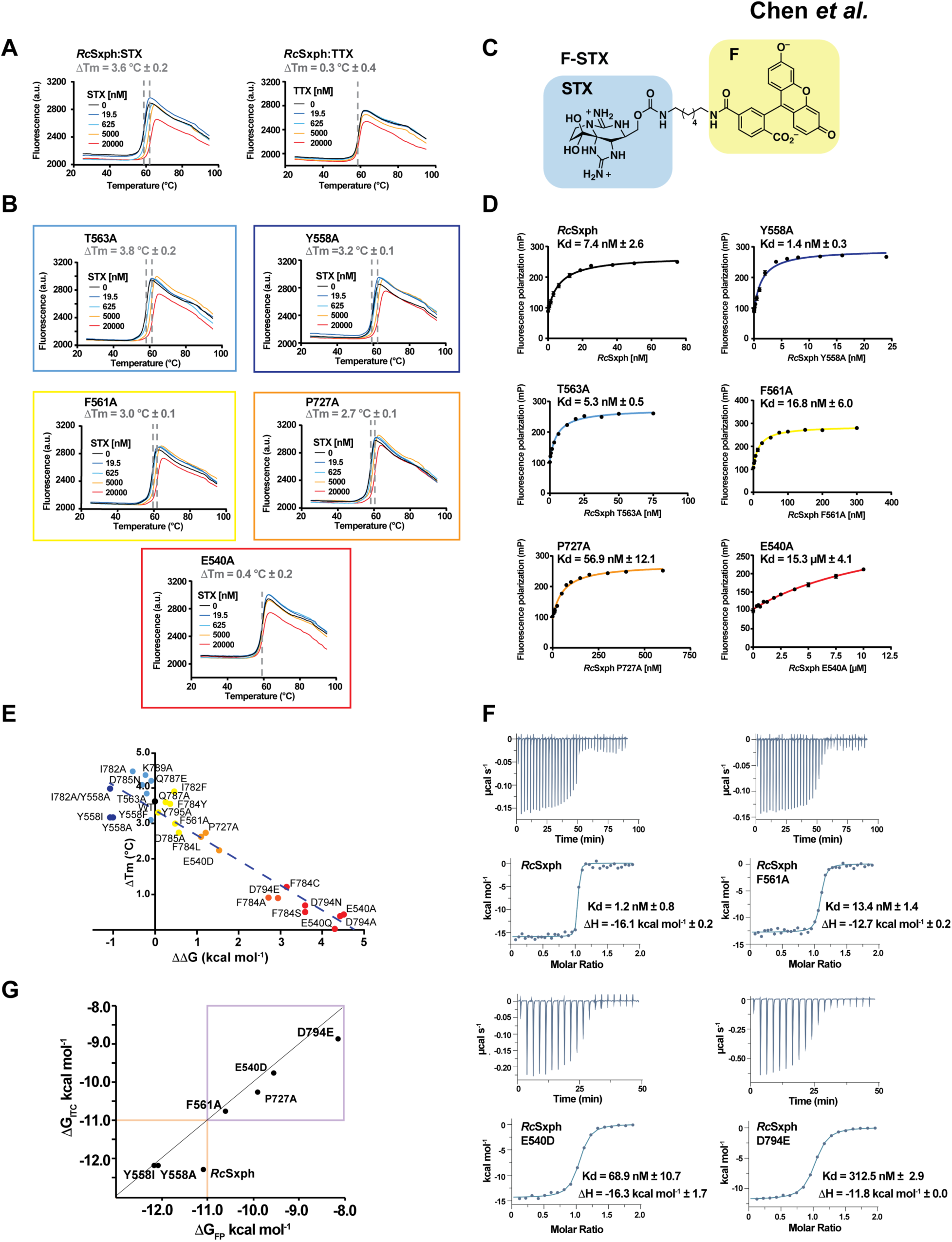
Alanine scan of *Rc*Sxph binding. **A**, and **B,** Exemplar thermofluor (TF) assay results for **A,** *Rc*Sxph in the presence of the indicated concentrations of STX (left) and TTX (right) and **B**, Select *Rc*Sxph mutants in the presence of STX. STX and TTX concentrations are 0 nM (black), 19.5 nM (blue), 625 nM (cyan), 5000 nM (orange), and 20000 nM (red). Grey dashed lines indicate ΔTm. **C,** F-STX diagram. STX and fluorescein (F) moieties are highlighted blue and yellow, respectively. **D,** Exemplar Fluorescence polarization (FP) binding curves and Kds for *Rc*Sxph and the indicated mutants. **E,** Comparison of *Rc*Sxph mutant ΔTm and ΔΔG values (line y = 3.49 −0.7523x, R^2^=0.886). **F,** Exemplar isotherms for titration of 100 µM STX into 10 µM *Rc*Sxph. 100 µM STX into 10 µM *Rc*Sxph F561A, 100 µM STX into 10 µM *Rc*Sxph E540D, and 300 µM STX into 30 µM *Rc*Sxph D794E. Kd and ΔH values are indicated. **G,** Comparison of ΔG_ITC_ for STX and ΔG_FP_ for F-STX for *Rc*Sxph and mutants. Purple box highlights region of good correlation. Orange box indicates region outside of the ITC dynamic range. (line shows x=y) Colors in ‘**B**’, ‘**D**’, and ‘**E**’ correspond to classifications in Table 1.

To investigate the contributions of residues that comprise the STX binding site, we coupled the TF assay with alanine scanning (*15*), as well as deeper mutagenesis studies, targeting the eight residues that directly contact STX (Glu540, Phe561, Thr563, Tyr558, Pro727, Phe784, Asp785, Asp794) (*10*) and four second shell sites that support these residues (Tyr795, Ile782, Gln787, and Lys789) (Figs. 1A-B and S1A). Measurement of the STX-induced ΔTm changes for the purified *Rc*Sxph mutants revealed ΔTm changes spread over a ∼4°C range that included ΔTm increases relative to wild-type (e.g. I782A and D785N) as well as those that caused complete loss of the thermal shift (e.g. E540A and D794A). All mutations had minimal effects on protein stability (Fig. S1B) and there was no evident correlation between Tm and ΔTm (Fig. S1C). Hence, the varied ΔTms indicate that each of the twelve positions contribute differently to STX binding.

Because ΔTm interpretation can be complex, especially in the case of a multidomain protein such as *Rc*Sxph, and may not necessarily indicate changes in ligand affinity (*27, 28*), we developed a second assay to measure the effects of mutations on *Rc*Sxph binding affinity. We synthesized a fluorescein-labeled STX derivative (F-STX) by functionalization of the pendant carbamate group with a 6-carbon linker and fluorescein (*29, 30*) (Figs. 1C and S2) and established a fluorescence polarization (FP) assay (*31, 32*) to measure toxin binding. FP measurements revealed a high-affinity interaction between F-STX and *Rc*Sxph (Kd = 7.4 nM ± 2.6) that closely agrees with prior radioligand assay measurements of *Rc*Sxph affinity for STX (∼1 nM) (*14*). The similarity between the F-STX and STX Kd values is consistent with the expectation from the *Rc*Sxph:STX structure that STX carbamate derivatization should have a minimal effect on binding, as this element resides on the solvent exposed side of the STX binding pocket (*10*). To investigate the F-STX interaction further, we soaked *Rc*Sxph crystals with F-STX and determined the structure of the *Rc*Sxph:F-STX complex at 2.65Å resolution by X-ray crystallography (Fig. 3A, Table S1). Inspection of the STX binding pocket revealed clear electron density for the F-STX bis-guanidinium core as well as weaker density that we could assign to the fluorescein heterocycle (Fig. S3A), although the high B-factors of the linker and fluorescein indicate that these moieties are highly mobile (Fig. S3B). Structural comparison with the *Rc*Sxph:STX complex (*10*) showed no changes in the core STX binding pose or STX binding pocket residues (RMSD_Cα_ = 0.279Å) (Fig. S3C). Together, these data demonstrate that both F-STX and STX bind to Sxph in the same manner and indicate that there are no substantial interactions with the fluorescein label.

FP measurement of the *Rc*Sxph alanine scan mutants uncovered binding affinity changes spanning a ∼13,000 fold range that correspond to free energy perturbations (ΔΔG) of up to ∼5.60 kcal mol^-1^ (Figs. 1D and S4, Table 1). The effects were diverse, encompassing enhanced affinity changes (Y558A Kd = 1.2 nM ± 0.3) and large disruptions (E540A Kd = 15.3 µM ± 4.1). As indicated by the TF data, each STX binding pocket residue contributes differently to STX recognition energetics. Comparison of the TF ΔTm and FP ΔΔG values shows a strong correlation between the two measurements (Fig.1E). This concordance between ΔTm and ΔΔG indicates that the changes in unfolding free energies caused by protein mutation and changes in STX binding affinity do not incur large heat capacity or entropy changes relative to the wild-type protein (*33, 34*). Hence, ΔTm values provide an accurate estimate of the STX binding affinity differences.

**Table 1.**
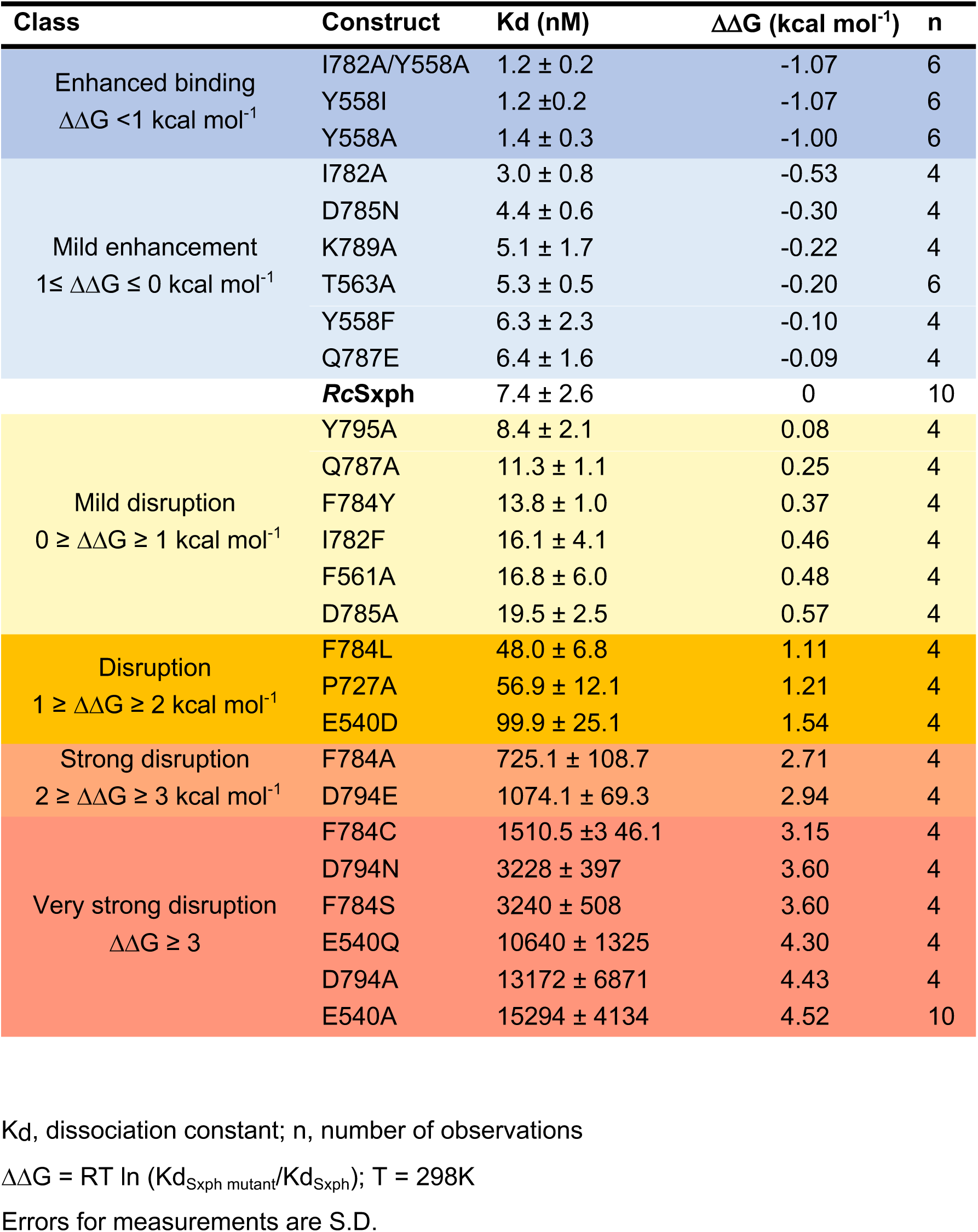
*Rc*Sxph STX binding pocket mutant binding parameters.

To investigate the STX affinity changes further, we used isothermal titration calorimetry (ITC) (Figs. 1F, and Table S2), a label-free method that reports directly on ligand association energetics (*35*), to examine the interaction of STX with *Rc*Sxph and six mutants having varied effects on binding (E540D, Y558I, Y558A, F561A, P727A, and D794E) (Figs. 1F, 2A, S5A-C, and Table S2). Experiments with *Rc*Sxph confirm the 1:1 stoichiometry and high affinity of the *Rc*Sxph:STX interaction (Kd ∼nM) reported previously (*8, 10, 14*) and reveal a large, favorable binding enthalpy (ΔH −16.1 ± 0.2 kcal mol^-1^) in line with previous radioligand binding studies (*14*). In almost all mutants, binding affinity loss correlated with a reduction in enthalpy, consistent with a loss of interactions (Table S2). The one exception to this trend is E540D for which STX association yielded a binding enthalpy (ΔH −16.3 ± 1.7 kcal mol^-1^) very similar to wild type *Rc*Sxph that was offset by a ∼two-fold unfavorable change in binding entropy. The ITC measurements were unable to measure the affinity enhancement for Y558A and Y558I accurately due to the fact that these mutants, as well as *Rc*Sxph have Kds at the detection limit of direct titration methods(∼1 nM) (*35*). Nevertheless, ΔG_ITC_ from mutants having STX Kds within the ITC dynamic range (Kds ∼30-300 nM) showed an excellent agreement with ΔG_FP_ measurements made with F-STX (Fig. 1G). These data further validate the TF and FP assay trends and support the conclusion that *Rc*Spxh:F-STX binding interactions are very similar to the *Rc*Spxh:STX interactions. Together, these three assays (Figs. 1E and G) provide a robust and versatile suite of options for characterizing STX:Sxph interactions.

**Figure 2.**
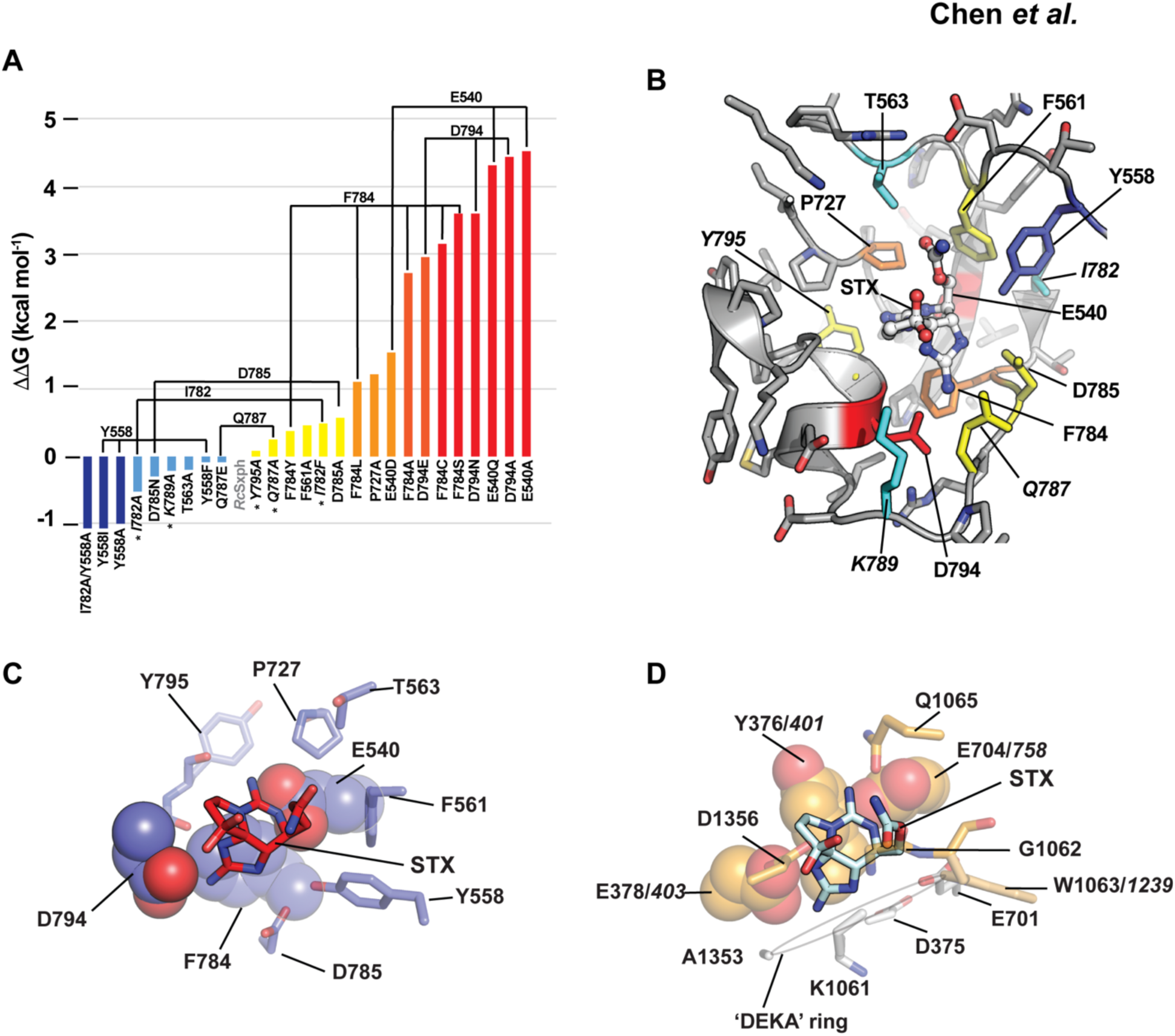
Energetic fingerprint of STX recognition by *Rc*Sxph. **A**, ΔΔG comparisons for the indicated *Rc*Sxph STX binding pocket mutants relative to wild-type *Rc*Sxph. Colors indicate:ΔΔG <-1 kcal mol^-1^ (blue), −1≤ ΔΔG ≤ 0 kcal mol^-1^ (light blue), 0 ≥ ΔΔG ≥ 1 kcal mol^-1^ (yellow), 1 ≥ ΔΔG ≥ 2 kcal mol^-1^ (orange), 2 ≥ ΔΔG ≥ 3 kcal mol^-1^ (red orange), and ΔΔG ≥ 3 (red). **B,** Energetic map of alanine scan mutations on STX binding to the *Rc*Sxph STX binding pocket (PDB:6O0F) (*10*). Second shell sites are in italics. Colors are as in ‘**A**’. **C** and **D**, Structural interactions of STX with **C,** *Rc*Sxph (PDB:6O0F) (*10*) and **D,** human Na_V_1.7 (PDB:6J8G) (*40*). Residues that are energetically important for the STX interaction are shown in space filling. Na_V_1.7 selectivity filter ‘DEKA’ ring residues are shown (white). Italics indicate corresponding residue numbers for rat Na_V_1.4 (*18*).

### Sxph STX binding code is focused on two sets of ‘hot spot’ residues

To understand the structural code underlying STX binding, we classified the effects of the alanine mutations into six groups based on ΔΔG values (Fig. 2A and Table 1) and mapped these onto the *Rc*Sxph structure (Fig. 2B). This analysis identified a binding ‘hot spot’ comprising three residues that directly contact the STX bis-guanidinium core (Glu540, Phe784, and Asp794) (*10*) and an additional site near the carbamate (Pro727) where alanine mutations caused substantial STX binding losses (ΔΔG ≥1 kcal mol^-1^). Conversely, we also identified a site (Tyr558) where alanine caused a notable enhancement of STX binding (ΔΔG ≤-1 kcal mol^-1^) (Fig. 2B, Table 1).

To examine the physicochemical nature of key residues critical for STX binding further, we made mutations at select positions guided by the alanine scan. Mutations at Glu540 and Asp794 (*10*), residues involved in charge pair interactions with the STX guanidinium rings, that neutralized the sidechain while preserving shape and volume (Fig. 2A, Table 1) disrupted binding strongly, similar to their alanine counterparts (ΔΔG = 4.30 and 3.60 kcal mol^-1^, for E540Q and D794N, respectively) (Figs. 2A, S4, Table 1). Altering sidechain length while preserving the negative charge at these sites also greatly diminished STX affinity, but was notably less problematic at Glu540 (ΔΔG = 1.54 and 2.94 kcal mol^-1^, for E540D and D794E, respectively). To probe contacts with Phe784, which makes a cation-π interaction (*17*) with the STX five-membered guanidinium ring (*10*), we tested changes that preserved this interface (F784Y), maintained sidechain volume and hydrophobicity (F784L), and that mimicked substitutions (F784C and F784S) found in the analogous residue in STX-resistant Na_V_s (Na_V_1.5, Na_V_1.8 and Na_V_1.9) (*10, 17, 19-22*) (Fig. 2A, Table 1). Preserving the cation-π interaction with F784Y caused a modest binding reduction (ΔΔG = 0.37 kcal mol^-1^), whereas F784L was disruptive (ΔΔG = 1.11 kcal mol^-1^) and F784C and F784S were even more destabilizing than F784A (ΔΔG = 3.15, 3.60, and 2.71 kcal mol^-1^, respectively).

We also examined two other positions that form part of the Sxph binding pocket near the five-membered STX guanidinium ring. Asp785 undergoes the most dramatic conformational change of any residue associated with STX binding moving from an external facing conformation to one that engages this STX element (*10*). Surprisingly, D785A and D785N mutations caused only relatively modest binding changes (Fig. 2A, Table 1) (ΔΔG = 0.57 and −0.30 kcal mol^-1^, for D785A and D785N, respectively). Because of the proximity of the second shell residue Gln787 to Asp785 and Asp794 (Fig. 2B), two residues that coordinate the five-membered STX guanidinium ring (*10*), we also asked whether adding additional negative charge to this part of the STX binding pocket would enhance toxin binding affinity. However, Q787E had essentially no effect on binding (ΔΔG = −0.09 kcal mol^-1^).

Two residues, Tyr558 and Ile782, stood out as sites where alanine substitutions enhanced STX affinity (Figs. 2A-B, Table 1). Tyr558 interacts with both the STX five-membered guanidinium ring and carbamate and moves away from the STX binding pocket upon toxin binding (*10*), whereas Ile782 is a second shell site that buttresses Tyr558. Hence, we hypothesized that affinity enhancements observed in the Tyr558 and Ile782 mutants resulted from the reduction of Tyr558-STX clashes. In accord with this idea, Y558F had little effect on STX binding (ΔΔG = −0.10 kcal mol^-1^), whereas shortening the sidechain but preserving its hydrophobic character, Y558I, enhanced binding as much as Y558A (ΔΔG =-1.07 and −1.00 kcal mol^-1^, respectively). Conversely, increasing the sidechain volume at the buttressing position, I782F, a change expected to make it more difficult for Tyr558 to move out of the binding pocket, reduced STX binding affinity (ΔΔG = 0.46 kcal mol^-1^). Combining the two affinity enhancing mutants, Y558A/I782A, yielded only a marginal increase in affinity in comparison to Y558A (ΔΔG =-1.07 and −1.00 kcal mol^-1^, respectively) but was better than I782A alone (ΔΔG = −0.53 kcal mol^-1^). This non-additivity in binding energetics (*36*) is in line with the physical interaction of the two sites and the direct contacts of Tyr558 with the toxin. Together, these data support the idea that the Tyr558 clash with STX is key factor affecting STX affinity and suggest that it should be possible to engineer Sxph variants with enhanced binding properties by altering this site.

Taken together, these studies of the energetic map of the *Rc*Sxph STX binding pocket highlight the importance of two amino acid triads. One (Glu540, Phe784, and Asp794) engages the STX bis-guanidinium core of the toxin. The second (Tyr558, Phe561, and Pro727), forms the surface surrounding the carbamate unit (Fig. 2B). The central role of the Glu540/Phe784/Asp794 triad in the energetics of binding the bis-guanidinium STX core underscores the toxin receptor site similarities between *Rc*Sxph and Na_V_s (*10*)(Figs. 2C-D). In both, STX binding relies on two acidic residues that coordinate the five and six-membered STX rings (Sxph Asp794 and Glu540, and rat Na_V_1.4 Glu403 and Glu758 (*18*)), and a cation-π interaction (Sxph Phe784 and ratNa_V_1.4 Tyr401 (*18*) and its equivalents in other Na_V_s (*19–22*)). Hence, both the basic architecture and binding energetics appear to be conserved even though the overall protein structures presenting these elements are dramatically different.

### Structures of enhanced affinity RcSxph mutants

To investigate the structural underpinnings of the affinity enhancement caused by mutations at the Tyr558 site, we determined crystal structures of *Rc*Sxph-Y558A and *Rc*Sxph-Y558I alone (2.60Å and 2.70Å resolution, respectively) and as co-crystallized STX complexes (2.60Å and 2.15Å, respectively) (Fig. S6A-D, Table S1). Comparison of the apo- and STX-bound structures reveals little movement in the STX binding pocket upon ligand binding (RMSD_Cα_ = 0.209 Å and 0.308 Å comparing apo- and STX-bound *Rc*Sxph-Y558A and *Rc*Sxph-Y558I, respectively) (Figs. 3A-B, Supplementary movies, M1 and M2). In both, the largest conformational change is the rotation of Asp785 into the binding pocket to interact with the five-membered guanidinium ring of STX, as seen for *Rc*Sxph (Fig. 3C) (*10*). By contrast, unlike in *Rc*Sxph, there is minimal movement of residue 558 and its supporting loop, indicating that both Y558A and Y558I eliminate the clash incurred by the Tyr558 sidechain. Comparison with the *Rc*Sxph:STX complex also shows that the STX carbamate in both structures has moved into a pocket formed by the mutation at Tyr558 (RMSD_Cα_ = 0.279 Å and 0.327 Å comparing *Rc*Sxph:STX with *Rc*Sxph-Y558A:STX and *Rc*Sxph-Y558I:STX, respectively) (Fig. 3C). This structural change involves a repositioning of the carbamate carbon by 2 Å in the *Rc*Sxph-Y558I:STX complex relative to the *Rc*Sxph:STX complex. These findings are in line with the nearly equivalent toxin binding affinities of Y558A and Y558I, as well as with the idea that changes at the Tyr558 buttressing residue, Ile782, relieve the steric clash with STX. They also demonstrate that one strategy for increasing STX affinity is to engineer a highly organized binding pocket that requires minimal conformational changes to bind STX.

**Figure 3.**
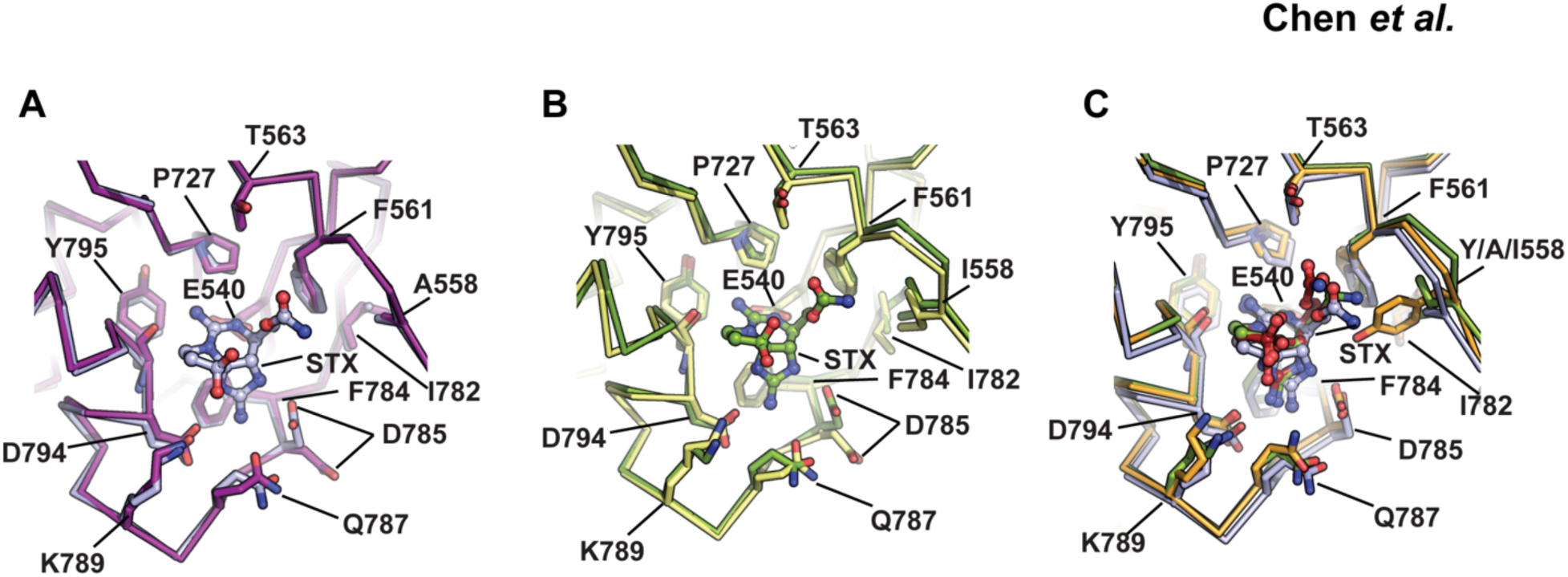
Structures of enhanced affinity *Rc*Sxph mutants. **A,** Superposition of the STX binding pockets of *Rc*Sxph-Y558A (purple) and the *Rc*Sxph-Y558A:STX complex (light blue). **B,** Superposition of the STX binding pockets of *Rc*Sxph-Y558I (pale yellow) and the *Rc*Sxph-Y558I:STX complex (splitpea). **C,** Superposition of the STX binding pockets of STX bound complexes of *Rc*Sxph (PDB: 6O0F) (*10*), *Rc*Sxph-Y558A (purple), and *Rc*Sxph-Y558I (splitpea). STX from the *Rc*Sxph is firebrick.

### Sxph STX binding affinity changes alter Na_v_ rescue from STX block

*Rc*Sxph acts as a ‘toxin sponge’ that can reverse STX inhibition of Na_V_s (*9*). To test the extent to which this property is linked to the intrinsic affinity of *Rc*Sxph for STX, we evaluated how STX affinity altering mutations affected *Rc*Sxph rescue of channels blocked by STX. As shown previously, titration of different *Rc*Sxph:STX ratios against *Phyllobates terribilis* Na_V_1.4 (*Pt*Na_V_1.4), a Na_V_ having an IC_50_ for STX of 12.6 nM (*9*), completely reverses the effects of STX at ratios of 2:1 *Rc*Sxph:STX or greater (Figs. 4A and F). Incorporation of mutations that affect STX affinity altered the ability of *Rc*Sxph to rescue Na_V_s and followed the binding assay trends. Mutants that increased STX affinity, Y558I and I782A, improved the ability of *Rc*Sxph to rescue *Pt*Na_V_1.4 (Effective Rescue Ratio_50_ (ERR_50_)= 0.81 ± 0.01, 0.87 ± 0.02, and 1.07 ± 0.02 for Y558I, I782A, and *Rc*Sxph, respectively), whereas mutations that compromised STX binding reduced (P727A, ERR_50_ >4) or eliminated (E540A) the ability of *Rc*Sxph to reverse the STX inhibition (Fig. 4B-F). This strong correlation indicates that the ‘toxin sponge’ property of Sxph (*9*) depends on the capacity of Sxph to sequester STX and adds further support to the idea that Sxph has a role in toxin resistance mechanisms (*8, 9*).

**Figure 4.**
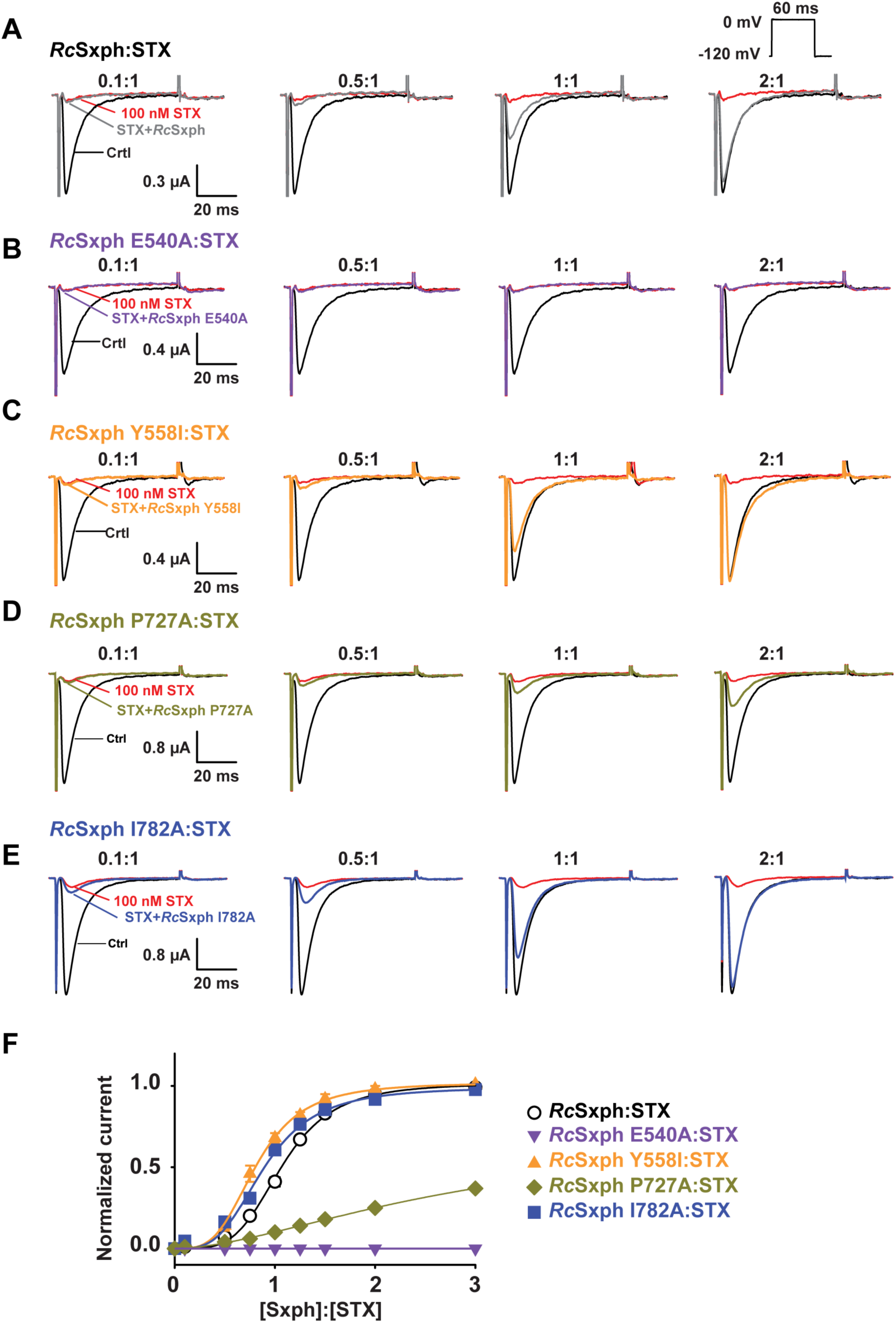
*Rc*Sxph mutants have differential effects on *Pt*Na_V_1.4 rescue from STX block. **A-E,** Exemplar two-electrode voltage clamp recordings of *Pt*Na_V_1.4 expressed in *Xenopu*s oocytes in the presence of 100 nM STX and indicated [Sxph]:[STX] ratios for **A,** *Rc*Sxph, **B,** *Rc*Sxph E540A, **C,** *Rc*Sxph Y558I, **D,** *Rc*Sxph P727A, and **E,** *Rc*Sxph I782A. Inset shows the stimulation protocol. **F,** [Sxph]:[STX] dose response curves for *Rc*Sxph (black, open circles), *Rc*Sxph E540A (purple inverted triangles), *Rc*Sxph Y558I (orange triangles), *Rc*Sxph P727A (gold diamonds), and *Rc*Sxph I782A (blue squares) in the presence of 100 nM STX. Lines show fit to the Hill equation.

### Expansion of the Sxph family

STX binding activity has been reported in the plasma, hemolymph, and tissues of diverse arthropods, amphibians, fish, and reptiles (*11, 13*), suggesting that many organisms harbor Sxph-like proteins. Besides *Rc*Spxh, similar Sxphs have been identified in only two other frogs, the High Himalaya frog *Nanorana parkeri* (*10*) and the little devil poison frog *Oophaga sylvatica* (*24*). As a number of poison frogs exhibit resistance to STX poisoning (*9*), we asked whether the STX binding site ‘recognition code’ could enable identification of Sxph homologs in other amphibians. To this end, we determined the sequences of ten new Sxphs (Figs. 5A and B, S7, and S8). These include six Sxphs in two poison dart frog families (Family *Dendrobatidae*: Dyeing poison dart frog, *Dendrobates tinctorius*; Little devil poison frog, *O. sylvatica*; Mimic poison frog*, Ranitomeya imitator*; Golden dart frog*, Phyllobates terribilis*; Phantasmal poison frog, *Epipedobates tricolor*; and Brilliant-thighed poison frog, *Allobates femoralis*; and Family: Mantellidae Golden mantella*, Mantella aurantiaca*), and three Sxphs in toads (Caucasian toad, *Bufo bufo;* Asiatic toad, *Bufo gargarizans*; and South American cane toad*, Rhinella marina*). The identification of the *Os*Sxph sequence confirms its prior identification by mass spectrometry (*24*) and the discovery of *Rm*Sxph agrees with prior reports of Sxph-like STX binding activity in the cane toad (*R. marinus*)(*13, 23*).

**Figure 5.**
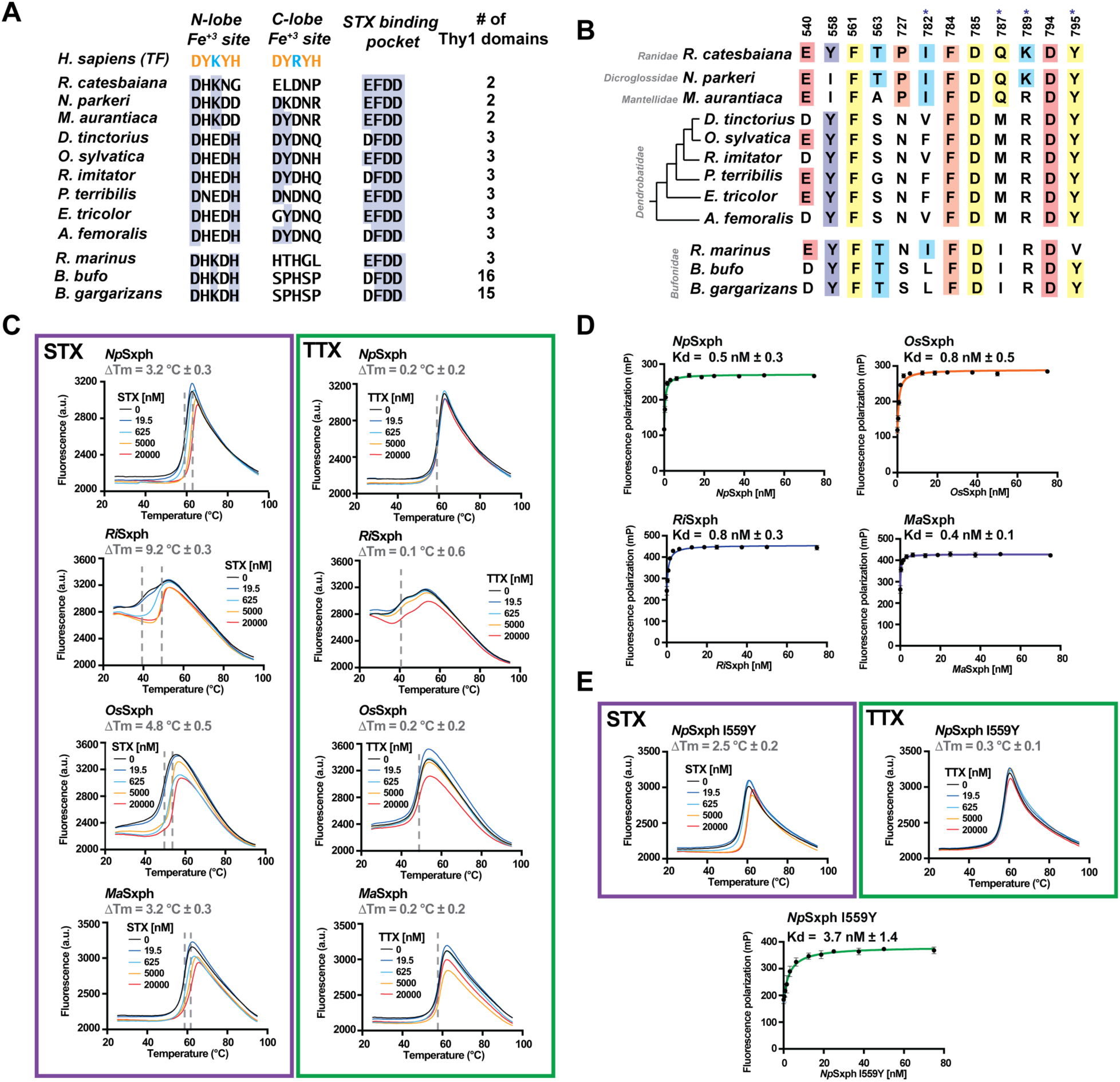
Sxph family member properties. **A**, Comparison of the human transferrin (TF) Fe^3+^ ligand positions (UniProtKB: P02787), *Rc*Sxph STX binding motif residues (*10*), and number of Thy1 domains for Sxphs from *R. castesbeiana* (PDB:6O0D) (*10*), *N. parkeri* (NCBI: XP_018410833.1) (*10*), *M. aurantiaca*, *D. tinctorius*, *O. sylvatica*, *R. imitator*, *P. terribilis*, *E. tricolor*, *A. femoralis*, *R. marinus*, *B. bufo* (NCBI:XM_040427746.1), and *B. garagarizans*. (NCBI:XP_044148290.1). TF Fe^3+^ (orange) and carbonate (blue) ligands are indicated. Blue highlights indicate residue conservation. **B,** Comparison of STX binding pocket for the indicated Sxphs. Numbers denote *Rc*Sxph positions. Colors indicate the alanine scan classes as in Fig. 2B. Conserved residues are highlighted. Asterix indicates second shell sites. **C,** Exemplar TF curves for *Np*Sxph, *Ri*Sxph, *Os*Sxph, and *Ma*Sxph in the presence of the indicated concentrations of STX (purple box) or TTX. (green box) ΔTm values are indicated. **D,** Exemplar FP binding curves and Kds for *Np*Sxph (green), *Ri*Sxph (blue), *Os*Sxph (orange), and *Ma*Sxph (purple) . **E,** Exemplar *Np*Sxph I559Y TF curves in the presence of the indicated concentrations of STX (purple box) or TTX (green box) and FP binding (green). ΔTm and Kd values are indicated. Error bars are S.E.M.

Sequence comparisons (Figs. S7 and S8) show that all of the new Sxphs share the transferrin fold found in *Rc*Sxph comprising N-and C-lobes each having two subdomains (N1, N2 and C1, C2, respectively) (*10, 12*) and the signature ‘EFDD’ motif (*10*) or a close variant in the core of the C-lobe STX binding site (Fig. 5A). Similar to *Rc*Sxph, the new Sxphs also have amino acid differences relative to transferrin that should eliminate Fe^3+^ binding (*10, 12, 37*), as well as a number of protease inhibitor thyroglobulin domains (Thy1) inserted between the N1 and N2 N-lobe subdomains (*10, 38*) (Fig. 5A and S7-S9). These Thy1 domain insertions range from two in *Rc*Sxph, *Np*Sxph, and *Ma*Sxph, to three in the dendrobatid poison frog and cane toad Sxphs, to 16 and 15 in toad *Bb*Sxph and *Bg*Sxph, respectively (Figs. 5A and S7-S9).

We used the STX recognition code defined by our studies as a template for investigating cross-species variation in the residues that contribute to STX binding (Fig. 5B). This analysis shows a conservation of residues that interact with the STX bis-guanidium core (Glu540, Phe784, Asp785, Asp794, and Tyr795) and carbamate (Phe561). Surprisingly, five of the Sxphs (*D. tinctorius*, *R. imitator*, *A. femoralis*, *B. bufo,* and *B. gargarizans*) have an aspartate instead of a glutamate at the Glu540 position in *Rc*Sxph that contributes the most binding energy (Fig. 2A). The equivalent change in *Rc*Sxph, E540D, reduced STX affinity by ∼100 fold (Table 1) and uniquely alters enthalpy and entropy binding parameters compared to other affinity-lowering mutations (Table S2). Additionally, we identified variations at two sites for which mutations increase *Rc*Sxph STX binding, Tyr558 and Ile782 (Figs. 2A, 4C, 4E, and Table 1). *Np*Sxph and *Ma*Sxph have an Ile at the Tyr558 site, whereas eight of the new Sxphs have hydrophobic substitutions at the Ile782 position (Fig. 5B). The striking conservation of the Sxph scaffold and STX binding site indicate that this class of ‘toxin sponge’ proteins is widespread among diverse Anurans, while the amino acid variations in key positions (Glu540, Tyr558, and Ile782), raise the possibility that the different Sxph homologs have varied STX affinity or selectivity for STX congeners.

### Diverse Sxph family members have conserved STX binding properties

To explore the STX binding properties of this new set of Sxphs and to begin to understand whether changes in the binding site composition affect toxin affinity, we expressed and purified four representative variants. These included two Sxphs having STX binding site sequences similar to *Rc*Sxph (*Np*Sxph and *Ma*Sxph) and two Sxphs bearing more diverse amino acid differences (*Ri*Sxph and *Os*Sxph), including one displaying the E540D substitution (*Ri*Sxph). This set also represents Sxphs having either two Thy1 domains similar to *Rc*Sxph (*Np*Sxph and *Ma*Sxph) or three Thy1 domains (*Os*Sxph and *Ri*Sxph) (Fig. 5A). TF experiments showed STX-dependent ΔTms for all four Sxphs. By contrast, equivalent concentrations of TTX had no effect (Fig. 5C), indicating that, similar to *Rc*Spxh (Fig. 1A) (*8, 9*), all four Sxphs bind STX but not TTX. Unlike the other Sxphs, the *Ri*Sxph melting curve showed two thermal transitions; however, only the first transition was sensitive to STX concentration (Fig. 5C). FP binding assays showed that all four Sxphs bound F-STX and revealed affinities stronger than *Rc*Sxph (Fig. 5D and Table 2). The enhanced affinity of *Np*Sxph and *Ma*Sxph for STX relative to *Rc*Sxph is consistent with the presence of the Y558I variant (Fig. 5B). Importantly, the observation that *Ri*Sxph has a higher affinity for STX than *Rc*Sxph despite the presence of the E540D difference suggests that the other sequence variations in the *Ri*Sxph STX binding pocket compensate for this Glu◊Asp change at Glu540.

**Table 2.**
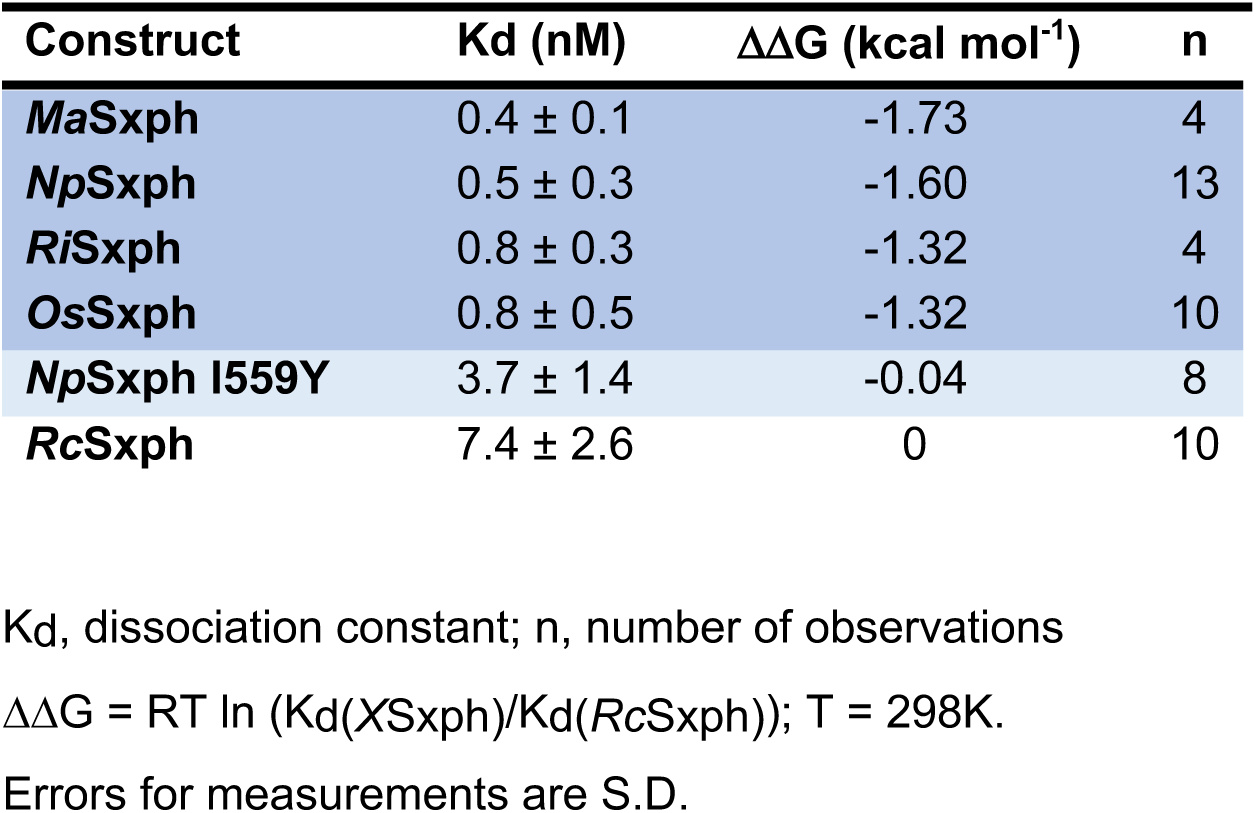
Comparison of Sxph STX binding properties.

| **Construct** | **Kd (nM)** | **$\Delta\Delta G$ (kcal mol<sup>-1</sup>)** | **n** |
| --- | --- | --- | --- |
| ***MaSxph*** | 0.4 ± 0.1 | -1.73 | 4 |
| ***NpSxph*** | 0.5 ± 0.3 | -1.60 | 13 |
| ***RiSxph*** | 0.8 ± 0.3 | -1.32 | 4 |
| ***OsSxph*** | 0.8 ± 0.5 | -1.32 | 10 |
| ***NpSxph I559Y*** | 3.7 ± 1.4 | -0.04 | 8 |
| ***RcSxph*** | 7.4 ± 2.6 | 0 | 10 |

Kd, dissociation constant; n, number of observations
$\Delta\Delta G = RT \ln (Kd(XSxph)/Kd(RcSxph))$ ; T = 298K.
Errors for measurements are S.D.

Because *Np*Sxph has a higher affinity for STX than *Rc*Sxph (Figs. 1D, 5D and Table 2) and has an isoleucine at the Tyr558 site (Fig. 5B), we asked whether the *Np*Sxph I559Y mutant that converts the *Np*Sxph binding site to match *Rc*Sxph would lower STX affinity. TF measurements showed that *Np*Sxph I559Y had a ∼1°C smaller ΔTm than *Np*Sxph (ΔTm = 3.2°C ± 0.3 versus 2.5°C ± 0.2 for *Np*Sxph and *Np*Sxph I559Y, respectively), indicative of a decreased binding affinity (Figs. 5C and E). This result was validated by FP (ΔΔG =-1.56 kcal mol^-1^), yielding a result of similar magnitude to the *Rc*Sxph Y558I differences (Fig. 5E, Tables 1 and 2). ITC confirmed the high affinity of the interaction (Fig. S5D-F), but could not yield an explicit Kd given its low nanomolar value (Fig. S5F). Nevertheless, these experiments validate the 1:1 stoichiometry of the *Np*Sxph:STX interaction (Table S2) and show that the I559Y change reduced the binding enthalpy, consistent with perturbation of *Np*Sxph:STX interactions (ΔH =-18.7 ± 0.2 vs. −16.8 ± 0.2 kcal mol^-1^, *Np*Sxph and *Np*Sxph I559Y, respectively) (Table S2). Taken together, these experiments establish the conserved nature of the STX binding pocket among diverse Sxph homologs and show that the STX recognition code derived from *Rc*Sxph studies (Fig. 5B) can identify key changes that influence toxin binding.

### Structures of apo- and STX bound NpSxph reveal a pre-organized STX binding site

To compare STX binding modes among Sxph family members, we crystallized and determined the structure of *Np*Sxph, alone and co-crystallized with STX. *Np*Sxph and STX:*Np*Sxph crystals diffracted X-rays to resolutions of 2.2Å and 2.0Å, respectively, and were solved by molecular replacement (Figs. 6A, S10A and B). As expected from the similarity to *Rc*Sxph, *Np*Sxph is built on a transferrin fold (Fig. 6A) and has the same 21 disulfides found in *Rc*Sxph, as well as an additional 22^nd^ disulfide in the Type 1A thyroglobulin domain of *Np*Sxph Thy1-2. However, structural comparison of *Np*Sxph and *Rc*Sxph reveals a number of unexpected large-scale domain rearrangements.

**Figure 6.**
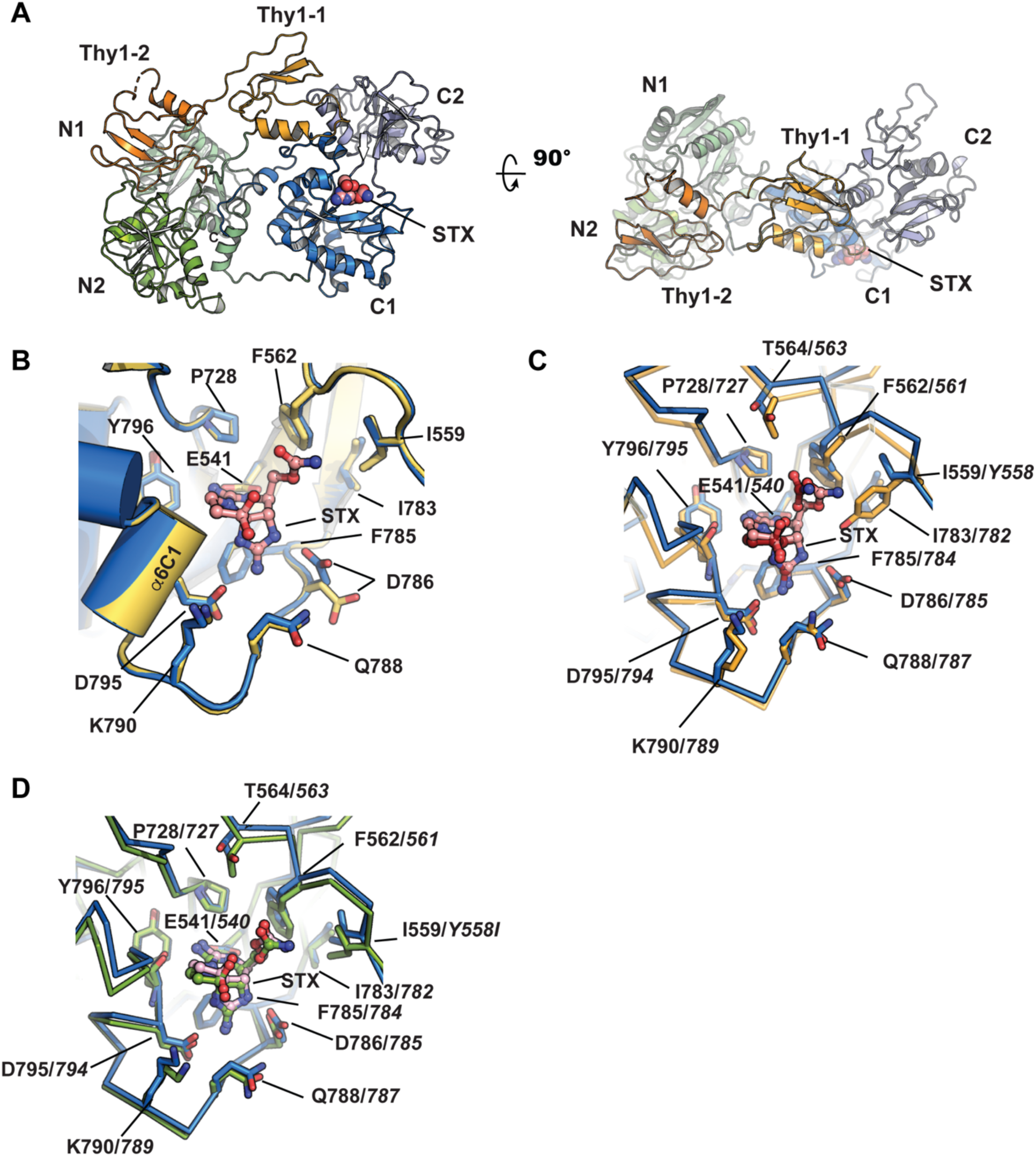
*Np*Sxph and *Np*Sxph:STX structures. **A**, Cartoon diagram of the *Np*Sxph:STX complex. N1 (light green), N2 (green), Thy1-1 (light orange), Thy1-2 (orange), C1 (marine), and C2 (light blue) domains are indicated. STX (pink) is shown in space filling representation. **B,** Comparison of STX binding pocket for apo-*Np*Sxph (yellow) and *Np*Sxph:STX (marine). STX (pink) is shown as ball and stick. **C,** Comparison of *Np*Sxph (marine) and *Rc*Sxph (orange) (PDB:6O0F) (*10*) STX binding sites. STX from *Np*Sxph and *Rc*Sxph complexes is pink and orange, respectively. **D**, Comparison of *Np*Sxph (marine) and *Rc*Sxph-Y558I (splitpea) STX binding sites. STX from *Np*Sxph and *Rc*Sxph-Y558I complexes is pink and splitpea, respectively. *Rc*Sxph and *Rc*Sxph-Y558I residue numbers in ‘**C**’ and ‘**D**’ are indicated in italics.

The *Np*Sxph N-lobe is displaced along the plane of the molecule by ∼30° and rotated around the central axis by a similar amount (Fig. S10C). *Np*Sxph N-lobe and C-lobe lack Fe^3+^ binding sites (Fig. 5A), and despite the N-lobe displacement relative to *Rc*Sxph adopt closed and open conformations, respectively as in *Rc*Sxph (*10*) (Fig. S10D-E) (RMSD_Cα_ = 1.160 Å and 1.373 Å for *Np*Sxph and *Rc*Sxph N- and C-lobes, respectively). Surprisingly, the two *Np*Sxph Thy1 domains are in different positions than in *Rc*Sxph and appear to move as a unit by ∼90° with respect to the central transferrin scaffold (Fig. S10F and Supplementary movie M3) and a translation of ∼30Å of Thy1-2 (Fig. S10G). Thy1-1 is displaced from a site over the N-lobe in *Rc*Sxph to one in which it interacts with the *Np*Sxph C-lobe C2 subdomain and Thy1-2 moves from between the N and C-lobes in *Rc*Sxph where it interacts with the C1 subdomain, to a position in *Np*Sxph where it interacts with both N-lobe subdomains. Consequently, the interaction between the C-lobe β−strand β7C1 and Thy1-2 β5 observed in *Rc*Sxph is absent in *Np*Sxph. Despite these domain-scale differences, Thy1-1 and Thy1-2 are structurally similar to each other (RMSD_Cα_ = 1.056 Å) and to their *Rc*Sxph counterparts (Fig. S10H) (RMSD_Cα_ = 1.107 Å and 0.837, respectively). Further, none of these large scale changes impact the STX binding site, which is found on the C1 domain as in *Rc*Sxph (Fig. 5A).

Comparison of the apo- and STX-bound *Np*Sxph structures shows that there are essentially no STX binding site conformational changes upon STX engagement, apart from the movement of Asp786 to interact with the STX five-membered guanidinium ring (Fig. 6B and Supplementary Movie M4). This conformational change is shared with *Rc*Sxph (*10*) and appears to be a common element of Sxph binding to STX. The movements of Tyr558 and its loop away from the STX binding site observed in *Rc*Sxph (*10*) are largely absent in *N*pSxph for the Tyr558 equivalent position, Ile559, and its supporting loop. Hence, the *Np*Sxph STX binding site is better organized to accommodate STX (Fig. 6B), similar to *Rc*Sxph Y558I (Fig. 3B). We also noted an electron density in the apo-*Np*Sxph STX binding site that we assigned as a PEG400 molecule from the crystallization solution (Fig. S10A). This density occupies a site different from STX and is not present in the STX-bound complex (Fig. S10B). Its presence suggests that other molecules may be able to bind the STX binding pocket.

We also determined the structure of an *Np*Sxph:F-STX complex at 2.2Å resolution (Table S1). This structure shows no density for the fluorescein moiety and has an identical STX pose to the *Np*Sxph:STX complex (Fig. S11). These data provide further evidence that fluorescein does not interact with Sxph (cf. Fig. S3) even though it is tethered to the STX binding pocket and support the idea that the FP assay faithfully reports on Sxph:STX interactions. Comparison of the *Np*Sxph and *Rc*Spxh STX poses shows essentially identical interactions with the tricyclic bis-guanidium core and reveals that the carbamate is able to occupy the pocket opened by the Y◊I variant (Fig. 6C), as observed in *Rc*Sxph Y558I (Fig. 3B). This change, together with the more rigid nature of the *Np*Sxph STX binding pocket likely contributes to the higher affinity of *Np*Sxph for STX relative to *Rc*Sxph (Table 1). Taken together, the various structures of Sxph:STX complexes show how subtle changes, particularly at the Tyr558 position can influence STX binding and underscore that knowledge of the STX binding code can be used to tune the STX binding properties of different Sxphs.

## Discussion

Our biochemical and structural characterization of a set of *Rc*Sxph mutants and Sxphs from diverse Anurans reveals a conserved STX recognition code centered around six amino acid residues comprising two triads. One triad engages the STX bis-guanidinium core using carboxylate groups that coordinate each ring (*Rc*Sxph Glu540 and Asp794) and an aromatic residue that makes a cation-π interaction (*Rc*Sxph Phe784) with the STX concave face. This recognition motif is shared with Na_V_s, the primary target of STX in PSP (*18, 39, 40*) (Fig. 2C and D), and showcases a remarkably convergent STX recognition strategy. The second amino acid triad (*Rc*Sxph residues Tyr558, Phe561, and Pro727) largely interacts with the carbamate moiety and contains a site, Tyr558, and its supporting residue, Ile782, where amino acid changes, including those found in some anuran Sxphs (Fig. 5), enhance STX binding. Structural studies of *Rc*Sxph mutants and the High Himalaya frog *Np*Sxph show that STX-affinity enhancing substitutions in this area of the binding pocket act by reducing the degree of conformational change associated with STX binding (Figs. 3 and 6 C-D). These findings reveal one strategy for creating high affinity STX binding sites. Importantly, enhancing the affinity of Sxph for STX through changes at either residue increases the capacity of *Rc*Sxph to rescue Na_V_s from STX block (Fig. 4), demonstrating that an understanding of the STX recognition code enables rational modification of Sxph binding properties. Thus, exploiting the information in the STX recognition code defined here should enable design of Sxphs as STX sensors or agents for treating STX poisoning.

Although STX binding activity has been reported in a variety of diverse invertebrates (*13*) and vertebrates (*13, 23*), only two types of STX binding proteins have been identified and validated, frog Sxphs (*8, 9*) and the pufferfish STX and TTX binding proteins (*41, 42*). Our discovery of a set of ten new Sxphs that bind STX with high affinity (Figs. 5, S7, and S8) represents a substantial expansion of the Sxph family and reveals natural variation in the residues that are important for STX binding (Fig. 5B). Most notably, E540D, a change that reduces STX binding in *Rc*Sxph by ∼14-fold, occurs in five of the newly identified Sxphs. Nevertheless, functional studies show that *Ri*Sxph, which bears an Asp at this site, binds STX more strongly than *Rc*Sxph (Table 2). Hence, the natural variations at other STX binding pocket residues must provide compensatory interactions to maintain a high STX binding affinity. Understanding how such variations impact STX engagement or influence the capacity of these proteins to discriminate among STX congeners (*13*) remain important unanswered questions. The striking abundance of Sxphs in diverse amphibians, representing linages separated by ∼140 million years (*25*) and that are not known to carry STX, raises intriguing questions regarding the selective pressures that have caused these disparate amphibians to maintain this STX binding protein and its capacity to sequester this lethal toxin.

Besides the conserved STX binding site, all of the amphibian Sxphs possess a set of Thy1 domains similar to those in *Rc*Sxph that have been shown to act as protease inhibitors (*38*). Comparison of anuran Sxphs shows that these domains are a common feature of the Sxph family and occur in strikingly varied numbers, comprising of 2-3 in most Sxphs but having a remarkable expansion to 15-16 in some toad Sxphs (Figs. 5A, and S7-S9). Structural comparisons between *Rc*Sxph and *Np*Sxph, representing the class that has two Thy1 domains (Fig. 5A), show that these domains can adopt different positions with respect to the shared, modified transferrin core (Fig. S10F). Whether the Sxph Thy1 domains and their varied numbers are important for Sxph-mediated toxin resistance mechanisms (*8, 9*) or serve some other function remains unknown. Our definition of the Sxph STX binding code, which provides a guide for deciphering variation in the Sxph STX binding site (Fig. 5B), and discovery of high variability in Thy1 repeats among Anuran Sxphs constitutes a template for identifying other Sxphs within this widespread and diverse family of amphibians.

STX interacts with a variety of target proteins including select Na_V_ isoforms (*43*), other channels (*44, 45*), diverse soluble STX binding proteins (*8, 10, 41, 42, 46, 47*), and some enzymes (*3, 48, 49*). The identification of the Sxph STX recognition code together with the substantial expansion of the Sxph family provides a foundation for developing a deeper understanding of the factors that enable proteins to bind STX. Exploration of such factors should be facilitated by the assays established here that allow Sxph:STX interactions to be probed using a range of sample quantities (TF: 25 µg Sxph;600 ng STX; FP: 3 µg Sxph;1 ng F-STX, ITC: 300 µg Sxph;5 µg STX) and that should be adaptable to other types of STX targets. In cases of limited samples, such as difficult to obtain STX congeners, the excellent agreement among the assays should provide a reliable basis for interpretation of Sxph binding properties. Our delineation of the Sxph STX binding code and discovery of numerous Sxph family homologs among diverse amphibians sets a broad framework for understanding the lethal effects of this potent neurotoxin and ‘toxin sponge’ STX resistance mechanisms (*8, 9*). This knowledge may enable the design of novel PSP toxin sensors and agents that could mitigate STX intoxication.

## Supporting information

Movie M1 RcSxph-Y558A conformational changes upon STX binding.

Movie M2 RcSxph-Y558I conformational changes upon STX binding.

Movie M3 Conformational changes between RcSxph and NpSxph.

Movie M4 NpSxph conformational changes upon STX binding.

## Acknowledgements

We thank Z. Wong and T.-J. Yen for technical help and K. Brejc and J. Gross for comments on the manuscript. This work was supported by grants DoD HDTRA-1-19-1-0040, HDTRA-1-21-1-10011, and UCSF Program for Breakthrough Biomedical Research, which is partially funded by the Sandler Foundation, to D.L.M., NSF-1822025 to L.A.O, NIH-NIGMS R01-GM117263-05 to J.D.B., an American Heart Association postdoctoral fellowship to F.A.-A, an NSF GRFP (DGE-1656518) and HHMI Gilliam Fellowship (GT13330) to A.A.-B., and a Department of Defense (DoD) National Defense Science and Engineering Graduate (NDSEG) Fellowship to H.S.H. H.S.H. is a Center for Molecular Analysis and Design Fellow at Stanford University.

## Author Contributions

Z.C., S.Z., H.H., A.A.-B., L.A.O., J.D.B., and D.L.M. conceived the study and designed the experiments. Z.C. established the TF and FP assays, made and characterized Sxph mutants, and determined crystal structures of *Rc*Sxph complexes and mutants. H.S.H. and J.D.B. designed and synthesized F-STX. S.Z. expressed, characterized, and determined the structures of *Np*Sxph complexes and characterized *Ri*Sxph, *Ma*Sxph, and *Os*Sxph. A.A.-B., M.B., D.R., L.A.O., and D.L.M. identified frog Sxphs. Z.C. and A.B.-B. cloned and sequenced *Pt*Sxph and *Rm*Sxph. F. A.-A. performed two-electrode voltage clamp electrophysiology experiments and analyzed the data. J.D.B., L.A.O., and D.L.M. analyzed data and provided guidance and support. Z.C., S.Z., H.S.H., A.A.-B., L.A.O., J.D.B., and D.L.M. wrote the paper.

## Competing interests

J.D.B. is a cofounder and holds equity shares in SiteOne Therapeutics, Inc., a start-up company interested in developing subtype-selective modulators of sodium channels.

The other authors declare no competing interests.

## Data and materials availability

Sequences of MaSxph, DtSxph, OsSxph, RiSxph, PtSxph, EtSxph, AfSxph, RmSxph, BbSxph, and BgSxph will be deposited with NCBI and released upon publication.

Coordinates and structure factors and for RcSxph-Y558A (8D6P), RcSxph-Y558A:STX (8D6S), RcSxph Y558I (8D6Q), RcSxph-Y558I:STX (8D6T), RcSxph:F-STX (8D6U), NpSxph (8D6G), NpSxph:STX (8D6M), and NpSxph:F-STX (8D6O) are deposited with the PDB and will be released upon publication.

Requests for material to D.L.M.

## Materials and Methods

### Expression and purification of Sxphs and mutants

*R. catesbeiana* Sxph (*Rc*Sxph) and mutants were expressed using a previously described *Rc*Sxph baculovirus expression system in which *Rc*Sxph carries in series, a C-terminal 3C protease cleavage site, green fluorescent protein (GFP), and a His_10_ tag (*10*). The gene encoding *Nanorana parkeri* Sxph (*Np*Sxph) including its N-terminal secretory sequence (GenBank: XM_018555331.1) was synthesized and subcloned into a pFastBac1 vector using NotI and XhoI restriction enzymes by GenScript and bears the same C-terminal tags as *Rc*Sxph. *Rc*Sxph and *Np*Sxph mutants were generated using the QuikChange site-directed mutagenesis kit (Stratagene). All constructs were sequenced completely. *Rc*Sxph, *Rc*Sxph mutants, *Np*Sxph and *Np*Sxph I559Y were expressed in *Spodoptera frugiperda* (Sf9) cells using a baculovirus expression system as described previously for *Rc*Sxph (*10*) and purified using a final size exclusion chromatography (SEC) run in 150 mM NaCl, 10 mM HEPES, pH 7.4. Protein concentrations were determined by measuring UV absorbance at 280 nm using the following extinction coefficients calculated using the ExPASY server (https://web.expasy.org/protparam/): *Rc*Sxph Y558 mutants, 94,875 M^−1^ cm^−1^; *Rc*Sxph F784C 96,490 M^−1^ cm^−1^; *Rc*Sxph F784Y 97,855 M^−1^ cm^−1^, *Rc*Sxph and all other *Rc*Sxph mutants 96,365 M^−1^ cm^−1^; *Np*Sxph 108,980 M^−1^ cm^−1^; and *Np*Sxph I559Y 110,470 M^−1^ cm^−1^.

### Thermofluor (TF) assay of toxin binding

Thermofluor assays for STX and TTX binding were developed as outlined (*27*). TTX was purchased from Abcam (Catalog # ab120054). 20 µL samples containing 1.1 µM *Rc*Sxph, *Np*Sxph, or mutants thereof, 5x SYPRO Orange dye (Sigma-Aldrich, S5692, stock concentration 5000x), 0-20 µM STX or TTX, 150 mM NaCl, 10 mM HEPES, pH 7.4 were set up in 96-well PCR plates (Bio-Rad), sealed with a microseal B adhesive sealer (Bio-Rad) and centrifuged (1 min, 230xg) prior to thermal denaturation. The real-time measurement of fluorescence using the HEX channel (excitation 515-535 nm, emission 560-580 nm) was performed in CFX Connect Thermal Cycler (Bio-Rad). Samples were heated from 25°C to 95°C at 0.2°C min^-1^. Melting temperature (Tm) was calculated by fitting the denaturation curves using a Boltzmann sigmoidal function and GraphPad Prism: F=F_min_+(F_max_-F_min_)/(1+exp((Tm-T)/C)), where F is the fluorescence intensity at temperature T, F_min_ and F_max_ are the fluorescence intensities before and after the denaturation transition, respectively, Tm is the midpoint temperature of the thermal unfolding transition, and C is the slope at Tm (*27*). ΔTm=Tm_Sxph+20µM toxin_-Tm_Sxph_.

### Fluorescence polarization assay

Fluorescence polarization assays were performed as described (*50*). 100 µL samples containing 1 nM fluorescein labeled STX (F-STX), 150 mM NaCl, 10 mM HEPES, pH 7.4, and Sxph variants at the following concentration ranges (*Rc*Sxph and *Rc*Sxph T563A, I782A, F784Y, D785N, Q787A, Q787E, K789A, and Y795A, 0-75 nM; *Rc*Sxph Y558A and I782A/Y558A, 0-24 nM; *Rc*Sxph Y558I, 0-37.5 nM; *Rc*Sxph Y558F, 0-100 nM; *Rc*Sxph I782F 0-150 nM; *Rc*Sxph F561A, 0-300 nM; *Rc*Sxph F784L, 0-500 nM; *Rc*Sxph E540D, P727A, and D785A, 0-600 nM; *Rc*Sxph F784A, 0-4.8 µM; *Rc*Sxph E540A and F784C, 0-10 µM; *Rc*Sxph D794A and D794E, 0-12.5 µM; *Rc*Sxph F784S, 0-17 µM; *Rc*Sxph D794N, 0-20 µM; *Rc*Sxph E540Q, 0-25 µM; *Np*Sxph and *Np*Sxph I559Y, 0-75 nM ) were prepared in 96-well black flat-bottomed polystyrene microplates (Greiner Bio-One) and sealed with an aluminum foil sealing film (AlumaSeal II), and incubated at room temperature for 0.5 h before measurement. Measurements were performed at 25°C on a Synergy H1 microplate reader (BioTek) using the polarization filter setting (excitation 485 nm, emission 528 nm). Binding curves for representative high affinity (*Rc*Sxph, *Np*Sxph, and *Rc*Sxph-Y558I) and low affinity (*Rc*Sxph-E540D) proteins were compared at 0.5 h, 1.5 h, 4.5 h, and 24 h, post mixing and indicated that equilibrium was reached by 0.5 h for all samples. The dissociation constants were calculated using GraphPad Prism by fitting fluorescence millipolarization (mP=P·10^−3^, where P is polarization) as a function of Sxph concentration using the equation: P={(P_bound_−Pfree) [Sxph]/(K_d_+[Sxph])}+P_free_, where P is the polarization measured at a given Sxph concentration, P_free_ is the polarization of Sxph in the absence of F-STX, and P_bound_ is the maximum polarization of Sxph bound by F-STX (*50, 51*).

### Isothermal titration calorimetry (ITC)

ITC measurements were performed at 25°C using a MicroCal PEAQ-ITC calorimeter (Malvern Panalytical). *Rc*Sxph, *Rc*Sxph mutants, *Np*Sxph, and *Np*Sxph I559Y were purified using a final size exclusion chromatography step in 150 mM NaCl, 10 mM HEPES, pH 7.4. 1 mM STX stock solution was prepared by dissolving STX powder in MilliQ water. This STX stock was diluted with the SEC buffer to prepare 100 µM or 300 µM STX solutions having a final buffer composition of 135 mM NaCl, 9 mM HEPES, pH 7.4. To match buffers between the Sxph and STX solutions, the purified protein samples were diluted with MilliQ water to reach a buffer concentration of 135 mM NaCl, 9 mM HEPES, pH 7.4. (30 µM for *Rc*Sxph D794E;10 µM for *Rc*Sxph, other *Rc*Sxph mutants, *Np*Sxph, and *Np*Sxph I559Y) Protein samples were filtered through a 0.22 µm spin filter (Millipore) before loading into the sample cell and titrated with STX (300 µM STX for *Rc*Sxph D794 and 100 µM STX for *Rc*Sxph, other *Rc*Sxph mutants, *Np*Sxph, and *Np*Sxph I559Y) using a schedule of 0.4 µL titrant injection followed by 35 injections of 1 µL for the strong binders (*Rc*Sxph, *Rc*Sxph Y558I, *Rc*Sxph Y558A, *Rc*Sxph F561A, *Np*Sxph, and *Np*Sxph I559Y) and a schedule of 0.4 µL titrant injection followed by 18 injections of 2 µL for the weak binders (*Rc*Sxph P727A, *Rc*Sxph E540D, and *Rc*Sxph D794E). The calorimetric experiment settings were: reference power, 5 µcal/s; spacing between injections, 150 s; stir speed 750 rpm; and feedback mode, high. Data were analyzed using MicroCal PEAQ-ITC Analysis Software (Malvern Panalytical) using a single binding site model. The heat of dilution from titrations of 100 µM STX in 135 mM NaCl, 9 mM HEPES, pH 7.4 into 135 mM NaCl, 9 mM HEPES, pH 7.4 was subtracted from each experiment to correct the baseline.

### Crystallization, structure determination, and refinement

*Rc*Sxph mutants were crystallized at 4°C as previously described for *Rc*Sxph (*10*). Briefly, purified protein was exchanged into a buffer of 10 mM NaCl, 10 mM HEPES, pH 7.4 and concentrated to 65 mg ml^-1^ using a 50-kDa cutoff Amicon Ultra centrifugal filter unit (Millipore). Crystallization was set up by hanging drop vapor diffusion using a 24-well VDX plate with sealant (Hampton Research) using 3 µL drops having a 2:1 (v:v) ratio of protein:precipitant. For co-crystallization with STX, STX and the target *Rc*Sxph mutants were mixed in a molar ratio of 1.1:1 STX:Sxph and incubated on ice for 1 hour before setting up crystallization. *Rc*Sxph-Y558I and *Rc*Sxph-Y558I:STX were crystallized from solutions containing 27% 2-methyl-2,4-pentanediol, 5% PEG 8000, 0.08-0.2 M sodium cacodylate, pH 6.5. *Rc*Sxph-Y558A and *Rc*Sxph-Y558A:STX were crystallized from solutions containing 33% 2-methyl-2,4-pentanediol, 5% PEG 8000, 0.08-0.2 M sodium cacodylate, pH 6.5. To obtain crystals of the *Rc*Sxph:F-STX complex, *Rc*Sxph was crystallized from solutions containing 33% 2methyl-2,4-pentanediol, 5% PEG 8000, 0.11-0.2 M sodium cacodylate, pH 6.5 and then soaked with F-STX (final concentration, 1 mM) for 5 hours before freezing.

For *Np*Sxph crystallization, protein was purified as described for *Rc*Sxph, except that the final size exclusion chromatography was done using 30 mM NaCl, 10 mM HEPES, pH 7.4. Protein was concentrated to 30-40 mg ml^-1^ using a 50-kDa cutoff Amicon Ultra centrifugal filter unit (Millipore). *Np*Sxph crystals were obtained by hanging drop vapor diffusion at 4°C using 1:1 v/v ratio of protein and precipitant. *Np*Sxph crystals were obtained from 400 nl drops set with Mosquito crystal (Sptlabtech) using 20-25% (v/v) PEG 400, 4-5% (w/v) PGA-LM, 100-200 mM sodium acetate, pH 5.0. For STX co-crystallization, *Np*Sxph and STX (5 mM stock solution prepared in MilliQ water) were mixed in a molar ratio of 1.2:1 STX:*Np*Sxph and incubated on ice for 1 hour before setting up the crystallization trays. For F-STX soaking, *Np*Sxph crystals were soaked with F-STX (final concentration, 1 mM) for 5 hours before freezing. Crystals of the *Np*Sxph:STX complex were grown in the same crystallization solution as *Np*Sxph. *Np*Sxph, *Np*Sxph:STX, and *Np*Sxph:F-STX crystals were harvested and flash-frozen in liquid nitrogen without additional cryoprotectant.

X-ray datasets for *Rc*Sxph mutants, *Rc*Sxph mutant:STX complexes, *Rc*Sxph: F-STX, *Np*Sxph, and *Np*Sxph:STX were collected at 100K at the Advanced Photon Source (APS) beamline 23 ID B of Argonne National Laboratory (Lemont, IL), processed with XDS (*52*) and scaled and merged with Aimless(*53*). *Rc*Sxph structures were determined by molecular replacement of *Rc*Sxph chain B from (PDB: 6O0F) using Phaser from PHENIX (*54*). The resulting electron density map was thereafter improved by rigid body refinement using phenix.refine. The electron density map obtained from rigid body refinement was manually checked and rebuilt in COOT (*55*) and subsequent refinement was performed using phenix.refine.

The *Np*Sxph structure was solved by molecular replacement using the MoRDa pipeline implemented in the Auto-Rikshaw, automated crystal structure determination platform (*56*). The scaled X-ray data and amino-acid sequence of *Np*Sxph were provided as inputs. The molecular replacement search model was identified using the MoRDa domain database derived from the Protein Data Bank (PDB). The MR solution was refined with REFMAC5 (*57*), density modification was performed using PIRATE(*58, 59*), and was followed by the automated model building in BUCCANEER (*60, 61*). The partial model was further refined using REFMAC5 and phenix.refine. Dual fragment phasing was performed using OASIS-2006 (*59*) based on the automatically refined model, and the resulting phases were further improved in PIRATE. The next round of model building was continued in ARP/wARP (*62*)and the resulting structure was refined in REFMAC5. The final model generated in Auto-Rikshaw (720 out of 825 residues built, and 625 residues automatically docked) was further used as a MR search model in Phaser from PHENIX (*54*). The quality of the electron density maps allowed an unambiguous assignment of most of the amino acid residues with the exception of the loop regions and the C2 subdomain showing poor electron density. The apo-*Np*Sxph structure was completed by manual model building in COOT (*55*)and multiple rounds of refinement in phenix.refine. The *Np*Sxph:STX: structure was solved by molecular replacement using the *Np*Sxph structure as a search model in Phaser from PHENIX (*54*). After multiple cycles of manual model rebuilding in COOT(*55*), iterative refinement was performed using phenix.refine. The quality of all models was assessed using MolProbity (*63*) and refinement statistics.

### RNA sequencing of O. sylvatica, D. tinctorius, R. imitator, E. tricolor, A. femoralis, and M. aurantiaca Sxphs

Nearly all poison frog species were bred in the O’Connell Lab or purchased from the pet trade (Josh’s Frogs) except for *O. sylvatica*, which was field collected as described in (*64*). *De novo* transcriptomes for *O. sylvatica*, *D. tinctorius*, *R. imitator*, *E. tricolor*, *A. femoralis*, and *M. aurantiaca* were constructed using different tissue combinations depending on the species. RNA extraction from tissues was performed using TRIzol™ Reagent (Thermo Fisher Scientific). Poly-adenylated RNA was isolated using the NEXTflex PolyA Bead kit (Bioo Scientific, Austin, USA) following manufacturer’s instructions. RNA quality and lack of ribosomal RNA was confirmed using an Agilent 2100 Bioanalyzer or Tapestation (Agilent Technologies, Santa Clara, USA). Each RNA sequencing library was prepared using the NEXTflex Rapid RNAseq kit (Bioo Scientific). Libraries were quantified with quantitative PCR (NEBnext Library quantification kit, New England Biolabs, Ipswich, USA) and an Agilent Bioanalyzer High Sensitivity DNA chip, according to manufacturer’s instructions. All libraries were pooled at equimolar amounts and were sequenced on four lanes of an Illumina HiSeq 4000 machine to obtain 150 bp paired-end reads. *De novo* transcriptomes were assembled using Trinity and once assembled were used to create a BLAST nucleotide database using the BLAST+ command line utilities. The amino acid Sxph sequence of *L. catesbeiana* was used as a query to tBLASTN against the reference transcriptome databases. The Sxph sequence for *O. sylvatica* was lacking the 5’ and 3’ ends, whose sequence was obtained using RACE as described above. After obtaining a full-length sequence, the top BLAST hits from each poison frog transcriptome were manually inspected and aligned to the *O. sylvatica* nucleotide sequence to find full sequences with high similarity. Either a single Sxph sequence from each transcriptome was found to be the best match, or there were multiple transcripts that aligned well, in which case a consensus alignment was created. The largest ORF from each species sequence was translated to create an amino acid sequence for alignment. For the *D. tinctorius*, *R. imitator*, and *A. femoralis* sequences, regions covering the STX binding site and transferrin-related iron-binding sites were confirmed by PCR and sanger sequencing.

### Identification of P. terribilis, R. marina, B. bufo, and B. gargarizans Sxphs

All *P. terribilis* frogs were captive bred in the O’Connell lab poison frog colony. All were sexually mature individuals housed in 18×18×18-inch glass terraria, brought up on a diet of *Drosophila melanogaster* without additional toxins. Frogs were euthanized according to the laboratory collection protocol detailed by Fischer et al. 2019 and tissues were stored in RNALater. Eye tissue was rinsed in PBS before being placed into the beadbug tubes (Sigma-Aldrich, Z763756) prefilled with 1 mL TRIzol (Thermo Fisher Scientific, 15596018) and then RNA was extracted following manufacturer instructions. RNA was reverse transcribed into cDNA following the protocol outlined in Invitrogen’s SuperScript IV Control Reactions First-Strand cDNA Synthesis reaction (Pub. no. MAN0013442, 16 Rev. B). After reverse transcription, cDNA concentration was checked via NanoDrop (Thermo Scientific, ND-ONE-W), and then aliquoted and stored at −20°C until used for PCR. Saxiphilin was amplified from cDNA from *P. terribilis* in 50 µL polymerase chain reactions following the New England Biolabs protocol for Phusion® High-Fidelity PCR Master Mix with HF Buffer (30) (included DMSO). Each reaction was performed with 1 µL of cDNA. PCR primers were designed based on a *O. sylvatica* saxiphilin cDNA sequence previously generated by the O’Connell lab. PCR products were cleaned up using the Thermo Scientific GeneJET Gel Extraction and DNA Cleanup Micro Kit (Catalog number K0832) dimer removal protocol, and then sent out for Sanger Sequencing via the GeneWiz “Premix” service. The segments from sequencing were aligned and assembled but found that the 5’ and 3’ ends of the Sxph sequence for *P. terribilis* were missing, thus the 5’ and 3’ end sequences were subsequently obtained using RACE. 5’ and 3’-RACE-Ready cDNA templates were synthesized using a SMARTer® RACE 5’/3’ Kit (Takara Bio, USA) and subsequently used to amplify 5’ and 3’ end sequences of *P. terribilis* Sxph using internal gene specific primers.

Initial Sxph sequence for *R. marina* was obtained from the genome by searching the draft Cane Toad genome (*65*) with tBLASTN using the *L. catesbeiana* Sxph amino acid sequence as a query. Matching segments from the genome were pieced together to produce an amino acid sequence, however, this sequence was missing part of the 3’ end. To obtain the 3’ residues, the nucleotide sequences from the genome were used to design primers for 3’ Rapid Amplification of cDNA Ends (RACE). One *R. marina* individual from a lab-housed colony was thus euthanized in accordance with UCSF IACUC protocol AN136799, and a portion of the liver was harvested for total RNA extraction using TRIzol™ Reagent (Thermo Fisher Scientific). Total RNA integrity was assessed on a denaturing formaldehyde agarose gel. 3’-RACE-Ready cDNA template was synthesized using a SMARTer® RACE 5’/3’ Kit (Takara Bio, USA) and subsequently used to amplify 3’ end sequences of *R. marina* Sxph using internal gene specific primers designed from *R. marina* genomic sequences. 3’ end sequences of *R. marina* Sxph were determined by gel extraction using QIAquick Gel Extraction Kit (QIAGEN) and verified by sanger sequencing.

Sequences for *Bufo bufo* (Common Toad) and *Bufo gargarizans* (Asiatic toad) Sxphs were identified as sequence searches (tBLASTN) using the *Rm*Sph sequence as a query.

### Two-electrode voltage clamp electrophysiology

Two-electrode voltage-clamp (TEVC) recordings were performed on defolliculated stage V–VI *Xenopus laevis* oocytes harvested under UCSF-IACUC protocol AN178461. Capped mRNA for *P. terribilis (Pt)* Na_V_1.4 (GenBank: MZ545381.1) expressed in a pCDNA3.1 vector (*9*) was made using the mMACHINE™ T7 Transcription Kit (Invitrogen). *Xenopus* oocytes were injected with 3– 6 ng of *Pt* Na_V_1.4 and TEVC experiments were performed 1–2 days post-injection. Data were acquired using a GeneClamp 500B amplifier (MDS Analytical Technologies) controlled by pClamp software (Molecular Devices), and digitized at 1 kHz using Digidata 1332A digitizer (MDS Analytical Technologies).

Oocytes were impaled with borosilicate recording microelectrodes (0.3–3.0 MΩ resistance) backfilled with 3 M KCl. Sodium currents were recorded using a bath solution containing the following, in millimolar: 96, NaCl; 1, CaCl_2_; 1, MgCl_2_; 2, KCl; and 5, HEPES (pH 7.5 with NaOH), supplemented with antibiotics (50 μg ml^− 1^ gentamycin, 100 IU ml^− 1^ penicillin and 100 μg ml^− 1^ streptomycin) and 2.5 mM sodium pyruvate. Sxph responses were measured using Sxph or Sxph mutants purified as described above. Following recording of channel behavior in the absence of toxin, 100 nM STX was applied to achieve ∼90% block. Sxph was then added directly to a 1 mL recording chamber containing the toxin to the desired concentration. For all [Sxph]:[STX] ratios, the concentration of the stock Sxph solution added to the chamber was adjusted so that the volume of the added Sxph solution was less than 1% of the total volume of the recording chamber. All toxin effects were assessed with 60-ms depolarization steps from −120 to 0 mV with a holding potential of −120 mV and a sweep-to-sweep duration of 10 s. Recordings were conducted at room temperature (23 ± 2 °C). Leak currents were subtracted using a P/4 protocol during data acquisition. Data Analysis was performed using Clampfit 10.6 (Axon Instruments) and SigmaPlot (Systat Software).

### F-STX synthesis

All reagents were obtained commercially unless otherwise noted. *N,N*-Dimethylformamide (DMF) was passed through two columns of activated alumina prior to use. High-performance liquid chromatography-grade CH_3_CN and H_2_O were obtained from commercial suppliers. Semi-preparative high-performance liquid chromatography (HPLC) was performed on a Varian ProStar model 210. A high-resolution mass spectrum of F-STX was obtained from the Vincent Coates Foundation Mass Spectrometry Laboratory at Stanford University. The sample was analyzed with HESI-MS by direct injection onto Waters Acquity UPLC and a Thermo Fisher Orbitrap Exploris^TM^ 240 mass spectrometer scanning m/z 100–1000. F-STX was quantified by ^1^H NMR spectroscopy on a Varian Inova 600 MHz NMR instrument using distilled DMF as an internal standard. A relaxation delay (d1) of 20 s and an acquisition time (at) of 10 s were used for spectral acquisition. The concentration of F-STX was determined by integration of ^1^H signals corresponding to F-STX and a fixed concentration of the DMF standard.

To an ice-cold solution of saxitoxin-N21-hexylamine (1.4 µmol) in 140 µL of pH 9.5 aqueous bicarbonate buffer (0.2 M aqueous NaHCO_3_, adjusted to pH 9.5 with 1 M aqueous NaOH) was added a solution of fluorescein NHS-ester, 6-isomer (2.0 mg, 4.2 μmol, 3.0 equiv, Lumiprobe) in 140 µL of DMSO. The reaction flask was stoppered, wrapped in foil, and placed in a sonication bath for 30 s. The reaction mixture was then stirred at room temperature for 4 h. Following this time, the reaction was quenched by the addition of 0.3 mL of 1% aqueous CF_3_CO_2_H. The reaction mixture was diluted with 1.1 mL of 10 mM aqueous CF_3_CO_2_H and 0.3 mL of DMSO and filtered through a VWR 0.22 µm PTFE filter. The product was purified by reverse-phase HPLC (Silicycle SiliaChrom dt C18, 5 µm, 10 x 250 mm column, eluting with a gradient flow of 10→40% CH_3_CN in 10 mM aqueous CF_3_CO_2_H over 40 min, 214 nm UV detection). At a flow rate of 4 mL/min, F-STX had retention time of 31.00 min and was isolated as a dark yellow powder following lyophilization (1.08 µmol, 77%, ^1^H NMR quantitation).

^1^H NMR (600 MHz, D_2_O) δ 8.05 (d, *J* = 8.1 Hz, 1H), 7.94 (d, *J* = 8.9 Hz, 1H), 7.48 (s, 1H), 6.95 (d, *J* = 9.0 Hz, 2H), 6.79 (s, 2H), 6.67 (dt, *J* = 9.1, 2.2 Hz, 2H), 4.60 (d, *J* = 1.2 Hz, 1H), 4.09–4.05 (m, 1H), 3.89 (dd, *J* = 11.6, 5.2 Hz, 1 H), 3.70 (dt, *J* = 10.1, 5.5 Hz, 1H), 3.64 (dd, *J* = 8.7, 5.4 Hz, 1H), 3.47–3.42 (m, 1H), 3.27 (t, *J* = 6.6 Hz, 2 H), 2.97–2.89 (m, 2H), 2.36–2.33 (m 1H), 2.30–2.24 (m, 1H), 1.48–1.45 (m, 2H), 1.32–1.29 (m, 2H), 1.25–1.21 (m, 4H) ppm. HRMS (ESI^+^) calcd for C_37_H_41_N_8_O_10_, 757.2940; found 757.2918 (M^+^).

## Supplementary material

Supplementary material containing Figures S1-S11, Movies M1-M4, and Tables S1-S2.

## Supplementary material

### Supplementary material inventory

**Movie M1 *Rc*Sxph-Y558A conformational changes upon STX binding.**

**Movie M2 *Rc*Sxph-Y558I conformational changes upon STX binding.**

**Movie M3 Conformational changes between *R*cSxph and *Np*Sxph.**

**Movie M4 *Np*Sxph conformational changes upon STX binding.**

---

**Figure S1.**
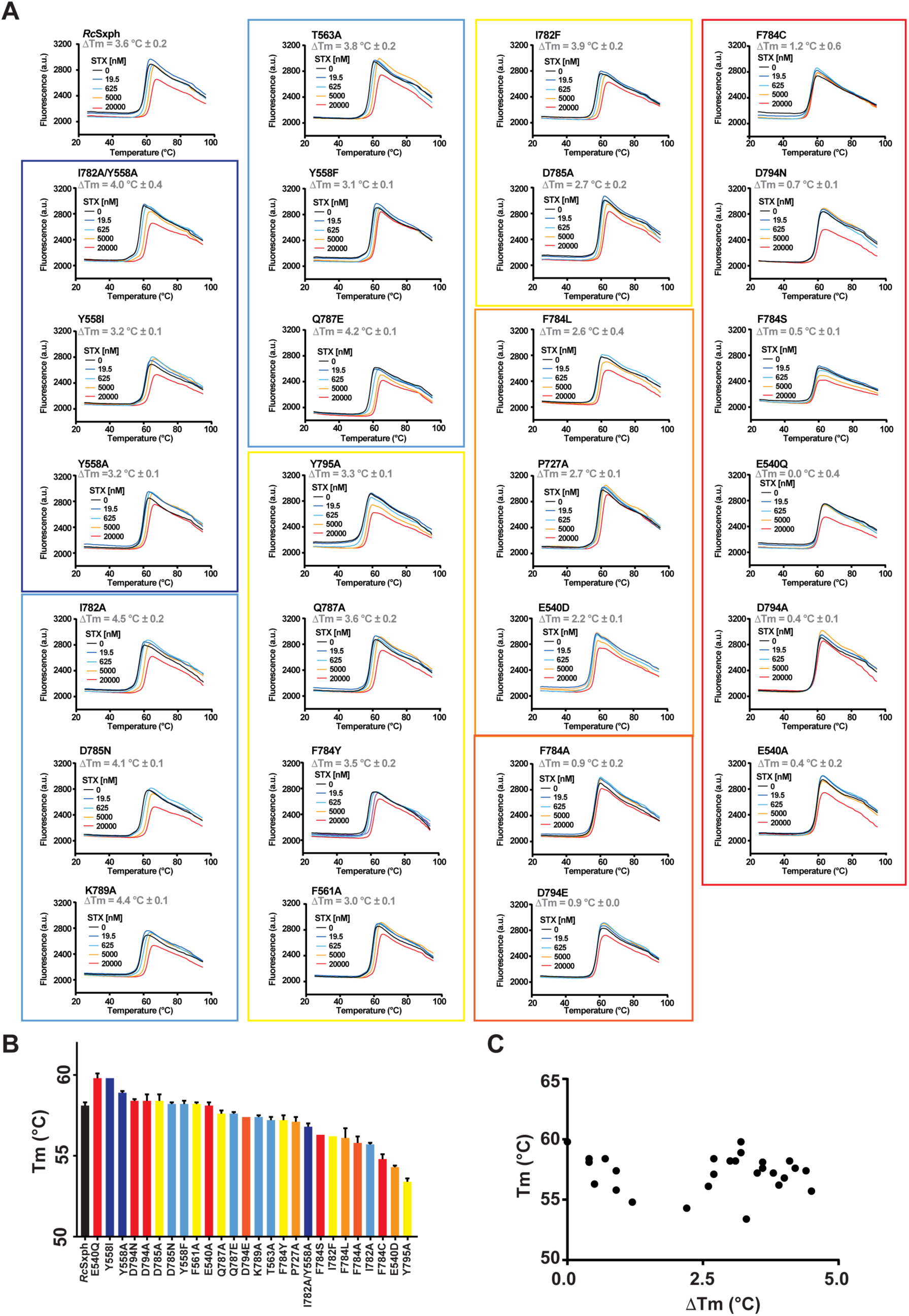
*Rc*Sxph thermofluor (TF) assay. **A,** Exemplar thermofluor (TF) assay results for *Rc*Sxph in the presence of the indicated concentrations of STX. Curves for *Rc*Sxph, E540A, P727A, Y558A, F561A, and T563A are identical to those shown in Figs. 1A and 1B. ΔTm values are indicated. **B,** Baseline Tm values for *Rc*Sxph and the indicated mutants. **C,** Plot of Tm vs. ΔTm for the proteins in ‘**B**’. Colored boxes in ‘A’ and bars in ‘**B**’ correspond to ΔΔG classifications in Table 1. Error bars are S.E.M.

**Figure S2.**
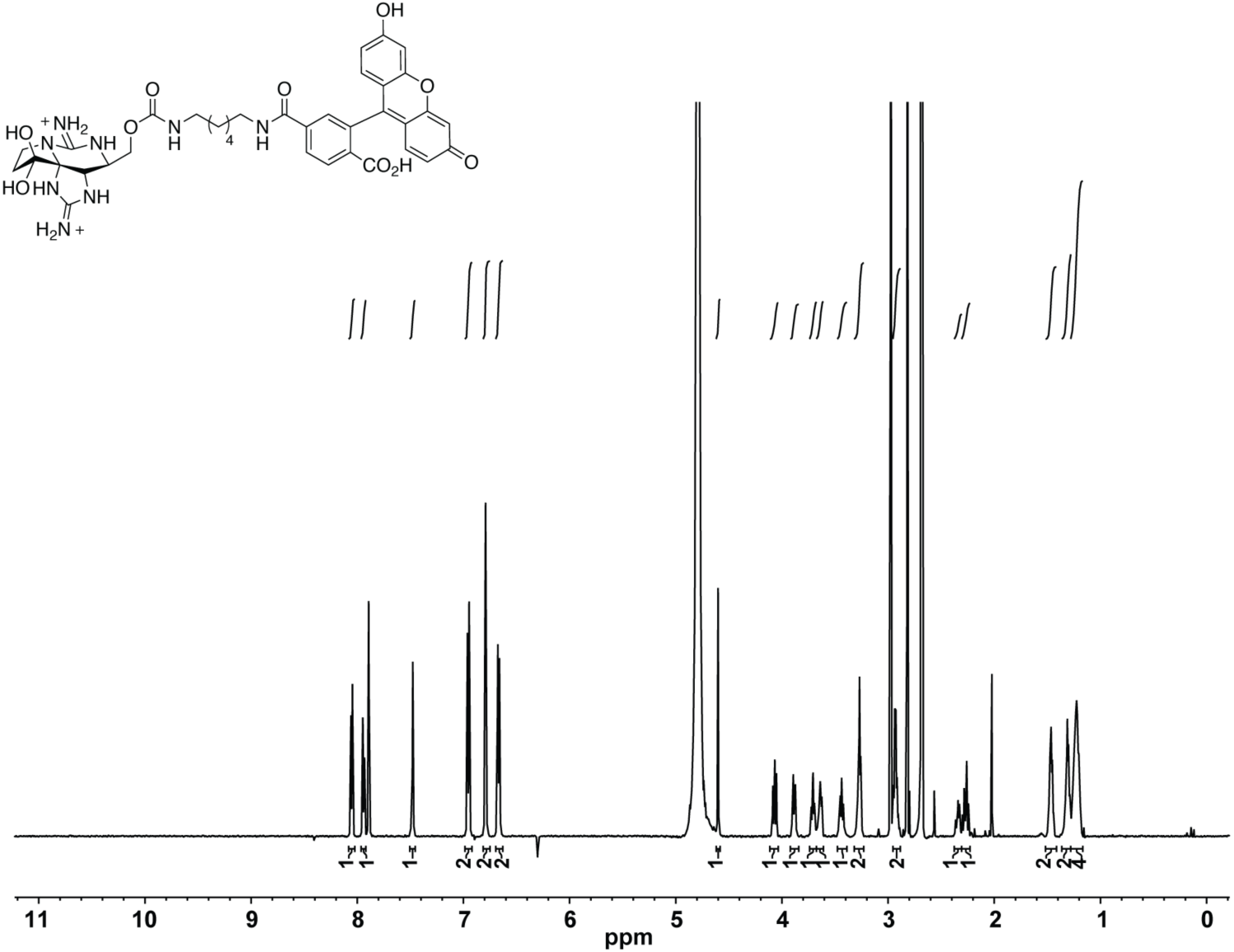
F-STX NMR spectrum. ^1^H NMR (600 MHz, D_2_O) δ 8.05 (d, *J* = 8.1 Hz, 1H), 7.94 (d, *J* = 8.9 Hz, 1H), 7.48 (s, 1H), 6.95 (d, *J* = 9.0 Hz, 2H), 6.79 (s, 2H), 6.67 (dt, *J* = 9.1, 2.2 Hz, 2H), 4.60 (d, *J* = 1.2 Hz, 1H), 4.09–4.05 (m, 1H), 3.89 (dd, *J* = 11.6, 5.2 Hz, 1 H), 3.70 (dt, *J* = 10.1, 5.5 Hz, 1H), 3.64 (dd, *J* = 8.7, 5.4 Hz, 1H), 3.47–3.42 (m, 1H), 3.27 (t, *J* = 6.6 Hz, 2 H), 2.97–2.89 (m, 2H), 2.36–2.33 (m 1H), 2.30–2.24 (m, 1H), 1.48–1.45 (m, 2H), 1.32–1.29 (m, 2H), 1.25–1.21 (m, 4H) ppm.

**Figure S3.**
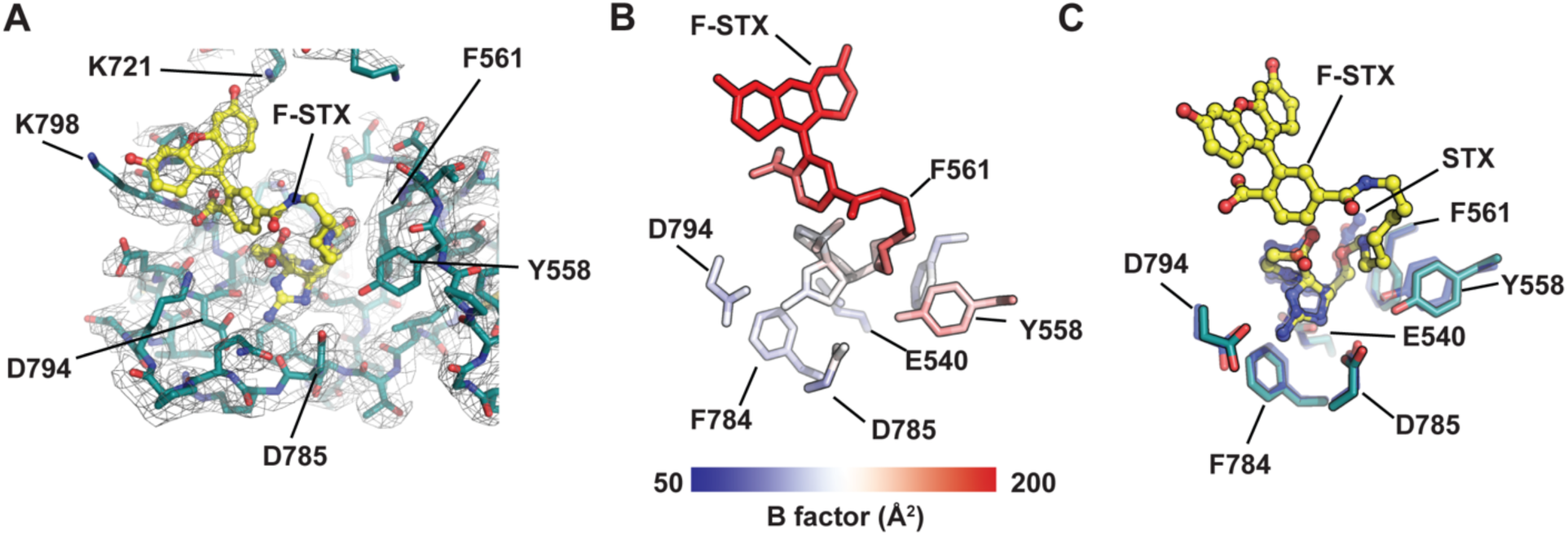
Structure of the *Rc*Sxph:F-STX: complex. **A,** Exemplar electron density (1σ) for *Rc*Sxph (deep teal) and F-STX (yellow). **B,** *Rc*Sxph:F-STX: B-factors for the F-STX ligand and select binding site residues. **C,** Superposition of the STX binding sites of the *Rc*Sxph:F-STX: and *Rc*Sxph:STX (PDB:6O0F) (blue) (*1*) complexes.

**Figure S4.**
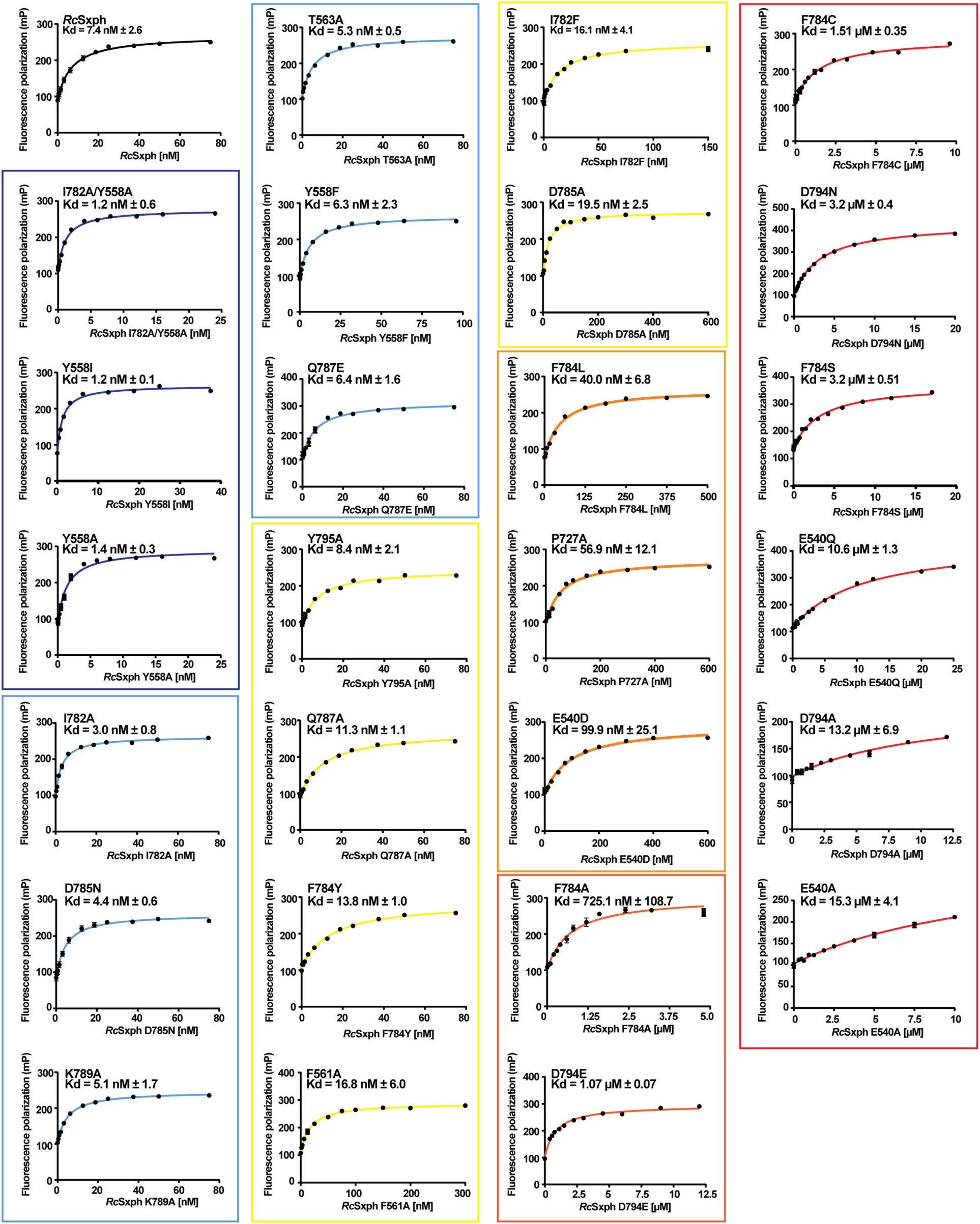
*Rc*Sxph fluorescence polarization (FP) assay. Exemplar FP binding curves and Kds for *Rc*Sxph and the indicated mutants. Curves for *Rc*Sxph, E540A, P727A, Y558A, F561A, and T563A are identical to those shown in Fig. 1D. Colored boxes and lines in correspond to ΔΔG classifications in Table 1. Error bars are S.E.M.

**Figure S5.**
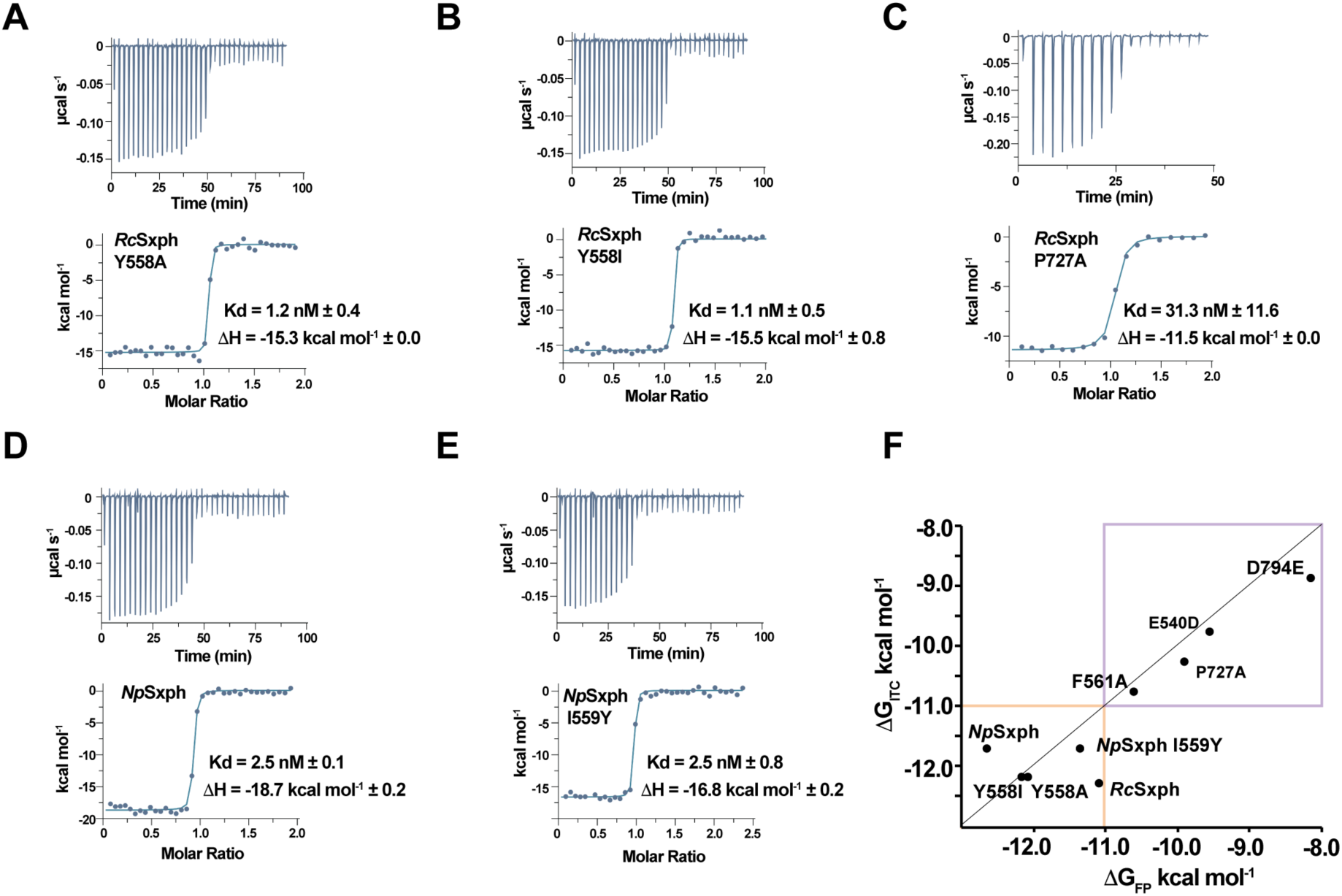
*Rc*Sxph and *Np*Sxph Isothermal titration calorimetry. Exemplar ITC isotherms for **A,** 100 µM STX into 10 µM *Rc*Sxph Y558A, **B,** 100 µM STX into 10 µM *Rc*Sxph Y558I, **C,** 100 µM STX into 10 µM *Rc*Sxph P727A, **D,** 100 µM STX into 9.7 µM *Np*Sxph, and **E,** 100 µM STX into 7.9 µM *Np*Sxph I559Y. **F,** Comparison of ΔG_ITC_ for STX and ΔG_FP_ for F-STX for *Rc*Sxph, *Np*Sxph, and indicated mutants. Purple box highlights region of good correlation. Orange box indicates region outside of the ITC dynamic range. *Rc*Sxph data are identical to Fig. 1G.

**Figure S6.**
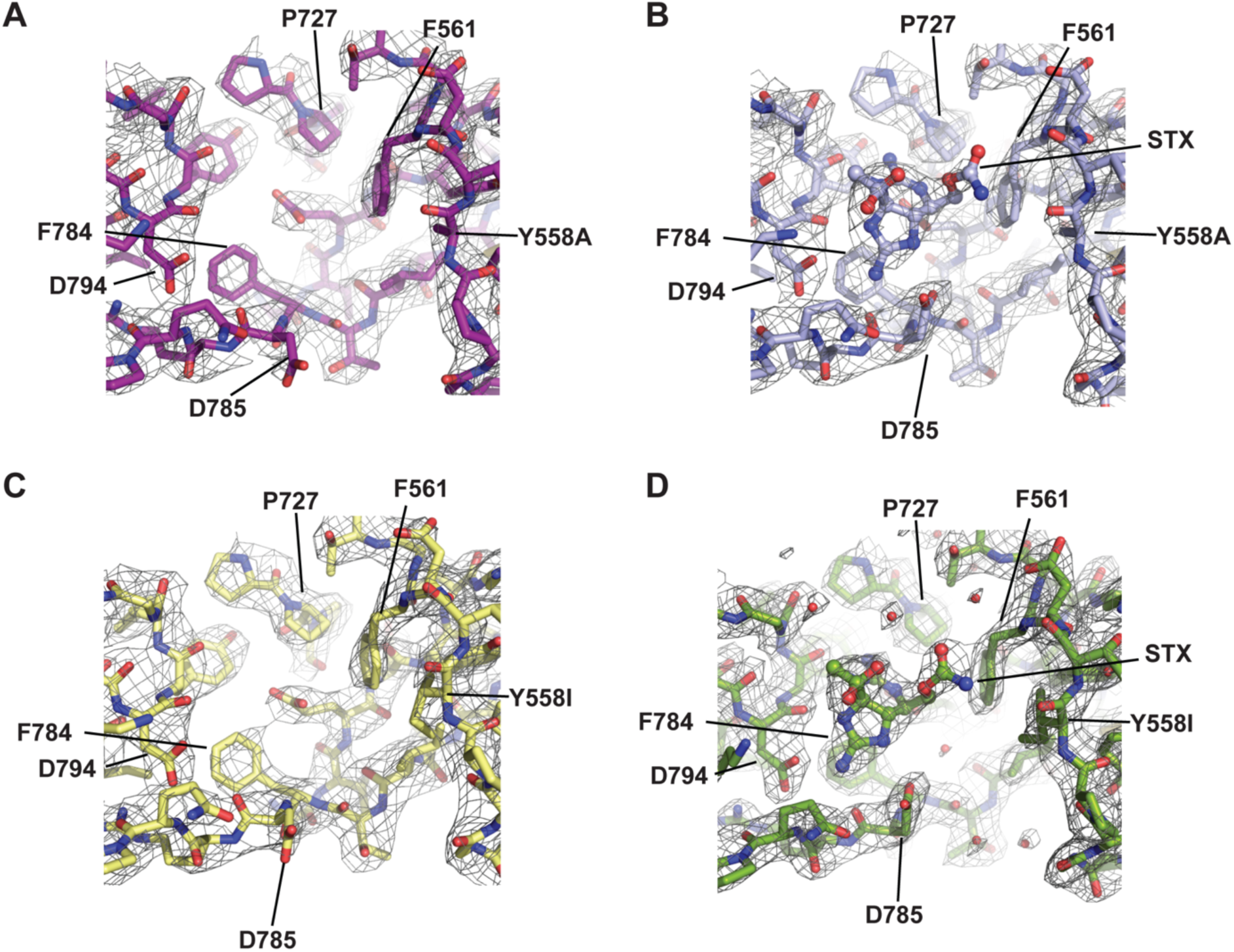
*Rc*Sxph Y558A and *Rc*Sxph-Y558I structures and STX complexes. Exemplar electron density (1.5 σ) for **A,** *Rc*Sxph Y558A (purple), **B,** *Rc*Sxph-Y558A:STX (light blue), **C,** *Rc*Sxph-Y558I (pale yellow), and **D,** *Rc*Sxph-Y558I:STX (splitpea). Select residues and STX are indicated.

**Figure S7.**
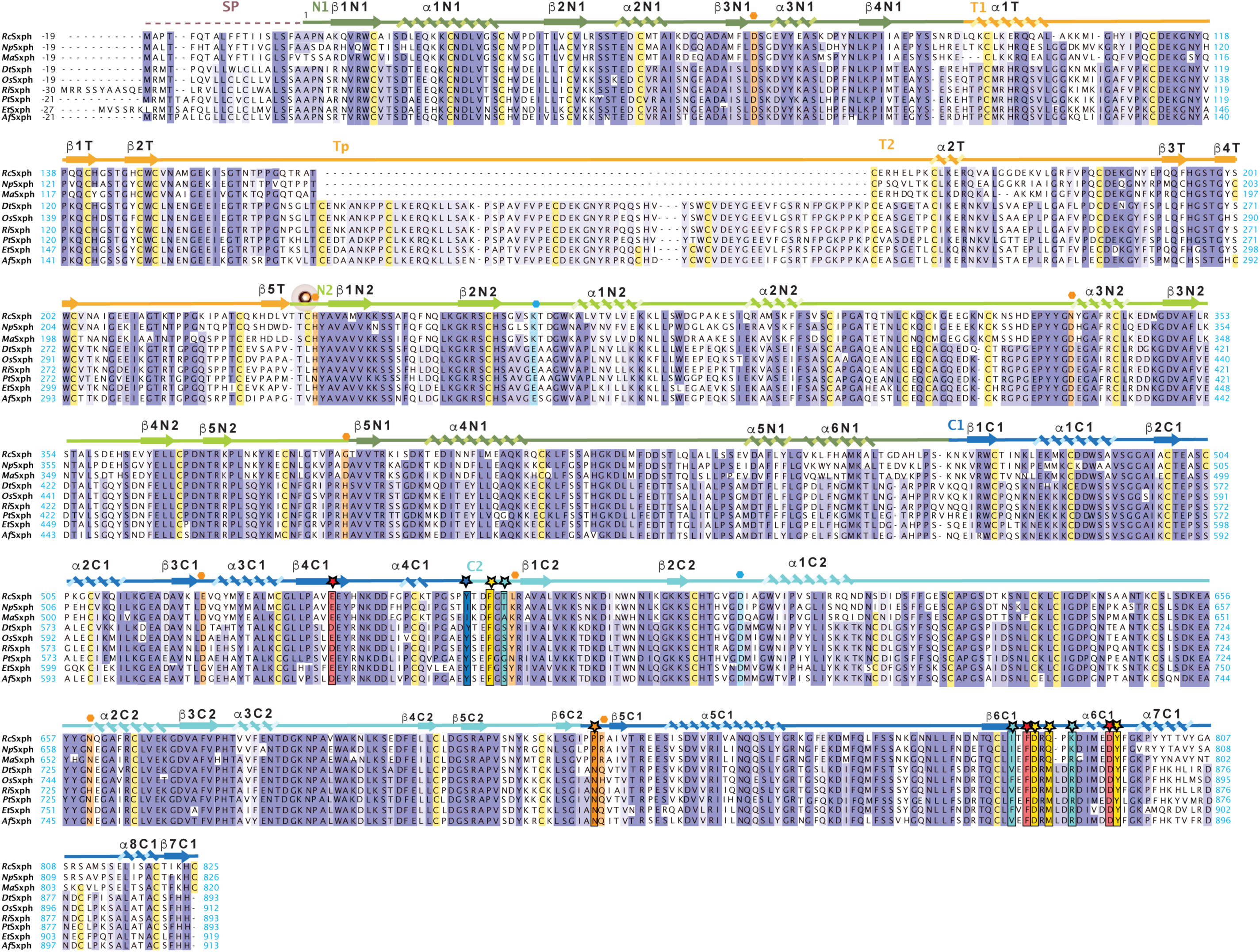
Frog Sxph sequence alignment. Sxph sequence alignment for *Rc*Sxph, *Np*Sxph, *Ma*Sxph, *Dt*Sxph, *Os*Sxph, *Ri*Sxph, *Pt*Sxph, *Et*Sxph, and *Af*Sxph. Domains and secondary structure are from *Rc*Sxph. N1 (dark green), N2 (light green), Thy1 domains (orange), C1 (marine), C2 (cyan). STX binding site residues are indicated by stars and colored based on the alanine scan results in Table 1. Residues corresponding to transferrin Fe^3+^ and carbonate ligands are indicated by orange and blue hexagons, respectively and highlighted (*1, 2*).

**Figure S8.**
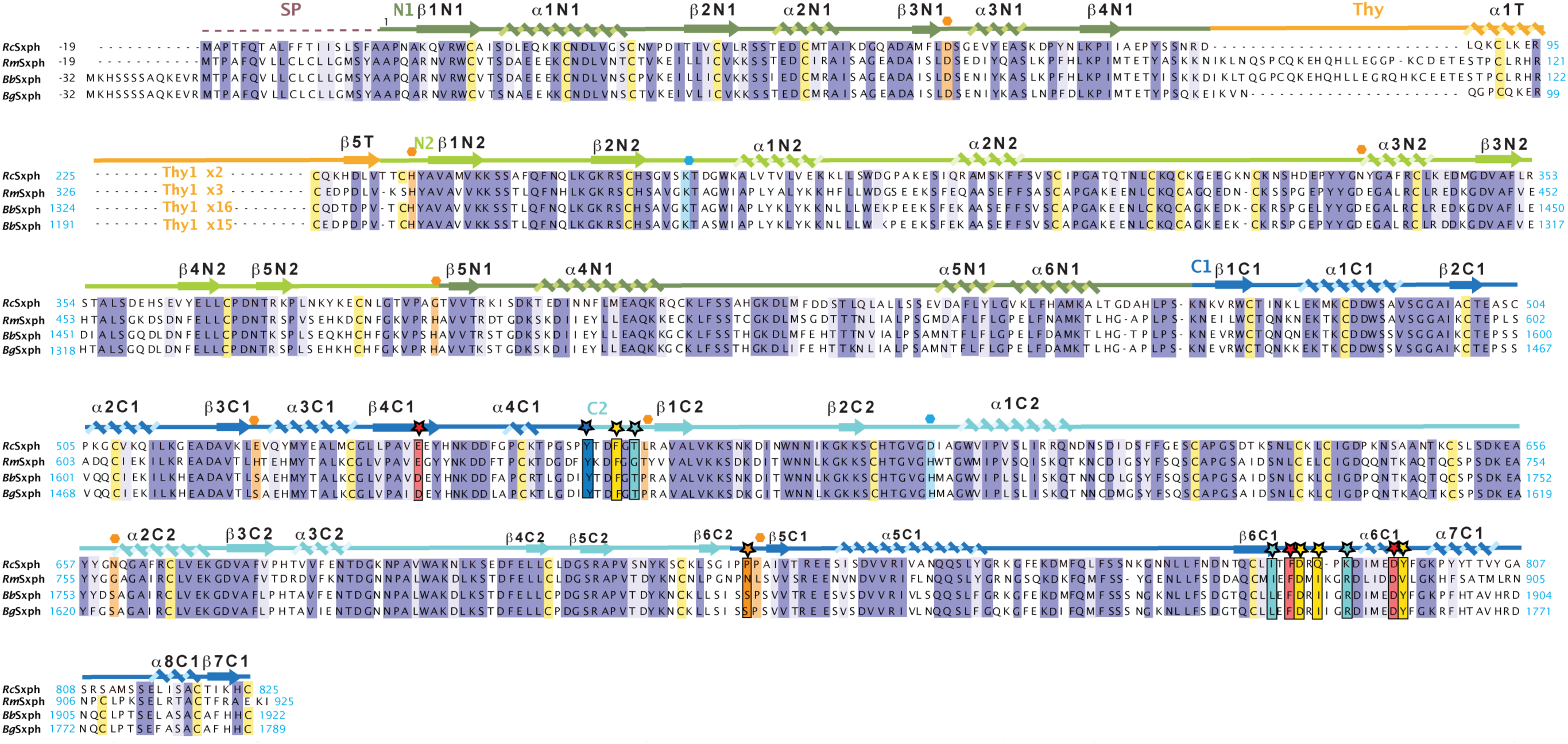
Toad Sxph sequence alignment. Sxph sequence alignment for *Rc*Sxph, and toad saxiphilins *Rm*Sxph, *Bb*Sxph (NCBI:XM_040427746.1), and *Bg*Sxph(NCBI:XP_044148290.1). Domains and secondary structure are from *Rc*Sxph. N1 (dark green), N2 (light green), Thy1 domains (orange), C1 (marine), C2 (cyan). STX binding site residues are indicated by stars and colored based on the alanine scan results in Table 1. Residues corresponding to transferrin Fe^3+^ and carbonate ligands (*1, 2*) are indicated by orange and blue hexagons, respectively and highlighted. Only beginning and ends of the Thy1 domains are shown. Total number of Thy1 domains are indicated.

**Figure S9.**
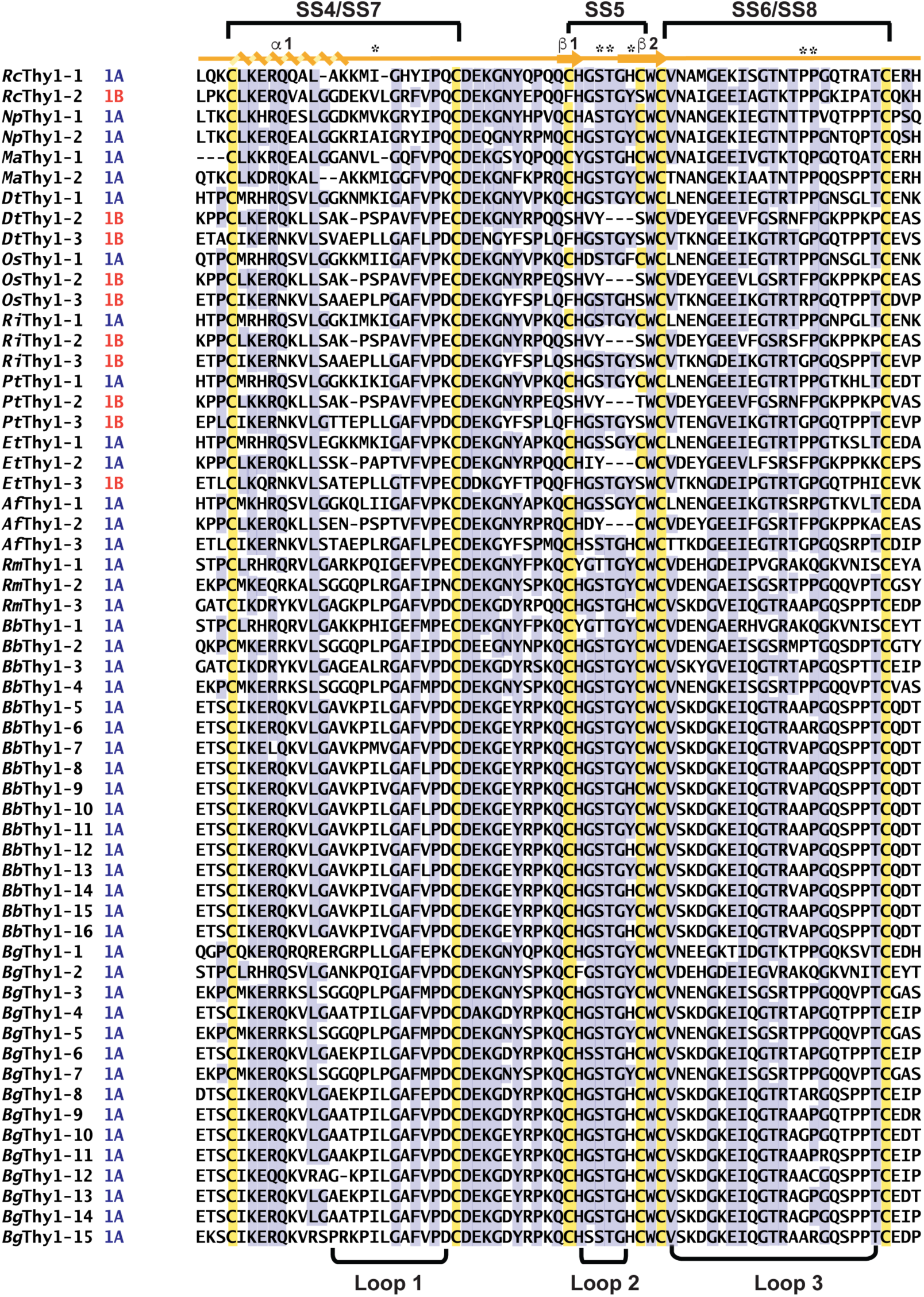
Thy1 domain sequence alignment. Thy1 domains from *Rc*Sxph, *Np*Sxph, *Ma*Sxph, *Dt*Sxph, *Os*Sxph, *Ri*Sxph, *Pt*Sxph, *Et*Sxph, *Af*Sxph, *Rm*Sxph, *Bb*Sxph, and *Bg*Sxph and the type (1A or 1B) are shown. Secondary structure from *Rc*Sxph Thy1-1 is shown. Cysteine are highlighted. SS4-SS8 indicate disulfide numbers from *Rc*Sxph. Loop regions are indicated.

**Figure S10.**
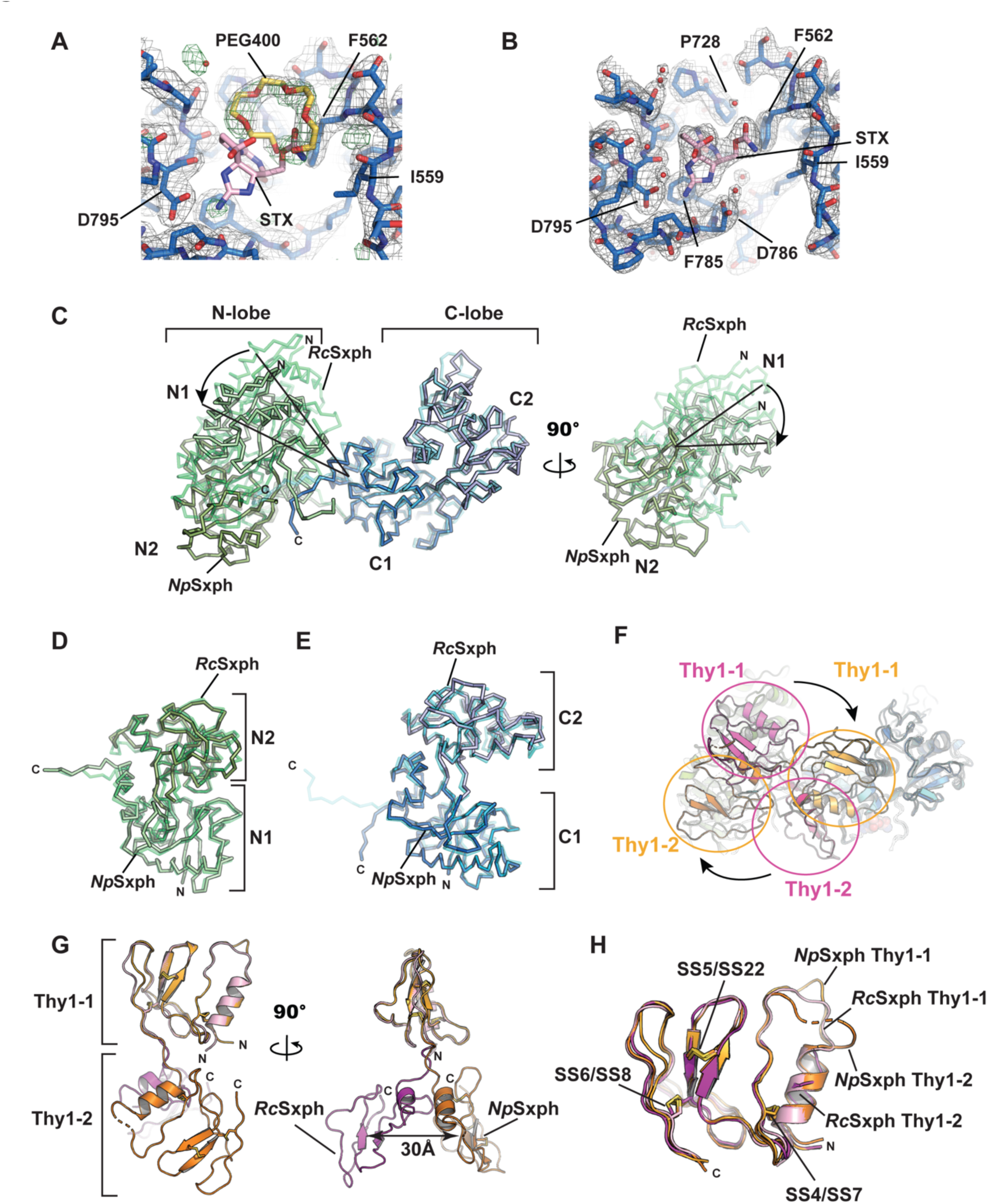
*Np*Sxph structure and comparisons with *Rc*Sxph. **A,** and **B,** Exemplar electron density for **A,** *Np*Sxph (2Fo-Fc, 1.5 σ, grey) and (Fo-Fc, 3.0 σ, green). **B,** *Np*Sxph:STX (2Fo-Fc, 1.5 σ, grey). *N*pSxph (marine), STX (pink), and PEG400 (yellow) are shown. STX (pink) from the *Np*Sxph:STX complex is shown in ‘**A’** to compare with the PEG400 position. Select residues are labelled. **C,** *Np*Sxph and *Rc*Sxph superposition using the C-lobes. N- and C-lobes are green/light green and marine/light blue for *Np*Sxph and *Rc*Sxph, respectively. Arrow indicate relationships between *Np*Sxph and *Rc*Sxph N-lobes. **D,** Superposition of *Np*Sxph (green) and *Rc*Sxph (light green) N-lobes. **E,** Superposition of *Np*Sxph (marine) and *Rc*Sxph (light blue) C-lobes. **F**, Cartoon diagram of *Np*Sxph and *Rc*Sxph superposition from ‘**C**’ showing the change in Thy1 domain positions. *N*pSxph Thy1 domains (orange) and *Rc*Sxph Thy1 domains (magenta) are indicated. **G**, Cartoon diagam of *Np*Sxph and *Rc*Sxph Thy1 domains superposed on Thy1-1. *Np*Sxph and *Rc*Sxph Thy1-1 and Thy1-2 are light orange and pink and orange and magenta, respectively. **H,** Superposition of individual *Np*Sxph and *Rc*Sxph Thy1-1 and Thy1-2 domains. Colors are as in ‘**G**’. Disulfide bonds are indicated.

**Figure S11.**
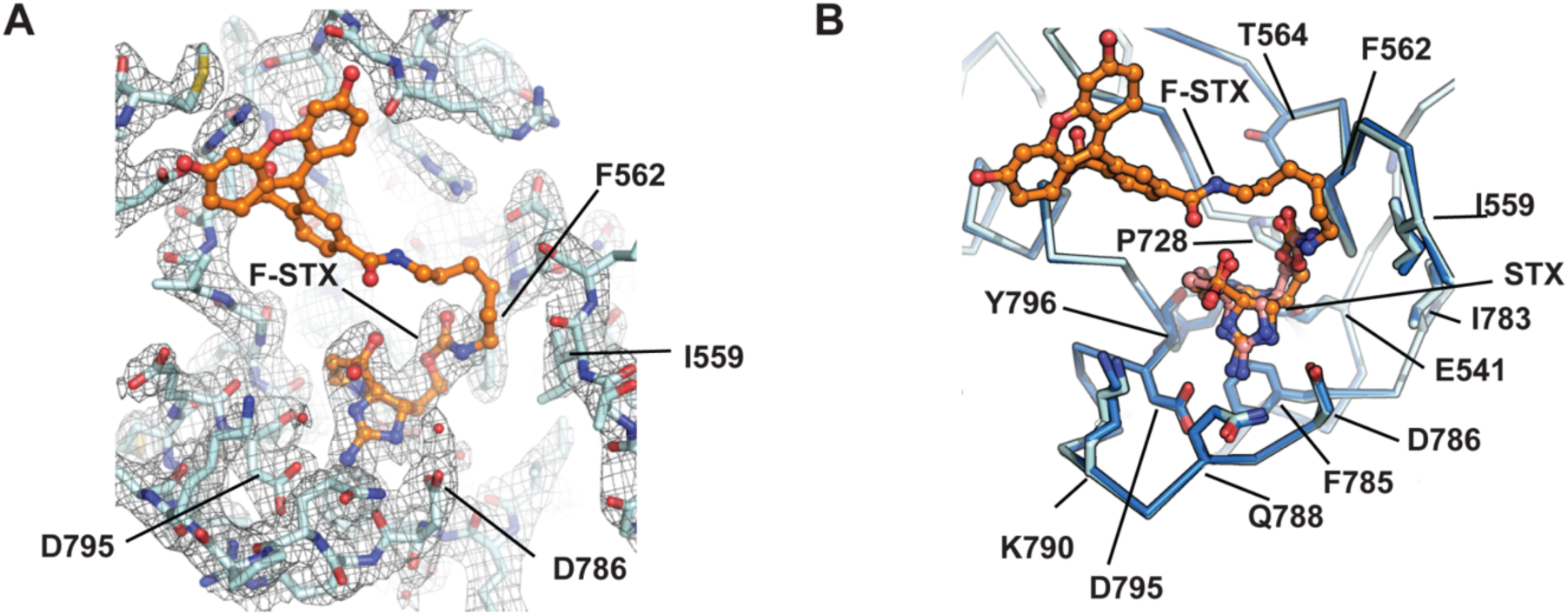
Structure of the *Np*Sxph:F-STX complex. **A,** Exemplar electron density for *Np*Sxph:F-STX(2Fo-Fc, 1.5 σ, grey). *Np*Sxph (cyan) and F-STX (orange). **B,** Comparison of *Np*Sxph:STX (marine) and *Np*Sxph:F-STX STX binding sites. STX from *Np*Sxph is pink. F-STX is orange. Select residues are indicated.

**Movie M1 *Rc*Sxph-Y558A conformational changes upon STX binding.** Morph between the apo-*Rc*Sxph-Y558A and *Rc*Sxph-Y558A:STX structures showing the STX binding pocket. Sidechains are shown as sticks. STX is red.

**Movie M2 *Rc*Sxph-Y558I conformational changes upon STX binding.** Morph between the apo-*Rc*Sxph-Y558I and *Rc*Sxph-Y558A:STX structures showing the STX binding pocket. Sidechains are shown as sticks. STX is red.

**Movie M3 Conformational changes between *R*cSxph and *Np*Sxph.** Morph between apo-*Rc*Sxph (PDB:6O0D) (*1*) (starting structure) and apo-*Np*Sxph (final structure). N-lobe (green), C-lobe (blue), and Thy domains (magenta) are shown. N1, N2, C1, and C2 subdomains and Thy1-1, and Thy1-2 are labeled.

**Movie M4 *Np*Sxph conformational changes upon STX binding.** Morph between the apo-*Np*Sxph and *Np*Sxph:STX structures showing the STX binding pocket. Sidechains are shown as sticks. STX is red.

**Table S1.**
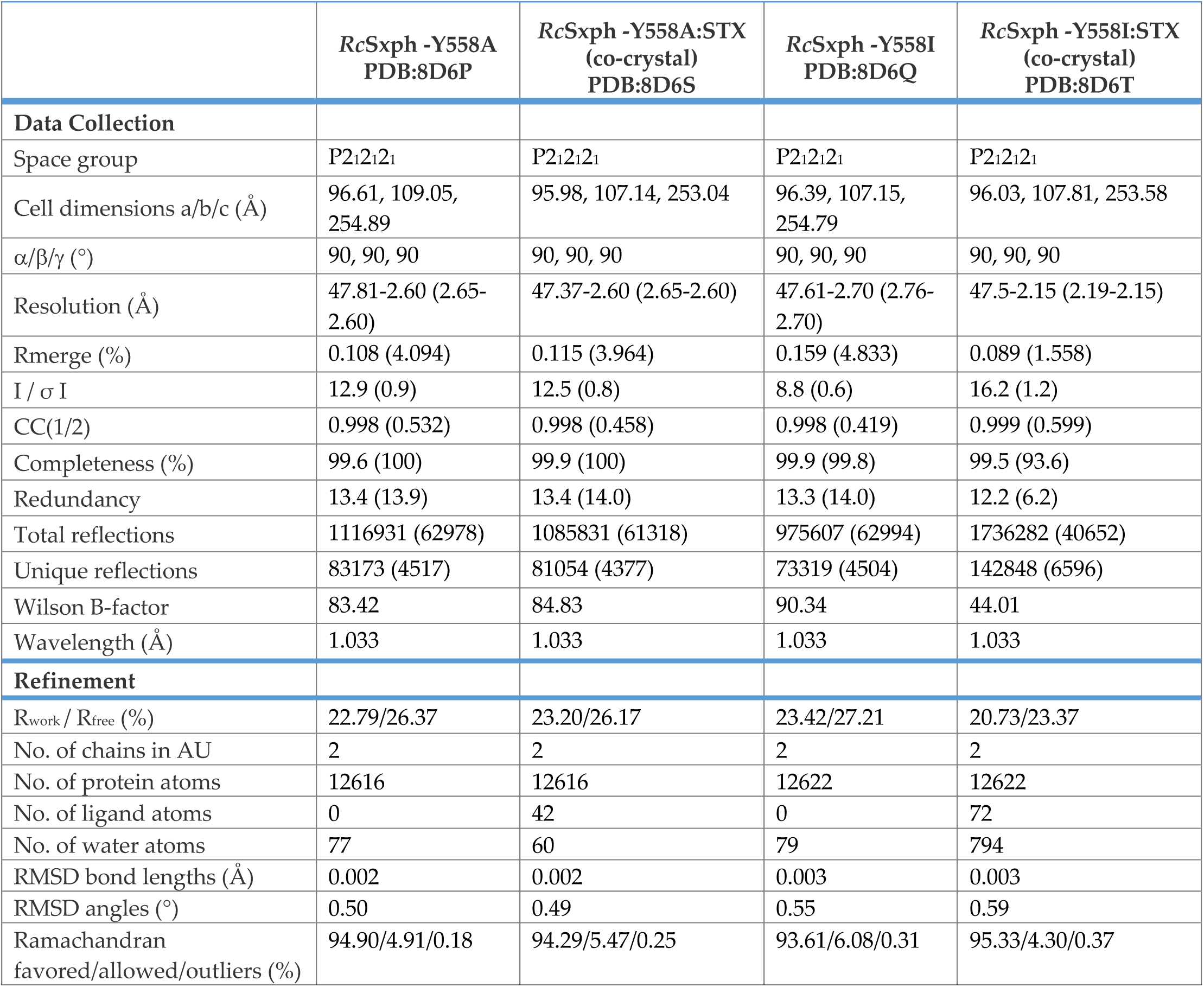
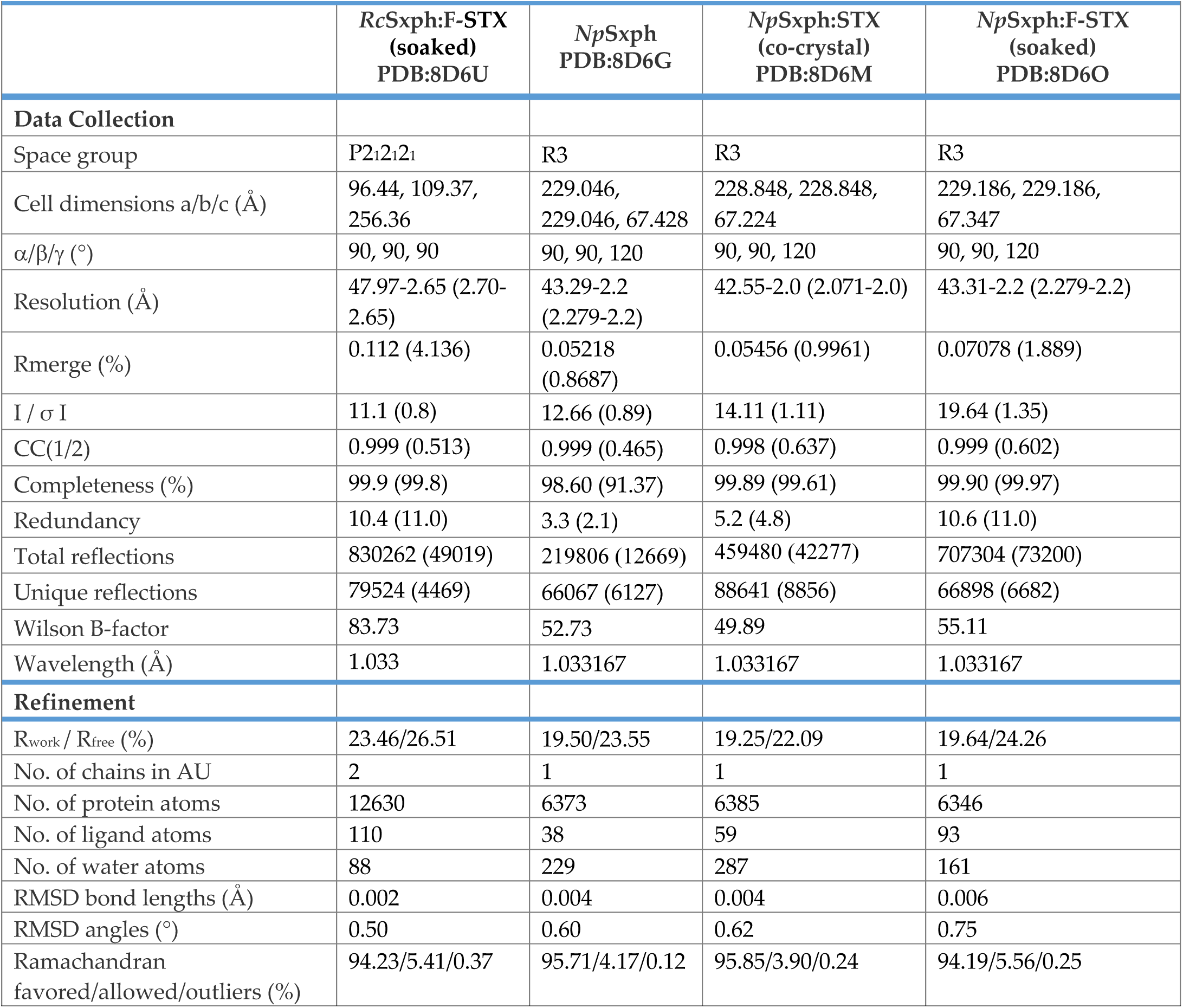
Crystallographic data collection and refinement statistics.

**Table S2.**
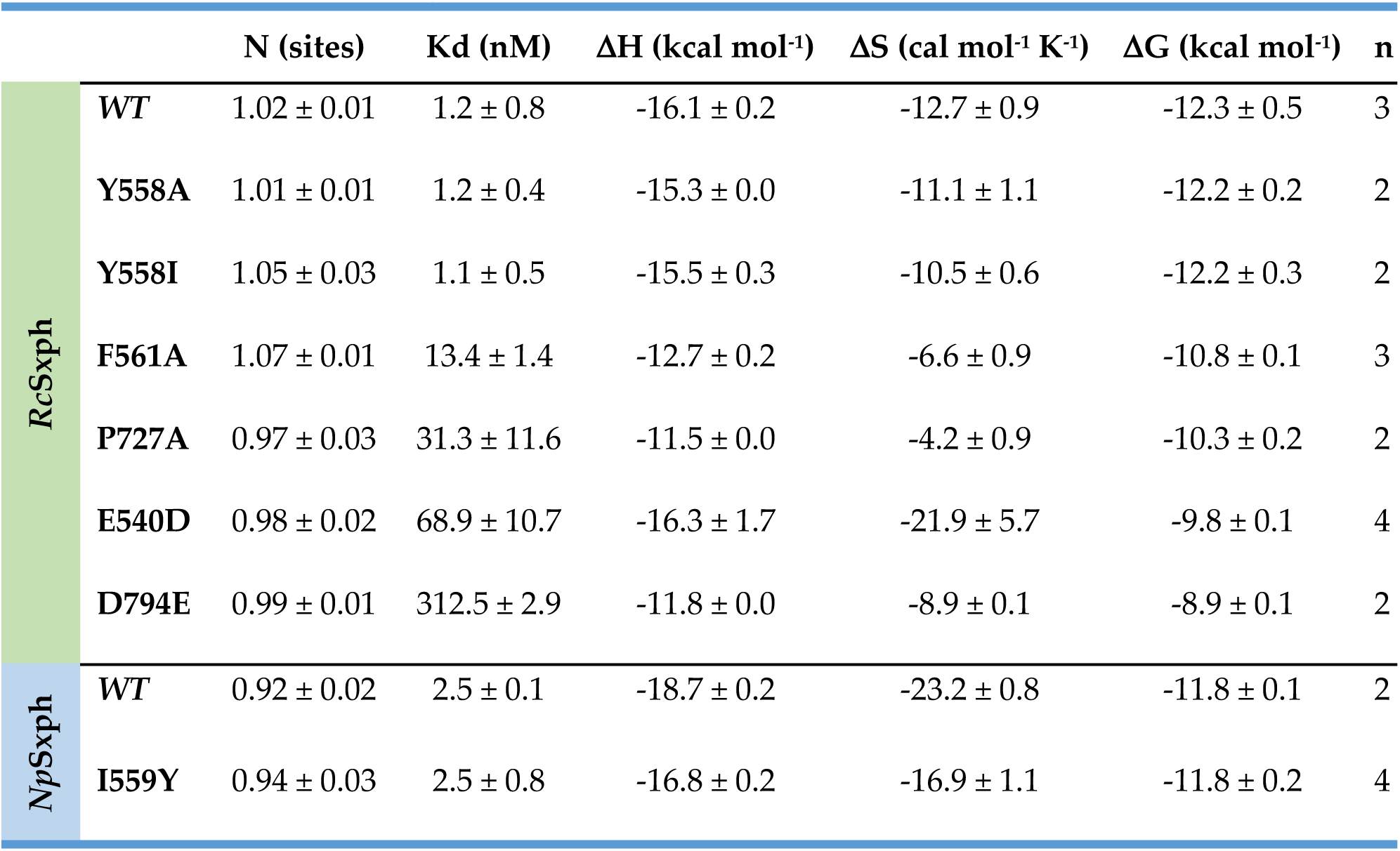
*Rc*Sxph:STX and *Np*Sxph:STX thermodynamic binding parameters.

