## Supplementary Figures and Tables for "Definition of a saxitoxin (STX) binding code enables discovery and characterization of the Anuran saxiphilin family"

8 June 2022

[7]Molecular Biophysics and Integrated Bio-imaging Division
Lawrence Berkeley National Laboratory, Berkeley, CA 94720 USA

[2]Department of Chemistry
Stanford University, Stanford, CA 94305

[3]Department of Biology
Stanford University, Stanford, CA 94305

[†]Equal contributions

Keywords: Saxitoxin, Toxin resistance, Saxiphilin

**Short Title: STX recognition code for saxiphilins**

**Supplementary material inventory.**

**Figure S1 *Rc*Sxph thermofluor (TF) assay.**

**Figure S2 F-STX NMR spectrum.**

**Figure S3 Structure of the *Rc*Sxph:F-STX complex.**

**Figure S4 *Rc*Sxph fluorescence polarization (FP) assay**

**Figure S5 *Rc*Sxph and *Np*Sxph Isothermal titration calorimetry.**

**Figure S6 *Rc*Sxph Y558A and *Rc*Sxph-Y558I structures and STX complexes.**

**Figure S7 Frog Sxph sequence alignment.**

**Figure S8 Toad Sxph sequence alignment.**

**Figure S9 Thy1 domain sequence alignment.**

**Figure S10 *Np*Sxph structure and comparisons with *Rc*Sxph.**

**Figure S11 Structure of the *Np*Sxph:F-STX complex.**

**Movie M1 *Rc*Sxph-Y558A conformational changes upon STX binding.**

**Movie M2 *Rc*Sxph-Y558I conformational changes upon STX binding.**

**Movie M3 Conformational changes between *R*cSxph and *Np*Sxph.**

**Movie M4 *Np*Sxph conformational changes upon STX binding.**

**Table S1 Crystallographic data collection and refinement statistics**

**Table S2 *Rc*Sxph:STX and *Np*Sxph:STX thermodynamic binding parameters.**

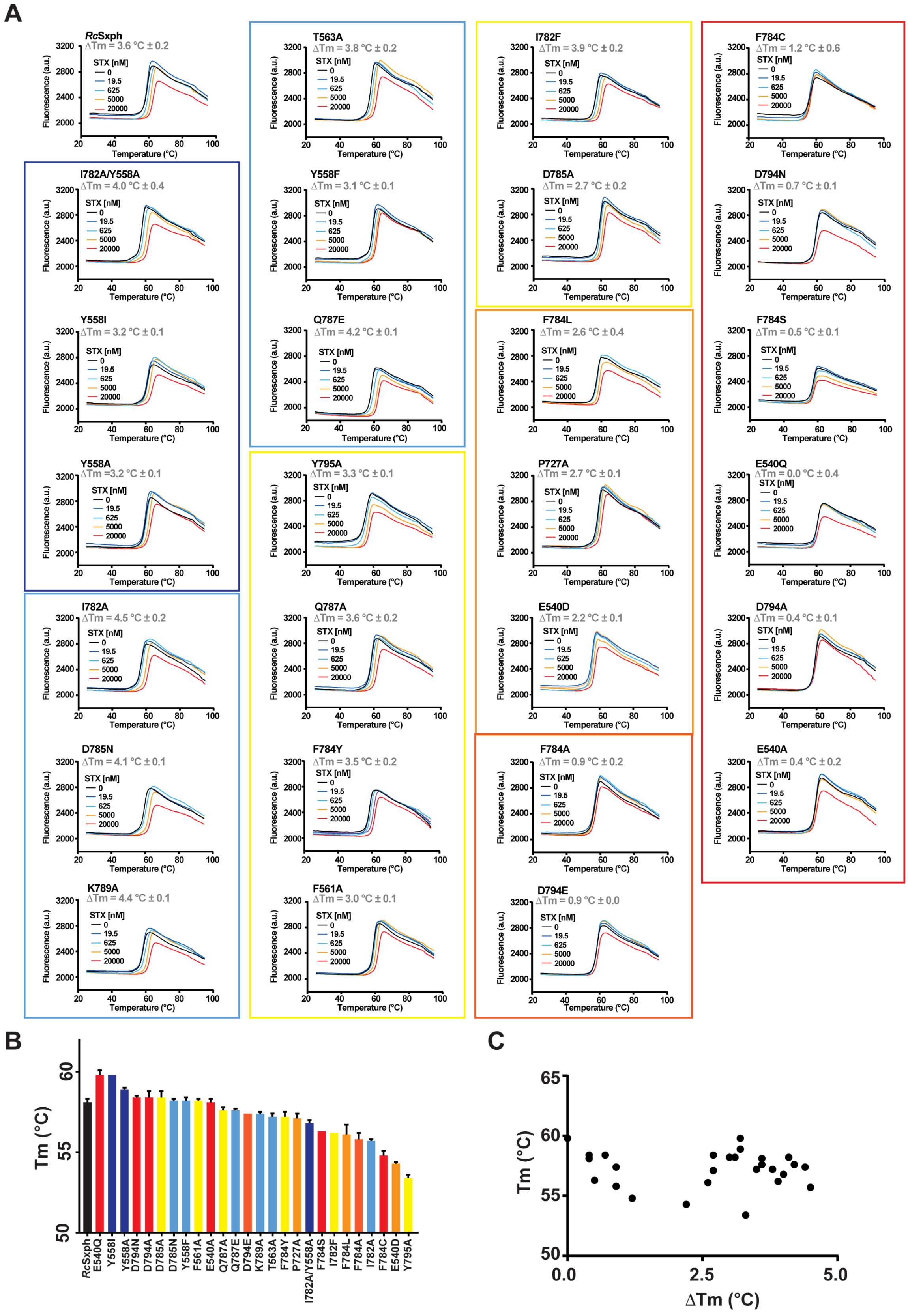
A
RcSxph ΔTm = 3.6 °C ± 0.2
T563A ΔTm = 3.8 °C ± 0.2
I782F ΔTm = 3.9 °C ± 0.2
F784C ΔTm = 1.2 °C ± 0.6
I782A/Y558A ΔTm = 4.0 °C ± 0.4
Y558F ΔTm = 3.1 °C ± 0.1
D785A ΔTm = 2.7 °C ± 0.2
D794N ΔTm = 0.7 °C ± 0.1
Y558I ΔTm = 3.2 °C ± 0.1
Q787E ΔTm = 4.2 °C ± 0.1
F784L ΔTm = 2.6 °C ± 0.4
F784S ΔTm = 0.5 °C ± 0.1
Y558A ΔTm =3.2 °C ± 0.1
Y795A ΔTm = 3.3 °C ± 0.1
P727A ΔTm = 2.7 °C ± 0.1
E540Q ΔTm = 0.0 °C ± 0.4
I782A ΔTm = 4.5 °C ± 0.2
Q787A ΔTm = 3.6 °C ± 0.2
E540D ΔTm = 2.2 °C ± 0.1
D794A ΔTm = 0.4 °C ± 0.1
D785N ΔTm = 4.1 °C ± 0.1
F784Y ΔTm = 3.5 °C ± 0.2
F784A ΔTm = 0.9 °C ± 0.2
E540A ΔTm = 0.4 °C ± 0.2
K789A ΔTm = 4.4 °C ± 0.1
F561A ΔTm = 3.0 °C ± 0.1
D794E ΔTm = 0.9 °C ± 0.0
STX [nM]
0
19.5
625
5000
20000
Fluorescence (a.u)
Temperature (°C)
B
Tm (°C)
RcSxph
E540Q
Y558I
Y558A
D794N
D794A
D785A
D785N
Y558F
F561A
E540A
Q787A
Q787E
D794E
K789A
T563A
F784Y
P727A
I782A/Y558A
F784S
I782F
F784L
F784A
I782A
F784C
E540D
Y795A
C
Tm (°C)
ΔTm (°C)

**Figure S1 *Rc*Sxph thermofluor (TF) assay. A,** Exemplar thermofluor (TF) assay results for *Rc*Sxph in the presence of the indicated concentrations of STX. Curves for *Rc*Sxph, E540A, P727A, Y558A, F561A, and T563A are identical to those shown in Figs. 1A and 1B. ΔTm values are indicated. **B,** Baseline Tm values for *Rc*Sxph and the indicated mutants. **C,** Plot of Tm vs. ΔTm for the proteins in '**B**'. Colored boxes in 'A' and bars in '**B**' correspond to ΔΔG classifications in Table 1. Error bars are S.E.M.

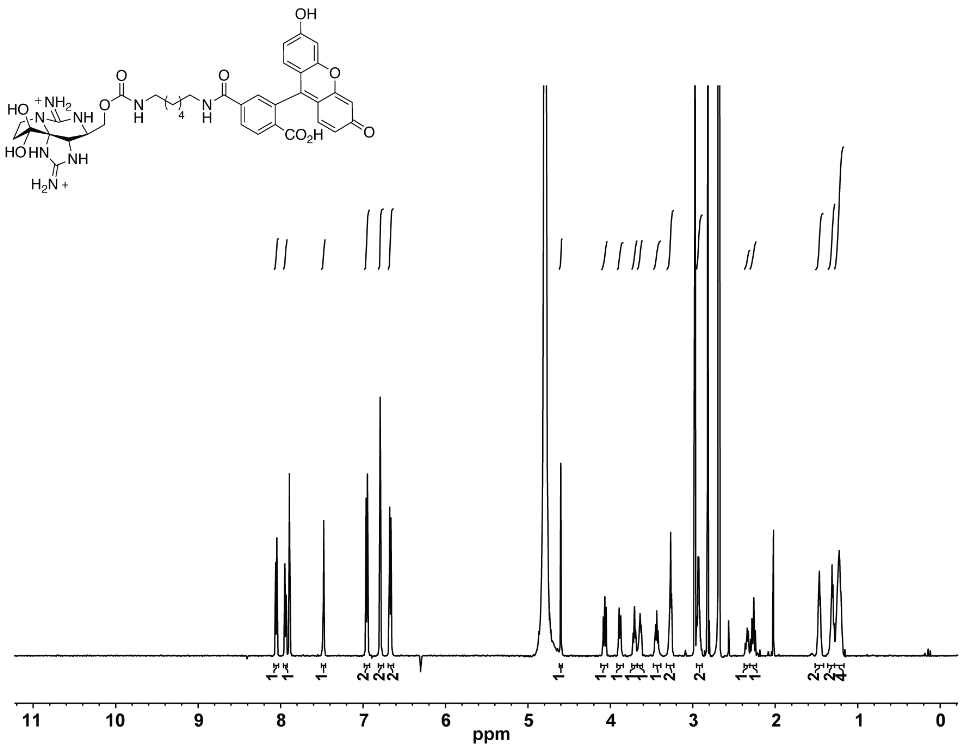

**Figure S2 F-STX NMR spectrum.** $^1$H NMR (600 MHz, $D_2O$) δ 8.05 (d, *J* = 8.1 Hz, 1H), 7.94 (d, *J* = 8.9 Hz, 1H), 7.48 (s, 1H), 6.95 (d, *J* = 9.0 Hz, 2H), 6.79 (s, 2H), 6.67 (dt, *J* = 9.1, 2.2 Hz, 2H), 4.60 (d, *J* = 1.2 Hz, 1H), 4.09–4.05 (m, 1H), 3.89 (dd, *J* = 11.6, 5.2 Hz, 1 H), 3.70 (dt, *J* = 10.1, 5.5 Hz, 1H), 3.64 (dd, *J* = 8.7, 5.4 Hz, 1H), 3.47–3.42 (m, 1H), 3.27 (t, *J* = 6.6 Hz, 2 H), 2.97–2.89 (m, 2H), 2.36–2.33 (m 1H), 2.30–2.24 (m, 1H), 1.48–1.45 (m, 2H), 1.32–1.29 (m, 2H), 1.25–1.21 (m, 4H) ppm.

**Figure S3** **Chen *et al.***

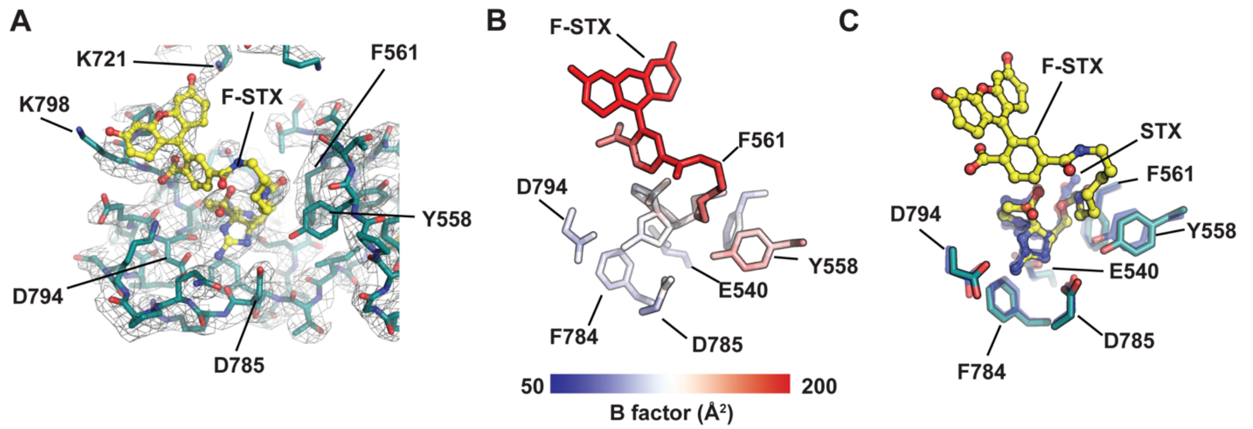


**Figure S3 Structure of the *Rc*Sxph:F-STX: complex. A,** Exemplar electron density (1σ) for *Rc*Sxph (deep teal) and F-STX (yellow). **B,** *Rc*Sxph:F-STX: B-factors for the F-STX ligand and select binding site residues. **C,** Superposition of the STX binding sites of the *Rc*Sxph:F-STX: and *Rc*Sxph:STX (PDB:6O0F) (blue) (*1*) complexes.

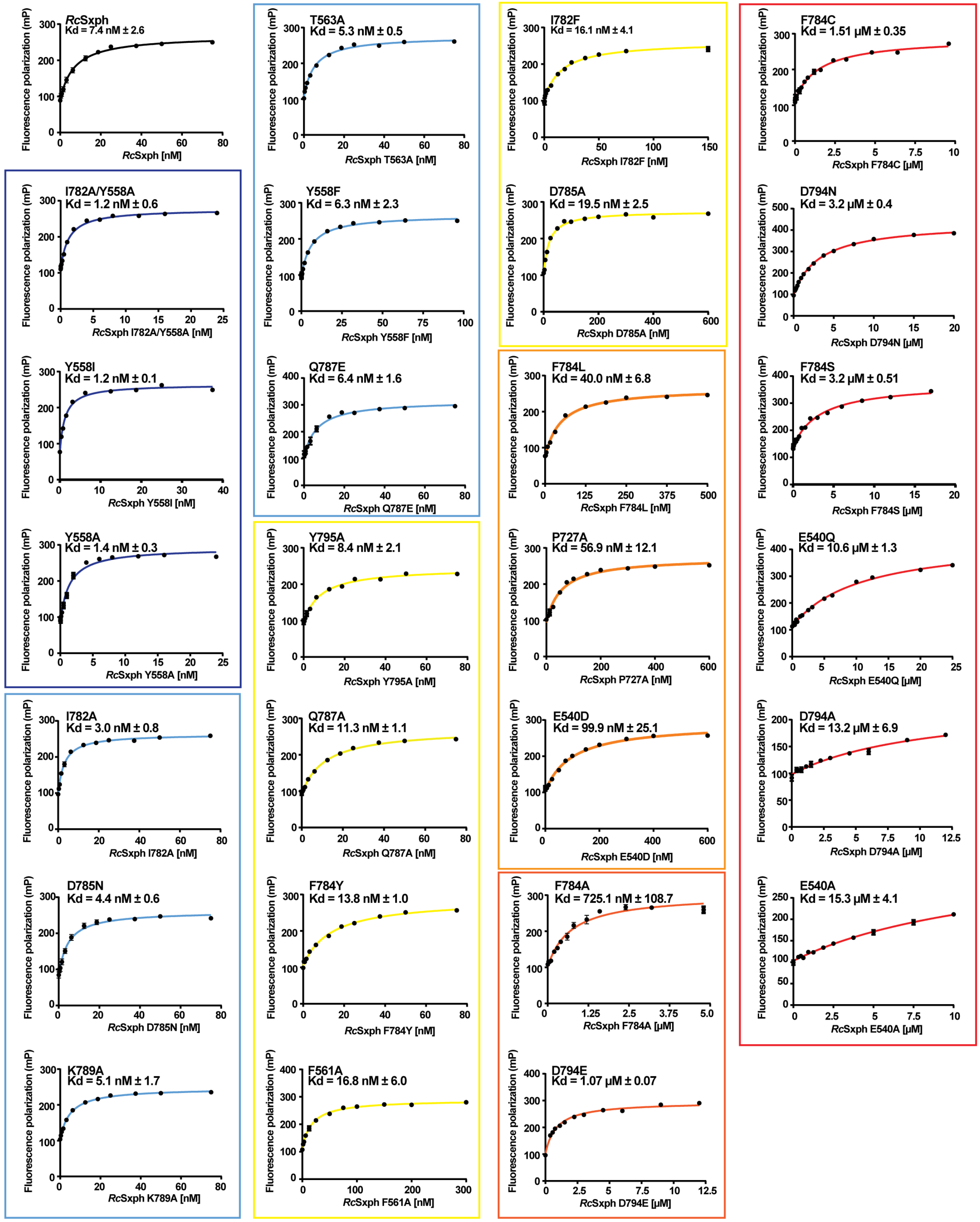


**Figure S4 *Rc*Sxph fluorescence polarization (FP) assay.** Exemplar FP binding curves and Kds for *Rc*Sxph and the indicated mutants. Curves for *Rc*Sxph, E540A, P727A, Y558A, F561A, and

T563A are identical to those shown in Fig. 1D. Colored boxes and lines in correspond to ΔΔG classifications in Table 1. Error bars are S.E.M.

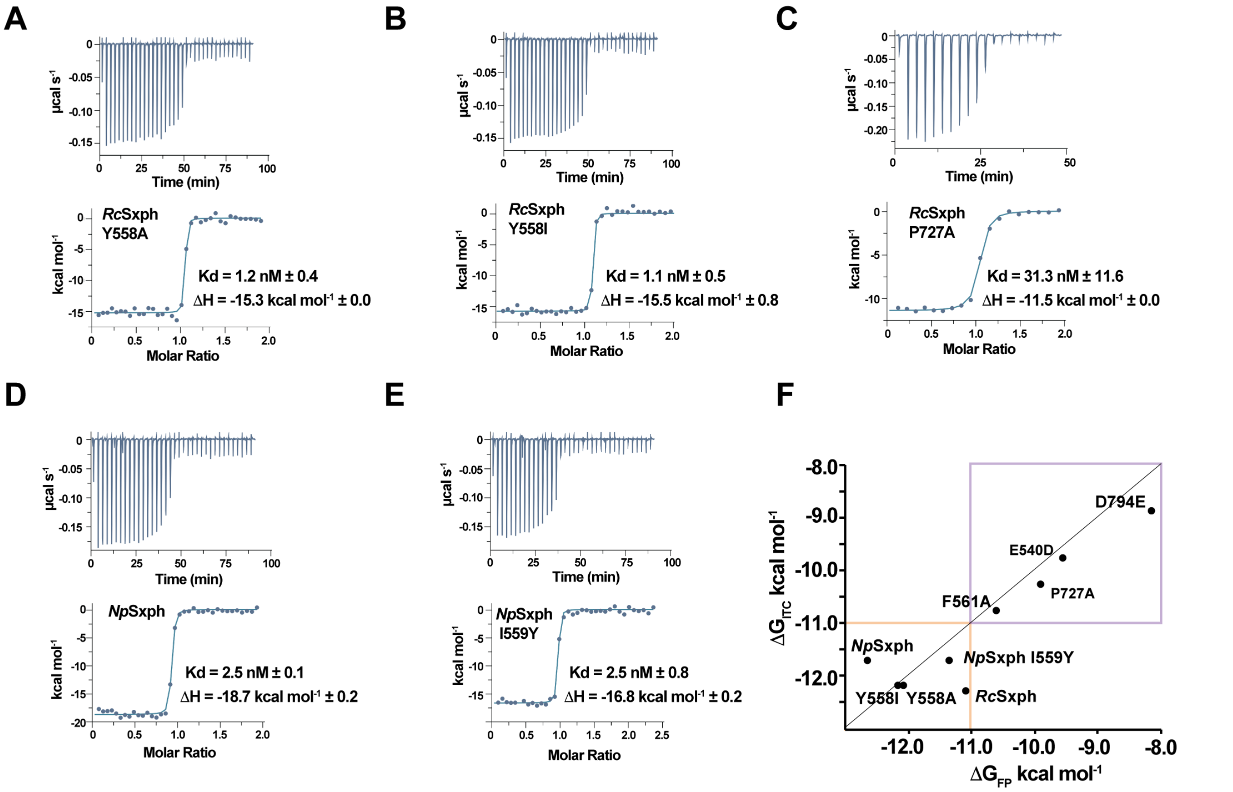


**Figure S5 *Rc*Sxph and *Np*Sxph Isothermal titration calorimetry.** Exemplar ITC isotherms for **A,** 100 µM STX into 10 µM *Rc*Sxph Y558A, **B,** 100 µM STX into 10 µM *Rc*Sxph Y558I, **C,** 100 µM STX into 10 µM *Rc*Sxph P727A, **D,** 100 µM STX into 9.7 µM *Np*Sxph, and **E,** 100 µM STX into 7.9 µM *Np*Sxph I559Y. **F,** Comparison of $\Delta G_{ITC}$ for STX and $\Delta G_{FP}$ for F-STX for *Rc*Sxph, *Np*Sxph, and indicated mutants. Purple box highlights region of good correlation. Orange box indicates region outside of the ITC dynamic range. *Rc*Sxph data are identical to Fig. 1G.

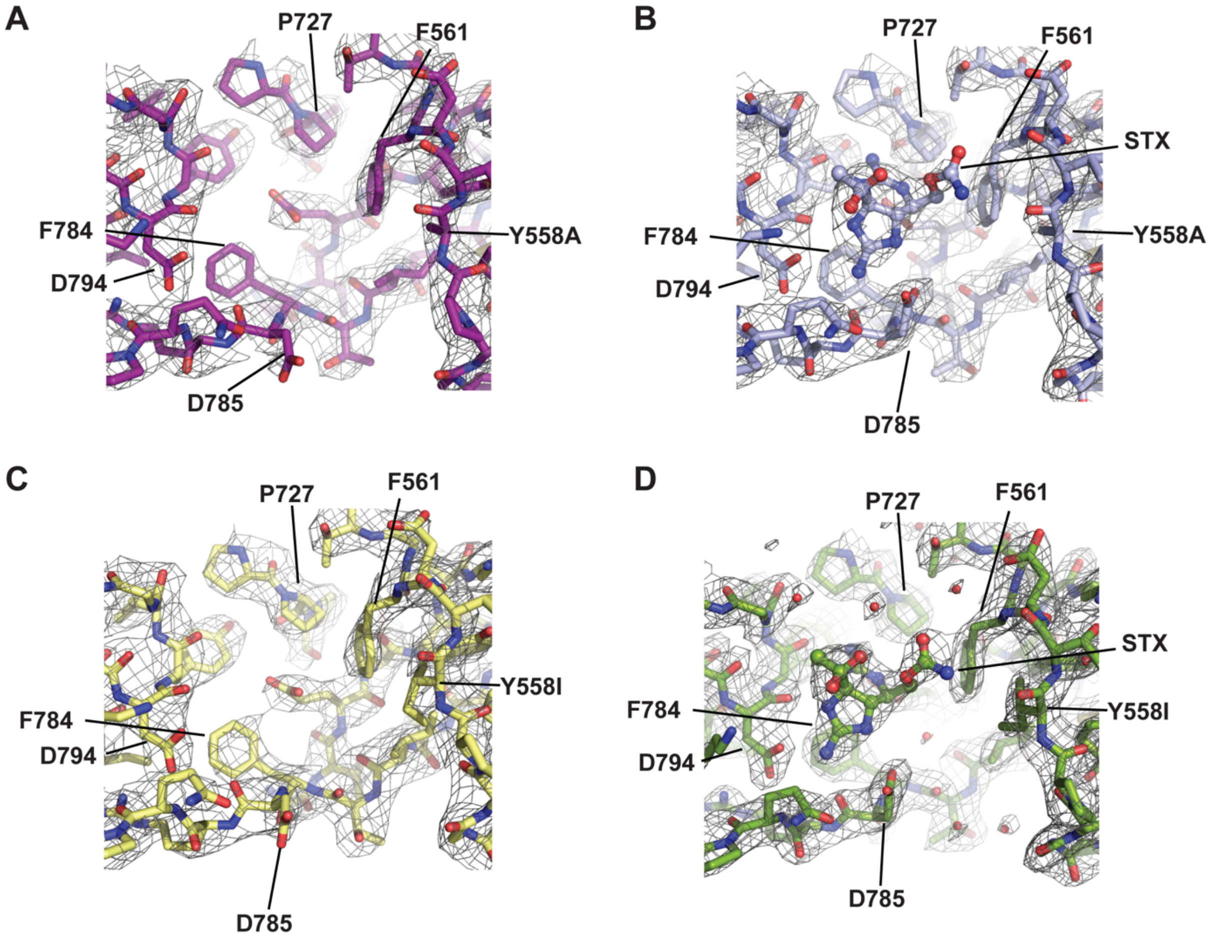


**Figure S6 *Rc*Sxph Y558A and *Rc*Sxph-Y558I structures and STX complexes.** Exemplar electron density (1.5 $\sigma$) for **A,** *Rc*Sxph Y558A (purple), **B,** *Rc*Sxph-Y558A:STX (light blue), **C,** *Rc*Sxph-Y558I (pale yellow), and **D,** *Rc*Sxph-Y558I:STX (splitpea). Select residues and STX are indicated.

8 June 2022

### Figure S7

Chen *et al.*

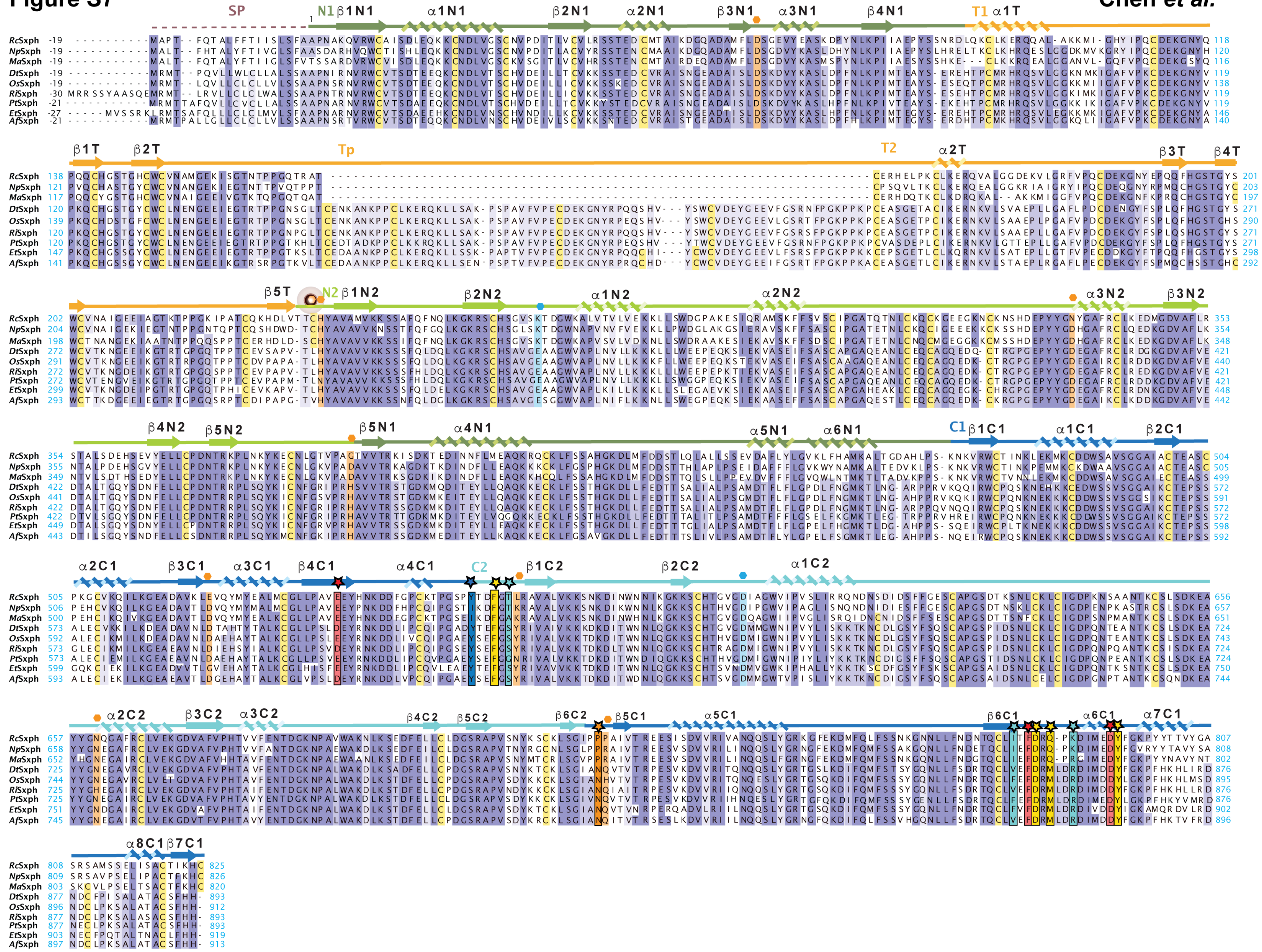

**Figure S7 Frog Sxph sequence alignment.** Sxph sequence alignment for *Rc*Sxph, *Np*Sxph, *Ma*Sxph, *Dt*Sxph, *Os*Sxph, *Ri*Sxph, *Pt*Sxph, *Et*Sxph, and *Af*Sxph. Domains and secondary structure are from *Rc*Sxph. N1 (dark green), N2 (light green), Thy1 domains (orange), C1 (marine), C2 (cyan). STX binding site residues are indicated by stars and colored based on the alanine scan results in Table 1. Residues corresponding to transferrin $Fe^{3+}$ and carbonate ligands are indicated by orange and blue hexagons, respectively and highlighted (*1, 2*).

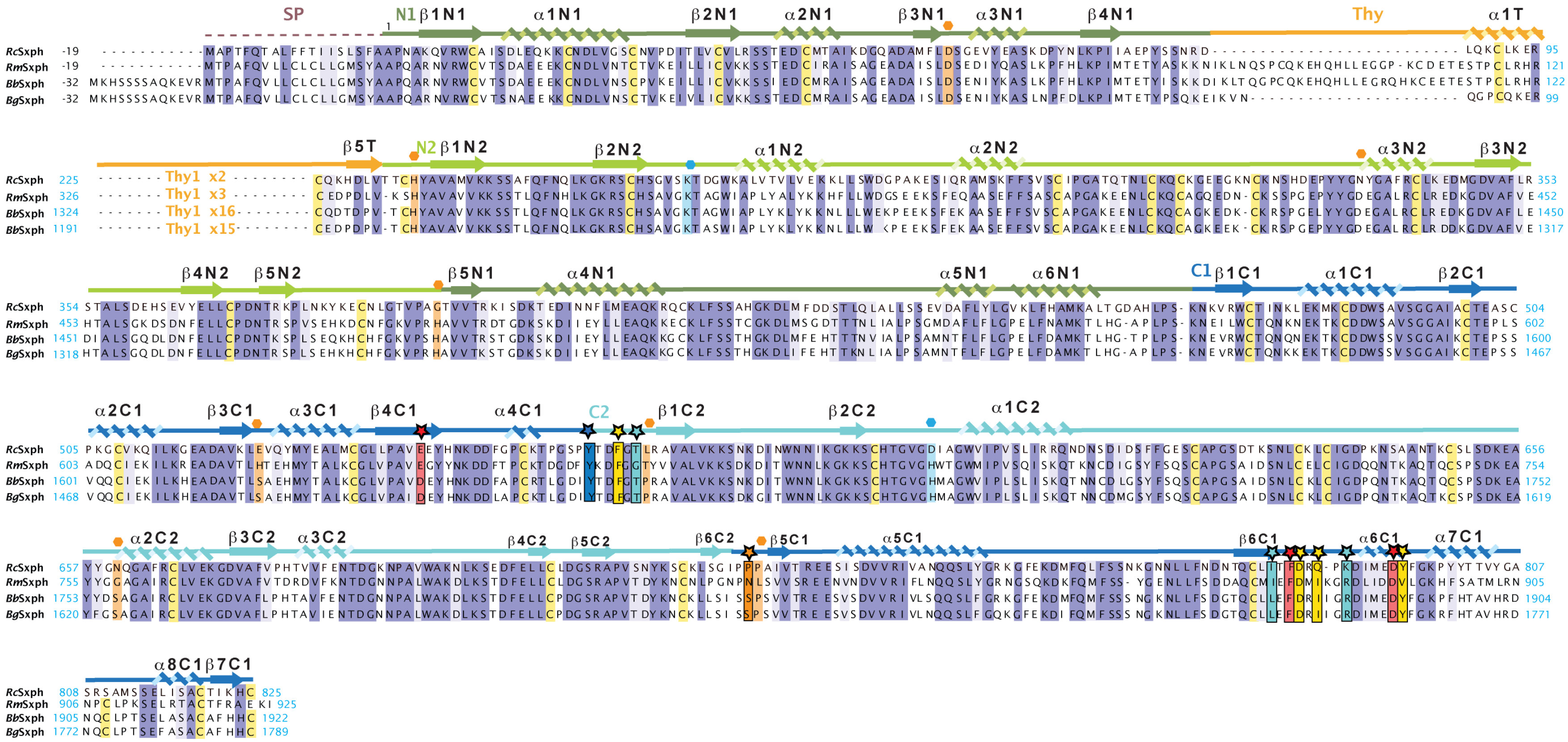

**Figure S8 Toad Sxph sequence alignment.** Sxph sequence alignment for *Rc*Sxph, and toad saxiphilins *Rm*Sxph, *Bb*Sxph (NCBI:XM_040427746.1), and *Bg*Sxph(NCBI:XP_044148290.1). Domains and secondary structure are from *Rc*Sxph. N1 (dark green), N2 (light green), Thy1 domains (orange), C1 (marine), C2 (cyan). STX binding site residues are indicated by stars and colored based on the alanine scan results in Table 1. Residues corresponding to transferrin $Fe^{3+}$ and carbonate ligands (*1, 2*) are indicated by orange and blue hexagons, respectively and highlighted. Only beginning and ends of the Thy1 domains are shown. Total number of Thy1 domains are indicated.

**Figure S9** **Chen *et al.***

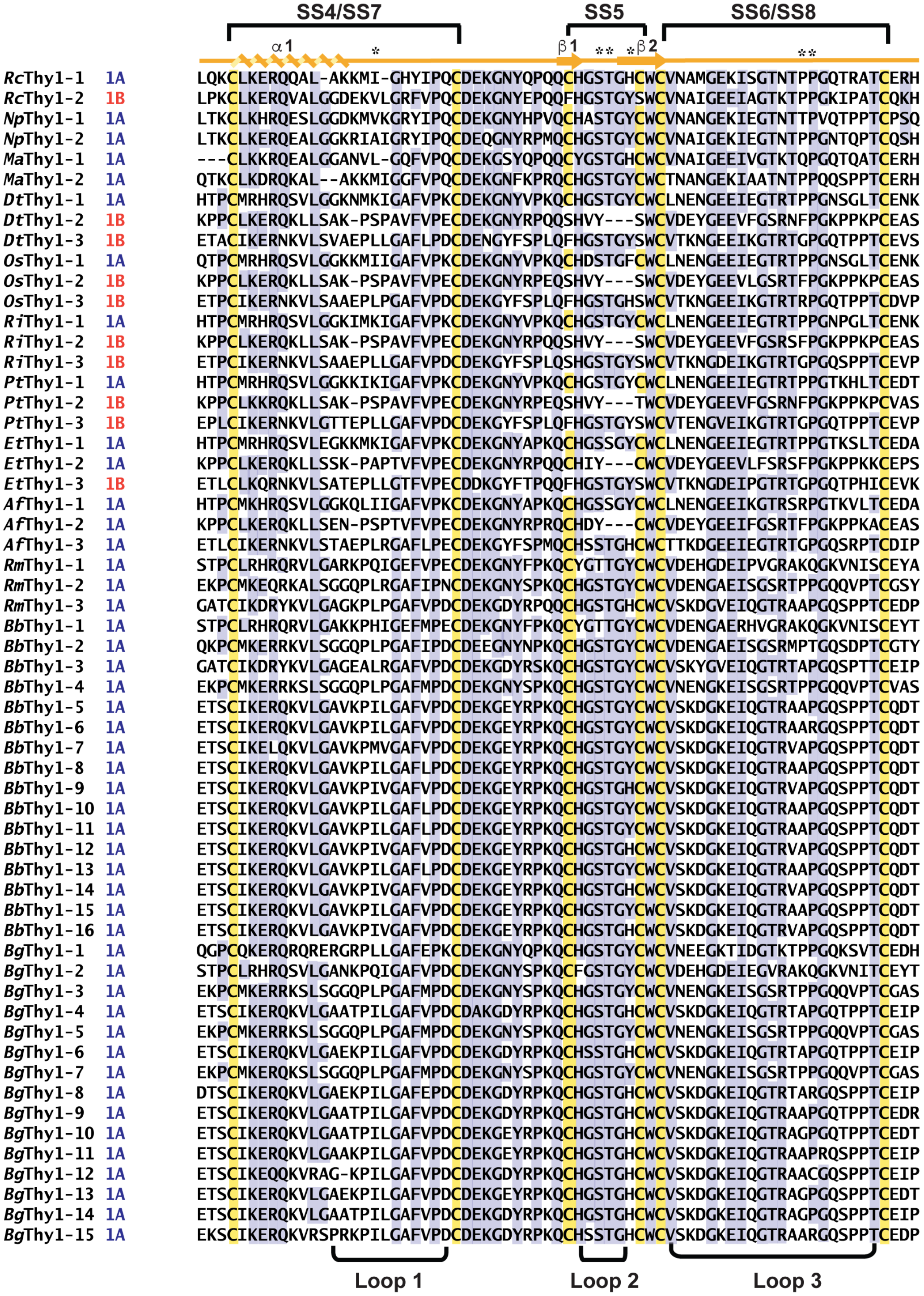


**Figure S9 Thy1 domain sequence alignment.** Thy1 domains from *Rc*Sxph, *Np*Sxph, *Ma*Sxph, *Dt*Sxph, *Os*Sxph, *Ri*Sxph, *Pt*Sxph, *Et*Sxph, *Af*Sxph, *Rm*Sxph, *Bb*Sxph, and *Bg*Sxph and the type (1A or 1B) are shown. Secondary structure from *Rc*Sxph Thy1-1 is shown. Cysteine are highlighted. SS4-SS8 indicate disulfide numbers from *Rc*Sxph. Loop regions are indicated.

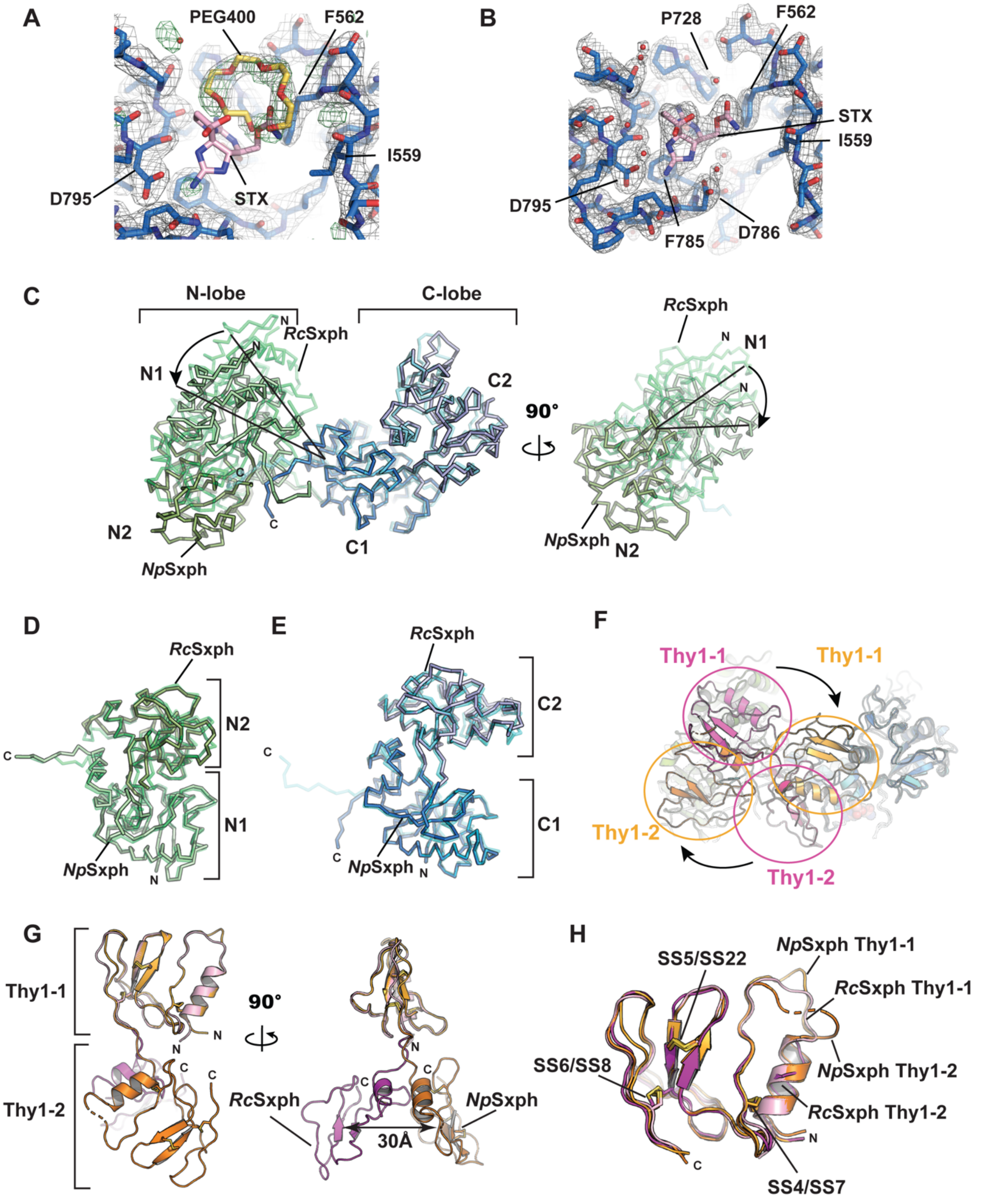


**Figure S10 *Np*Sxph structure and comparisons with *Rc*Sxph. A,** and **B,** Exemplar electron density for **A,** *Np*Sxph (2Fo-Fc, 1.5 σ, grey) and (Fo-Fc, 3.0 σ, green). **B,** *Np*Sxph:STX (2Fo-Fc, 1.5 σ, grey). *N*pSxph (marine), STX (pink), and PEG400 (yellow) are shown. STX (pink) from the

*Np*Sxph:STX complex is shown in ‘**A’** to compare with the PEG400 position**.** Select residues are labelled. **C,** *Np*Sxph and *Rc*Sxph superposition using the C-lobes. N- and C-lobes are green/light green and marine/light blue for *Np*Sxph and *Rc*Sxph, respectively. Arrow indicate relationships between *Np*Sxph and *Rc*Sxph N-lobes. **D,** Superposition of *Np*Sxph (green) and *Rc*Sxph (light green) N-lobes. **E,** Superposition of *Np*Sxph (marine) and *Rc*Sxph (light blue) C-lobes. **F**, Cartoon diagram of *Np*Sxph and *Rc*Sxph superposition from ‘**C**’ showing the change in Thy1 domain positions. *N*pSxph Thy1 domains (orange) and *Rc*Sxph Thy1 domains (magenta) are indicated. **G**, Cartoon diagam of *Np*Sxph and *Rc*Sxph Thy1 domains superposed on Thy1-1. *Np*Sxph and *Rc*Sxph Thy1-1 and Thy1-2 are light orange and pink and orange and magenta, respectively. **H,** Superposition of individual *Np*Sxph and *Rc*Sxph Thy1-1 and Thy1-2 domains. Colors are as in ‘**G**’. Disulfide bonds are indicated.

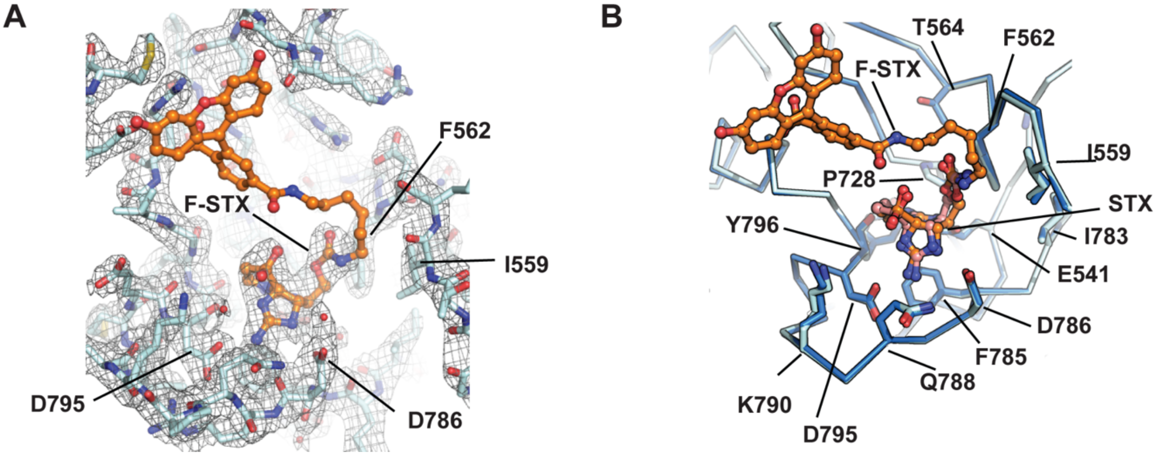


**Figure S11 Structure of the *Np*Sxph:F-STX complex. A,** Exemplar electron density for *Np*Sxph:F-STX(2Fo-Fc, 1.5 σ, grey). *Np*Sxph (cyan) and F-STX (orange). **B,** Comparison of *Np*Sxph:STX (marine) and *Np*Sxph:F-STX STX binding sites. STX from *Np*Sxph is pink. F-STX is orange. Select residues are indicated.

**Movie M1 *Rc*Sxph-Y558A conformational changes upon STX binding.** Morph between the apo-*Rc*Sxph-Y558A and *Rc*Sxph-Y558A:STX structures showing the STX binding pocket. Sidechains are shown as sticks. STX is red.

**Movie M2 *Rc*Sxph-Y558I conformational changes upon STX binding.** Morph between the apo-*Rc*Sxph-Y558I and *Rc*Sxph-Y558A:STX structures showing the STX binding pocket. Sidechains are shown as sticks. STX is red.

**Movie M3 Conformational changes between *R*cSxph and *Np*Sxph.** Morph between apo-*Rc*Sxph (PDB:6O0D) (*1*) (starting structure) and apo-*Np*Sxph (final structure). N-lobe (green), C-lobe (blue), and Thy domains (magenta) are shown. N1, N2, C1, and C2 subdomains and Thy1-1, and Thy1-2 are labeled.

**Movie M4 *Np*Sxph conformational changes upon STX binding.** Morph between the apo-*Np*Sxph and *Np*Sxph:STX structures showing the STX binding pocket. Sidechains are shown as sticks. STX is red.

| Table S1 Crystallographic data collection and refinement statistics | | | | |
|---|---|---|---|---|
| | ***Rc*Sxph -Y558A PDB:8D6P** | ***Rc*Sxph -Y558A:STX (co-crystal) PDB:8D6S** | ***Rc*Sxph -Y558I PDB:8D6Q** | ***Rc*Sxph -Y558I:STX (co-crystal) PDB:8D6T** |
| **Data Collection** | | | | |
| Space group | $P2_12_12_1$ | $P2_12_12_1$ | $P2_12_12_1$ | $P2_12_12_1$ |
| Cell dimensions a/b/c (Å) | 96.61, 109.05, 254.89 | 95.98, 107.14, 253.04 | 96.39, 107.15, 254.79 | 96.03, 107.81, 253.58 |
| α/β/γ (°) | 90, 90, 90 | 90, 90, 90 | 90, 90, 90 | 90, 90, 90 |
| Resolution (Å) | 47.81-2.60 (2.65-2.60) | 47.37-2.60 (2.65-2.60) | 47.61-2.70 (2.76-2.70) | 47.5-2.15 (2.19-2.15) |
| Rmerge (%) | 0.108 (4.094) | 0.115 (3.964) | 0.159 (4.833) | 0.089 (1.558) |
| I / σ I | 12.9 (0.9) | 12.5 (0.8) | 8.8 (0.6) | 16.2 (1.2) |
| CC(1/2) | 0.998 (0.532) | 0.998 (0.458) | 0.998 (0.419) | 0.999 (0.599) |
| Completeness (%) | 99.6 (100) | 99.9 (100) | 99.9 (99.8) | 99.5 (93.6) |
| Redundancy | 13.4 (13.9) | 13.4 (14.0) | 13.3 (14.0) | 12.2 (6.2) |
| Total reflections | 1116931 (62978) | 1085831 (61318) | 975607 (62994) | 1736282 (40652) |
| Unique reflections | 83173 (4517) | 81054 (4377) | 73319 (4504) | 142848 (6596) |
| Wilson B-factor | 83.42 | 84.83 | 90.34 | 44.01 |
| Wavelength (Å) | 1.033 | 1.033 | 1.033 | 1.033 |
| **Refinement** | | | | |
| $R_{work}$ / $R_{free}$ (%) | 22.79/26.37 | 23.20/26.17 | 23.42/27.21 | 20.73/23.37 |
| No. of chains in AU | 2 | 2 | 2 | 2 |
| No. of protein atoms | 12616 | 12616 | 12622 | 12622 |
| No. of ligand atoms | 0 | 42 | 0 | 72 |
| No. of water atoms | 77 | 60 | 79 | 794 |
| RMSD bond lengths (Å) | 0.002 | 0.002 | 0.003 | 0.003 |
| RMSD angles (°) | 0.50 | 0.49 | 0.55 | 0.59 |
| Ramachandran favored/allowed/outliers (%) | 94.90/4.91/0.18 | 94.29/5.47/0.25 | 93.61/6.08/0.31 | 95.33/4.30/0.37 |

| Table S1 Crystallographic data collection and refinement statistics (continued) | | | | |
|---|---|---|---|---|
| | *Rc*Sxph:F-STX (soaked) PDB:8D6U | *Np*Sxph PDB:8D6G | *Np*Sxph:STX (co-crystal) PDB:8D6M | *Np*Sxph:F-STX (soaked) PDB:8D6O |
| **Data Collection** | | | | |
| Space group | $P2_12_12_1$ | R3 | R3 | R3 |
| Cell dimensions a/b/c (Å) | 96.44, 109.37, 256.36 | 229.046, 229.046, 67.428 | 228.848, 228.848, 67.224 | 229.186, 229.186, 67.347 |
| α/β/γ (°) | 90, 90, 90 | 90, 90, 120 | 90, 90, 120 | 90, 90, 120 |
| Resolution (Å) | 47.97-2.65 (2.70-2.65) | 43.29-2.2 (2.279-2.2) | 42.55-2.0 (2.071-2.0) | 43.31-2.2 (2.279-2.2) |
| Rmerge (%) | 0.112 (4.136) | 0.05218 (0.8687) | 0.05456 (0.9961) | 0.07078 (1.889) |
| I / σ I | 11.1 (0.8) | 12.66 (0.89) | 14.11 (1.11) | 19.64 (1.35) |
| CC(1/2) | 0.999 (0.513) | 0.999 (0.465) | 0.998 (0.637) | 0.999 (0.602) |
| Completeness (%) | 99.9 (99.8) | 98.60 (91.37) | 99.89 (99.61) | 99.90 (99.97) |
| Redundancy | 10.4 (11.0) | 3.3 (2.1) | 5.2 (4.8) | 10.6 (11.0) |
| Total reflections | 830262 (49019) | 219806 (12669) | 459480 (42277) | 707304 (73200) |
| Unique reflections | 79524 (4469) | 66067 (6127) | 88641 (8856) | 66898 (6682) |
| Wilson B-factor | 83.73 | 52.73 | 49.89 | 55.11 |
| Wavelength (Å) | 1.033 | 1.033167 | 1.033167 | 1.033167 |
| **Refinement** | | | | |
| $R_{work}$ / $R_{free}$ (%) | 23.46/26.51 | 19.50/23.55 | 19.25/22.09 | 19.64/24.26 |
| No. of chains in AU | 2 | 1 | 1 | 1 |
| No. of protein atoms | 12630 | 6373 | 6385 | 6346 |
| No. of ligand atoms | 110 | 38 | 59 | 93 |
| No. of water atoms | 88 | 229 | 287 | 161 |
| RMSD bond lengths (Å) | 0.002 | 0.004 | 0.004 | 0.006 |
| RMSD angles (°) | 0.50 | 0.60 | 0.62 | 0.75 |
| Ramachandran favored/allowed/outliers (%) | 94.23/5.41/0.37 | 95.71/4.17/0.12 | 95.85/3.90/0.24 | 94.19/5.56/0.25 |

**Table S2 *Rc*Sxph:STX and *Np*Sxph:STX thermodynamic binding parameters**

| | | N (sites) | Kd (nM) | ΔH (kcal mol$^{-1}$) | ΔS (cal mol$^{-1}$ K$^{-1}$) | ΔG (kcal mol$^{-1}$) | n |
|---|---|---|---|---|---|---|---|
| ***Rc*Sxph** | ***WT*** | 1.02 ± 0.01 | 1.2 ± 0.8 | -16.1 ± 0.2 | -12.7 ± 0.9 | -12.3 ± 0.5 | 3 |
| | **Y558A** | 1.01 ± 0.01 | 1.2 ± 0.4 | -15.3 ± 0.0 | -11.1 ± 1.1 | -12.2 ± 0.2 | 2 |
| | **Y558I** | 1.05 ± 0.03 | 1.1 ± 0.5 | -15.5 ± 0.3 | -10.5 ± 0.6 | -12.2 ± 0.3 | 2 |
| | **F561A** | 1.07 ± 0.01 | 13.4 ± 1.4 | -12.7 ± 0.2 | -6.6 ± 0.9 | -10.8 ± 0.1 | 3 |
| | **P727A** | 0.97 ± 0.03 | 31.3 ± 11.6 | -11.5 ± 0.0 | -4.2 ± 0.9 | -10.3 ± 0.2 | 2 |
| | **E540D** | 0.98 ± 0.02 | 68.9 ± 10.7 | -16.3 ± 1.7 | -21.9 ± 5.7 | -9.8 ± 0.1 | 4 |
| | **D794E** | 0.99 ± 0.01 | 312.5 ± 2.9 | -11.8 ± 0.0 | -8.9 ± 0.1 | -8.9 ± 0.1 | 2 |
| ***Np*Sxph** | ***WT*** | 0.92 ± 0.02 | 2.5 ± 0.1 | -18.7 ± 0.2 | -23.2 ± 0.8 | -11.8 ± 0.1 | 2 |
| | **I559Y** | 0.94 ± 0.03 | 2.5 ± 0.8 | -16.8 ± 0.2 | -16.9 ± 1.1 | -11.8 ± 0.2 | 4 |
